# Multiplex single-cell analysis of serotonergic neuron function in planarians reveals widespread effects in diverse cell types

**DOI:** 10.1101/2024.02.28.581916

**Authors:** Elena Emili, Dianalí Rodríguez-Fernández, Alberto Pérez-Posada, Helena García-Castro, Jordi Solana

**Affiliations:** Department of Biological and Medical Sciences, Oxford Brookes University, Oxford, UK

## Abstract

Neurons function by interacting with each other and with other cell types, often exerting organism-wide regulation. Serotonergic neurons play a systemic role in processes such as appetite, sleep and motor control. Functional studies in the planarian *Schmidtea mediterranea* have shown that impairment of serotonergic neurons results in systemic effects. Studying neurons and the tissues they interact with is challenging using either bulk or single-cell analysis techniques. While bulk methods merge the information from all cell types, single-cell methods show promise in overcoming this limitation. However, current single-cell approaches encounter other challenges including stress of cell dissociation, high cost, multiplexing capacity, batch effects, replication and statistical analysis. Here we used ACME and SPLiT-seq to generate a multiplex single-cell analysis of serotonergic neuron function in planarians by inhibiting *pitx* and *lhx1/5-1*, two transcription factors expressed in them. We recovered single-cell transcriptomic profiles of 47,292 cells from knockdown and control animals, including biological and technical replicates. Our results show that epidermal, muscular and the recently described parenchymal cell types are affected the most by serotonergic neuron impairment. By computationally dissecting each cell type, we elucidated gene expression changes in each, including changes in epidermis cilia genes and myofiber genes in muscle. Interestingly, parenchymal cells downregulate genes involved in neurotransmitter recycling, suggesting a glial-like function of these recently described enigmatic cell types. Our results will allow disentangling the complexity of serotonergic neuron inhibition by studying the downstream effectors and the affected tissues, and offer new data on the function of parenchymal cells in planarians. Ultimately, our results pave the way for dissecting complex phenotypes through multiplex single-cell transcriptomics.

## Introduction

Living systems are complex: the knockdown of just one gene can trigger widespread effects throughout the organism. First direct effects might affect the cell type or tissue where the gene is expressed, eliminating or altering the function of the cells. These effects can later propagate to other tissues, creating an indirect cascade that can escalate to systemic effects. This is especially important for neuronal cell types as they exert control over the rest of the body.

For instance, the perturbation of serotonergic neurons triggers widespread effects in different organisms. The serotonergic system is a fundamental component of the nervous system in bilaterian animals [1], playing a crucial role in regulating a wide range of physiological processes and behaviours [2–4]. Serotonin is a neurotransmitter associated in humans with functions such as mood regulation, appetite, sleep, motor control, and cognitive processes. The study of serotonergic systems has predominantly focused on vertebrate models, but in recent years attention has also turned to invertebrates [5, 6].

Planarians are an ideal model to study neuronal cell types [7] and their function as they have stem cells that constantly differentiate into all cell types, including neurons [8–10]. Furthermore, they can also regenerate a complete head, including its central nervous system (CNS) in a matter of days [11, 12]. Finally, RNAi allows perturbing these processes in adults, allowing the study of their associated phenotypes [13, 14]. Planarians have a nervous system comprising a compartmentalised brain [15], two ventral nerve cords with interconnecting commissures, as well as several neuronal plexuses [7]. Planarian neurons have been characterised as dopaminergic, serotonergic, GABAergic, cholinergic, and octopaminergic in cellular studies [7, 12]. Single cell studies have also begun to characterise the diversity of planarian neuronal populations [16–20].

Planarian neuron specification and differentiation is dependent on transcription factors (TFs) such as *coe*, *sim*, *hesl-3*, *otxA*, *soxB1-2* and others [21–27]. In planarians, TFs *Smed-pitx* and *Smed-lhx1/5-1* (hereafter *pitx* and *lhx1/5-1* respectively) are expressed in a relatively small population of serotonergic *tph*+ neurons [24, 26], but their silencing induces systemic effects. These entail defects in locomotion, elongation of the body and behavioural defects [24, 26]. The cascade leading to these effects is unknown and difficult to study with conventional methods such as gene specific assays of *in situ* hybridisation, qPCR or immunohistochemistry, as these approaches require prior information about the effects to design the experiments. Transcriptome-wide approaches such as RNA-seq are devoid of this problem but preclude the distinction between direct and indirect effects in different cell types and tissues. Thus, although the role of serotonin in planarians has been studied [12, 24, 26], a comprehensive understanding of the molecular mechanisms of serotonin biology is lacking.

Single-cell analysis has emerged in the last decade as a powerful approach to query living systems with cell type resolution [28–30]. For instance, single-cell transcriptomics has been key to obtaining cell type atlases of diverse organisms based on their gene expression profiles [31]. However, the application of single-cell approaches to dynamic scenarios of gene knockdown or other functional studies is still challenged by aspects related to the stress of cell dissociation protocols, batch effects, integration algorithms, lack of technical and biological variation estimation and experimental cost [29, 30, 32–42]. Recently, we introduced ACME as a method that allows simultaneous cell dissociation and fixation, avoiding the stress associated with live dissociation protocols [43]. Multiplexing approaches such as SPLiT-seq [44], a combinatorial barcoding single-cell transcriptomics technique, allow performing single-cell transcriptomic experiments including different samples and replicates in the same experiment, minimising batch effects and avoiding the need for integration methods [44–46]. Collectively, ACME and SPLiT-seq are well suited for undertaking functional genomic approaches with single-cell resolution, opening a new avenue for using single-cell approaches to query complex phenotypes.

Here we used ACME and SPLiT-seq to study the effects of impaired serotonergic function in planarians. Our results show that knockdown of *pitx* and *lhx1/5-1* triggered effects on serotonergic *tph*+ neurons, which are likely a direct effect of TF knockdown. We also observed effects on cell types that do not express these TFs, such as the epidermis and muscle, which explain effects in locomotion and body elongation. Interestingly, we observed effects in parenchymal cells (also known as *cathepsin*+ cells), which have been only recently described at the molecular level [17, 19, 47]. These cells include a glial cell type in planarians [47, 48], but the function of the remaining cells in the group, as well as their association with glial cells, is still debated [49, 50]. We showed that another group of parenchymal cells, the *pgrn*+ cells, are the major cell type that responds to serotonergic gene knockdown, besides neurons. Differential gene expression and functional enrichment analyses indicated that *pgrn+* parenchymal cells modulate genes involved in neurotransmitter recycling in response to serotonergic neuron knockdown. This suggests that these cells play a role in neurotransmitter catabolism related to that of glial cells. Our results provide a comprehensive picture of the role of serotonin in several tissues, and unveil an unexpected function of newly described planarian parenchymal cell types. Ultimately, our results underscore the power of multiplex single-cell transcriptomics experiments and show that they will be key to decode complex biological phenotypes.

### High-resolution multiplex single-cell study of serotonergic neuron transcription factor knockdown in planarians

To determine the effect of *pitx* and *lhx1/5-1* on serotonergic function in planarians at single-cell resolution, we performed an RNAi experiment by feeding uninjured worms with dsRNA five times (see Methods). We then examined the animals 10 days after the last feeding for any changes in their phenotype (Figure 1A). After silencing *pitx*, all animals showed impaired gliding locomotion and whole body elongation (20/20, 100%) in contrast to *control*(*RNAi*) worms (Figure 1A). Likewise, all *lhx1/5-1*(*RNAi*) animals displayed locomotion and behavioural defects (20/20, 100%). These comprise an inability of their tails to detach from the surface of the petri dish, and their head region being stretched laterally and tilting up more frequently compared to the *control*(*RNAi*) group (Figure 1A).

**Figure 1:**
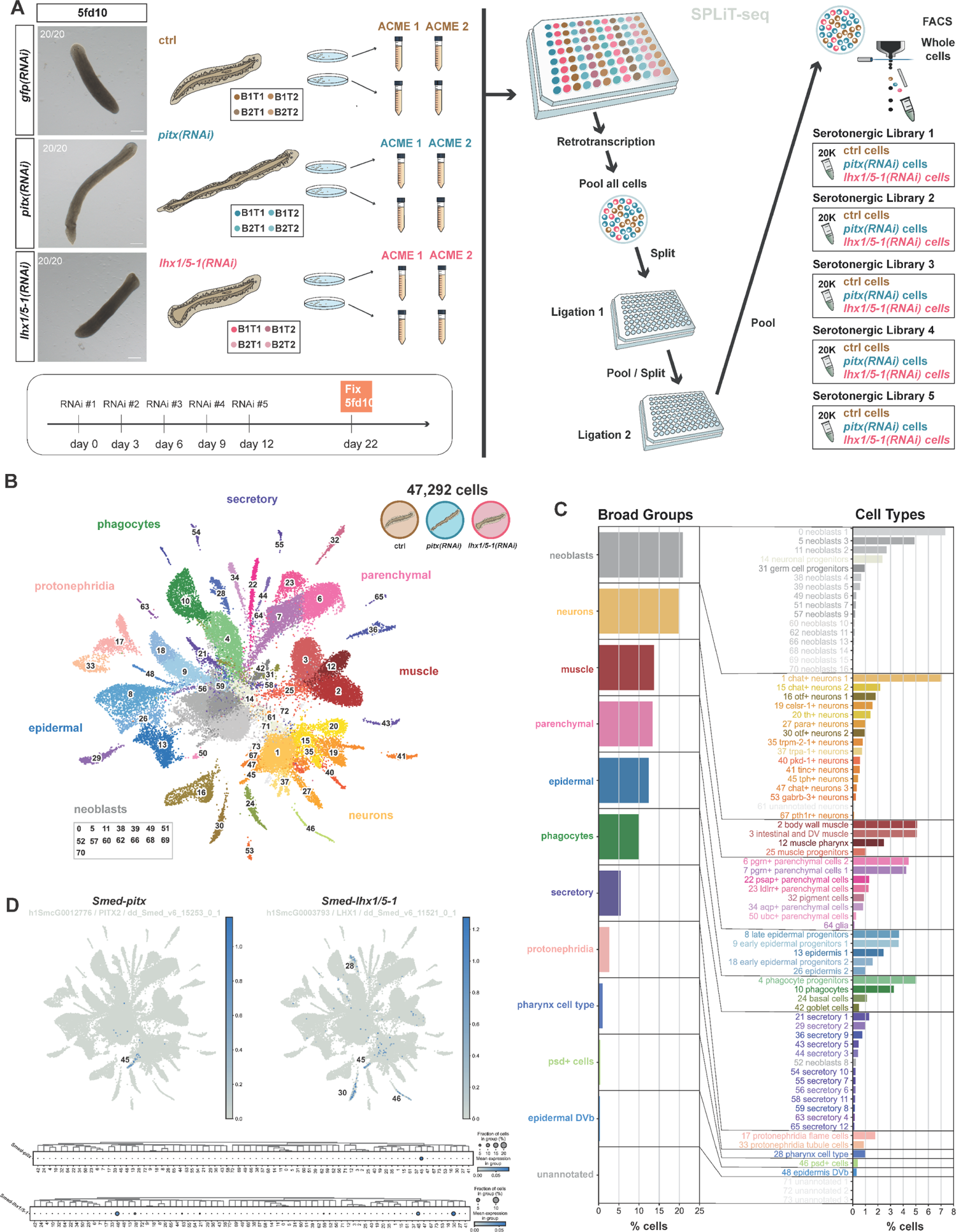
Functional single-cell study of serotonergic neurons. **A**: RNAi phenotypes and experimental design. Animals were fed 5 times and fixed 10 days after the last feeding (5fd10). B1-B2: biological replicate 1-2, T1-T2: technical replicate 1-2. **B**: UMAP visualisation of the high-resolution multiplex cell type study of serotonergic neuron transcription factor knockdown in planarians with clusters coloured according to their cell cluster classification. **C**: Annotated broad groups and cell types, and their frequencies (percentages). **D**: UMAP features plots and dotplots showing the expression of *pitx* and *lhx1/5-1*.

We then generated a multi-sample experiment by performing RNAi in biological replicates, with the animals kept and fed in independent containers, by different researchers. Each of these biological replicates was then subdivided in two technical replicates during the dissociation process. We obtained cell suspensions using ACME [43, 51] and processed them for scRNA-seq combinatorial barcoding using a modified version of SPLiT-seq (Figure 1A). The major variations were the use of a 96×96×96×5 combinatorial strategy (Supplementary File 1), the substitution of the oligo-dT(15)VN of the original SPLiT-seq protocol for an oligo-dT(30) [46, 52], and the inclusion of a FACS sorting step after the 3^rd^ round of combinatorial barcoding, as previously described [16] (see Methods). We multiplexed all samples and replicates in one single experiment and obtained five sub-libraries each one containing 20k cells from all samples, as counted by FACS sorting (Figure 1A). After sequencing and computational analysis, we obtained a dataset of 47,292 total cells, including *pitx*(*RNAi*) and *lhx*(*RNAi*) knockdowns and *control*(*RNAi*) samples of all biological and technical replicates (Figure 1B, Supplementary Figure 1A-B). With this experimental workflow the use of integration algorithms was not needed. Leiden clustering at resolution 3 resolved 74 cell populations, largely corresponding to planarian well known cell types (Figure 1B, Supplementary Figure 1C). We grouped these cell type identities in broad groups by hierarchical clustering (Figure 1C, Supplementary Figure 1D). We annotated cell clusters using previous literature and cross referencing published markers with markers obtained from this experiment (Figure 1B-C, Supplementary Figure 1C, Supplementary File 2-4).

As in previous experiments, SPLiT-seq resulted in a low UMI and gene per cell content (Supplementary Figure 1A-B), but allowed us to distinguish a high number of validated cell clusters that recapitulate *Schmidtea* cell type diversity. The average mean counts and genes per cluster varied from cluster to cluster (Supplementary Figure 1E) and the number of genes detected in each cell largely correlated with the number of counts obtained.

Our dataset resolved 16 neuronal cell populations, including aminergic, cholinergic, glutamatergic, glycinergic, GABAergic, sensory ciliated neurons, neuronal progenitors and clusters containing relatively rare cell types such as the glia and the photoreceptors (Figure 1C, Supplementary File 5). Each individual neuron has a distinctive gene-expression signature that shapes its specific form and function. This signature results from the combinatorial expression of enzymes, receptors, and structural proteins, which collectively define the neuron’s unique neural identity. For serotonergic neurons, tryptophan hydroxylase (*tph*) is the rate limiting enzyme that catalyses the first step in the synthesis of 5-hydroxytryptamine, also known as serotonin [2–4]. By comparing the expression patterns of this gene and other established marker genes, such as the serotonergic transporter *sert*, we were able to assign the serotonergic neuron cell type identity to cluster 45 (*tph*+ neurons). Other neuronal cell types were assigned identities by assessing their gene markers in a similar manner.

Previous *in situ* hybridization data showed that *pitx* and *lhx1/5-1* are co-expressed in ventrally located serotonergic neurons [24, 26]. In addition, a strong *lhx1/5-1* expression has also been previously observed in the pharynx by in situ hybridization and in the *otf*+ neurons in previously published single-cell data [17, 19, 53]. In agreement with these previous reports, our dataset showed expression of both *pitx* and *lhx1/5-1* in cluster 45 (*tph*+ neurons) (Figure 1D). Additionally, *lhx1/5-1* was also expressed in clusters 28 (pharynx cell type), 30 (*otf*+ neurons 2) and 46 (*psd*+ cells) (Figure 1D).

Collectively, these results show that our multiplex single-cell study of serotonergic neuron knockdown is a good quality dataset that resolves well-known planarian cell types in 74 cell clusters that are well integrated between RNAi samples. This includes up to 16 distinct neuronal cell populations, including a *tph*+ serotonergic neuron cell population that expresses both *pitx* and *lhx1/5-1*.

### Dynamic cellular changes in response to serotonergic loss-of-function

Previous experiments have shown the loss of serotonergic neurons upon RNAi knockdown of either *pitx* or *lhx1/5-1*, as well as the loss of a small GABAergic neuronal subpopulation after *lhx1/5-1* knockdown [24, 26, 53]. To explore changes in cell type abundances, we initially aimed to determine if the two experimental groups had the same cell type composition compared to the control group. Our multiplexed analysis allows detecting all cell clusters in both RNAi samples and the control (Figure 2A). The UMAPs of the three individual conditions displayed a similar distribution of cell types with no populations being restricted to a specific treatment. We then explored cell percentages distributions per clusters at the annotated Leiden resolution 3, confirming that all clusters have cells from all experimental groups. Overall, this analysis shows that, after silencing either *pitx* or *lhx1/5-1*, the animals contain the same essential cell clusters, and that no major batch effects are observed even in the absence of batch correction algorithms. Careful observation of the cell percentages for each cluster suggested differences in cell type abundances, specifically concerning the serotonergic cluster among others (Figure 2A). To analyse which clusters differ in cell proportions, we analysed cell type frequencies using a Fisher’s Exact Test (Supplementary Figure 2). This analysis revealed that, in *pitx*(*RNAi*) samples, cluster 13 (epidermis 1), cluster 31 (germ cell progenitors) and cluster 45 (*tph*+ neurons) were significantly depleted compared to *control*(*RNAi*) samples. In contrast, cluster 7 (*pgrn*+ parenchymal cells 1) and cluster 22 (*psap*+ parenchymal cells) were significantly enriched (Figure 2B, D). In *lhx1/5-1*(*RNAi*) samples, on the other hand, cluster 13 (epidermis 1), cluster 31 (germ cell progenitors), cluster 10 (phagocytes), cluster 46 (psd+ cells) and cluster 21 (secretory 1) were significantly depleted compared to *control*(*RNAi*) samples. In contrast, cluster 34 (*aqp*+ parenchymal cells), cluster 32 (pigment cells), cluster 6 (*pgrn*+ parenchymal cells 2), and cluster 7 (*pgrn*+ parenchymal cells 1) were significantly enriched (Figure 2C, E).

**Figure 2:**
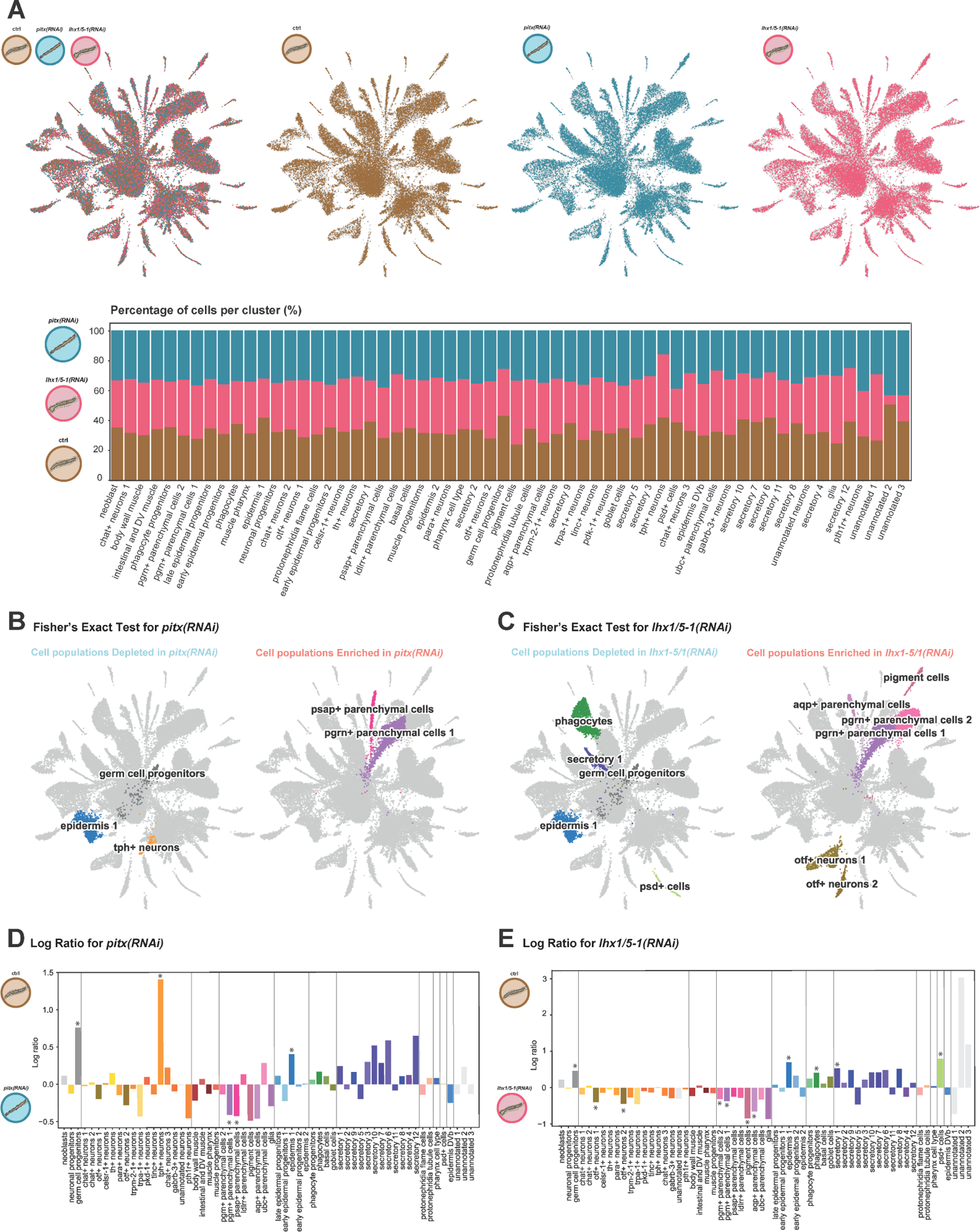
Differences in cell proportions after *pitx* and *lhx1/5-1* silencing. **A**: UMAP representation of *control*(*RNAi*), *pitx*(*RNAi*) and *lhx1/5-1*(*RNAi*) cells merged and separated by experimental groups (upper). Stacked bar plot of cell percentages by experimental conditions based on clustering annotation (lower). **B**: UMAP representation of the results of the Fisher’s Exact Test showing depleted and enriched cell populations in *pitx*(*RNAi*) compared to *control*(*RNAi*) animals. **C**: UMAP representation of the results of the Fisher’s Exact Test showing depleted and enriched cell populations in *lhx1/5-1*(*RNAi*) compared to *control*(*RNAi*) animals. **D**: Bar plot of percentage log2 ratios of cell clusters in *pitx*(*RNAi*) vs. *control*(*RNAi*) planarians. **E**: Bar plot of percentage log2 ratios of cell clusters in *lhx1/5-1*(*RNAi*) vs. *control*(*RNAi*) planarians. Significance levels are based on p-values < 0.01.

Interestingly, *tph*+ neurons are depleted significantly in *pitx*(*RNAi*) but not in *lhx1/5-1*(*RNAi*). Previous immunohistochemistry and *in situ* hybridization studies have shown the loss of a population of serotonergic *tph*+ neurons scattered along the planarian body after silencing *lhx1/5-1*, while *tph*+ expressing cells that locate around the pharynx appeared unaffected [24, 26, 53]. This likely explains the lack of *tph*+ cell depletion after *lhx1/5-1* knockdown.

### Multiplex differential gene expression analysis by cluster reveals major serotonin-responsive cell types

Planarian serotonergic *tph+* neurons are a relatively unabundant (0.41%) cell population. However, knockdown of *pitx* and *lhx1/5-1* triggers widespread effects with impairment in neuronal development, physiology, and overall animal behaviour. To understand complex phenotypes such as this, it is important to disentangle the direct effects that happen in the serotonergic neurons from the indirect downstream effects that may happen in different cell types that respond to serotonin. Traditional bulk RNA-seq measures the average gene expression levels across the entire cell population, providing an overall snapshot of the transcriptome (Figure 3A), and therefore it has limitations in distinguishing between direct, indirect, and compensatory effects in loss-of-function experiments. To overcome these limitations, single-cell RNA-seq using ACME and SPLiT-seq provides information about gene expression at the single-cell level, making it possible to identify cell-specific responses and changes with cell type resolution, by aggregating counts from each cell cluster in pseudobulk samples (Figure 3A). Our multiplexed approach contains biological and technical replicates which are key in bulk RNA-seq to detect significantly regulated genes. Several authors have pointed out that the need for replicates also applies to single-cell pseudobulk approaches [35], but this remains challenging due to its high cost. Single-cell combinatorial barcoding approaches such as SPLiT-seq are therefore an obvious solution as they can multiplex several samples in one experiment, reducing cost and batch effects.

**Figure 3:**
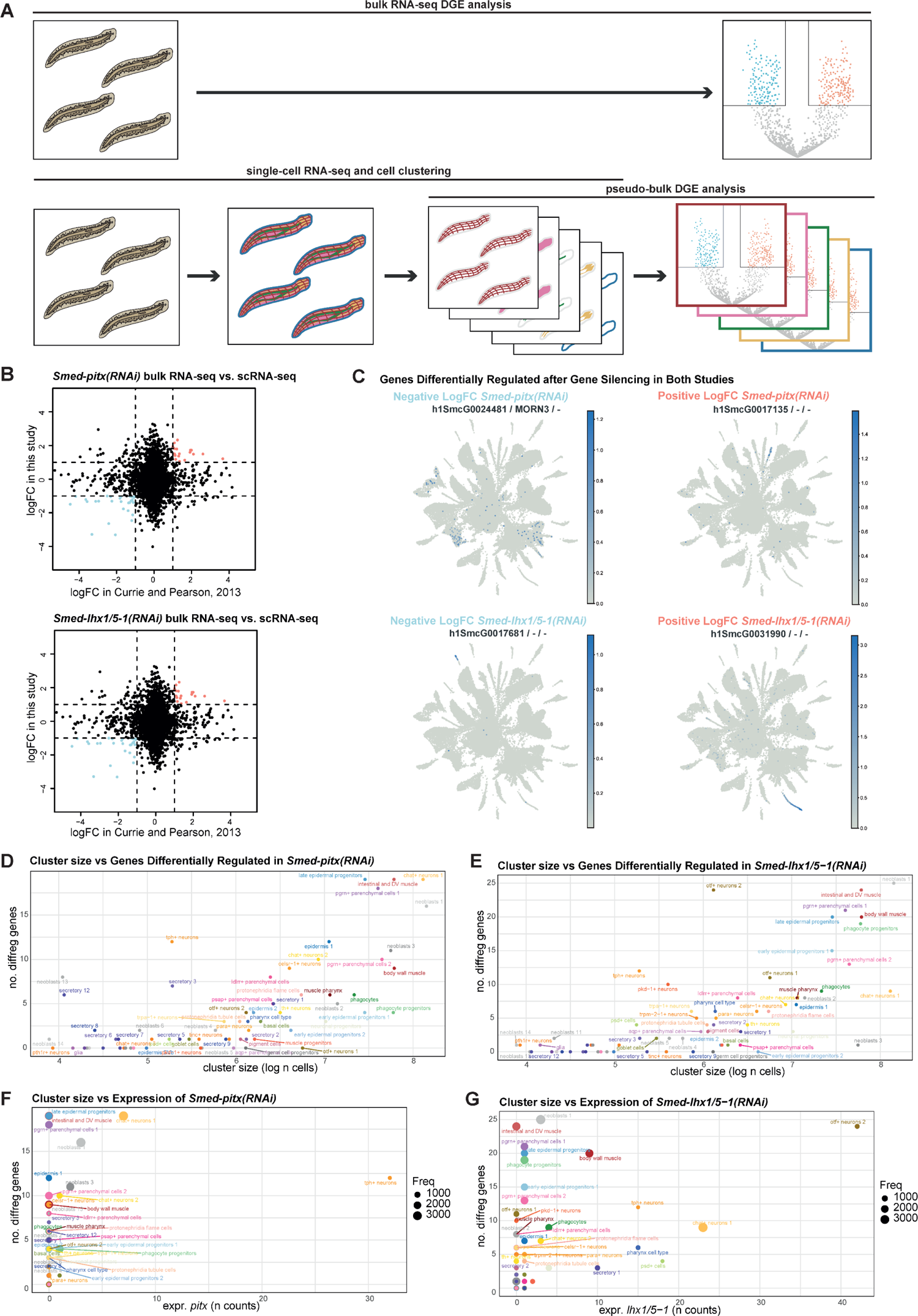
Differential gene expression analysis with DEseq2 in bulk RNA-seq and single-cell RNA-seq. **A**: Cartoon representation of DGE analysis in bulk RNA-seq and single-cell cluster pseudo-bulk. **B**: Correlation between results of DESeq2 in bulk RNA-seq (Currie and Pearson, 2013) vs. whole pseudo-bulk single-cell RNA-seq of this study. **C**: Top differentially regulated genes present in both analyses expressed in non-neuronal cell types. **D**: Scatter plot showing the number of differentially regulated genes per cluster compared to the log-transformed cell count within each cluster for *pitx*(*RNAi*) DGE analysis. **E**: Scatter plot showing the number of differentially regulated genes per cluster compared to the log-transformed cell count within each cluster for *lhx1/5-1*(*RNAi*) DGE analysis. **F**: Scatter plot showing the number of differentially regulated genes per cluster compared to the expression of *pitx* within each cluster for *pitx*(*RNAi*) DGE analysis. **G**: Scatter plot showing the number of differentially regulated genes per cluster compared to the expression of *lhx1/5-1* within each cluster for *lhx1/5-1*(*RNAi*) DGE analysis.

Previous studies conducted bulk RNA-seq in *pitx*(*RNAi*) and *lhx1/5-1*(*RNAi*) intact animals [24] and *pitx*(*RNAi*) only in regenerating animals [26]. We reanalysed the bulk RNA-seq dataset of *pitx*(*RNAi*) and *lhx1/5-1*(*RNAi*) conducted in intact animals performing DGE analysis with DESeq2 [54] (Supplementary Figure 3A-C, Supplementary File 6) and compared it with the single-cell dataset of our study analysed in pseudo-bulk with cells from all clusters together (Figure 3B). The correlation between these two studies was not robust given the variations in sequencing platforms, and differences in methodologies (Figure 3B). Despite these disparities, there was concordance in the up- and down-regulation of certain key genes, such as *sert* (Figure 3B). Inspection of the top genes that exhibit differential regulation in both studies, showed that some of these genes are expressed in locations beyond neuronal cell types (Figure 3C) and are therefore likely downstream, indirect effects of serotonergic neurons. Interestingly, some of these genes were expressed in clusters that are affected in cell number in our previous analysis, such as cluster 13 (epidermis 1) for *pitx*(*RNAi*) and cluster 46 (*psd*+ cells) for *lhx1/5-1*(*RNAi*) (Figure 3C). Altogether, this analysis showcases a moderate level of concordance between previous bulk RNA-seq experiments and the pooled pseudobulk of our single cell experiment, and that the major differentially regulated genes are not all expressed in serotonergic neurons, confirming the merging of all effects at bulk or pseudobulk level.

We then aimed at increasing the resolution of the previous analysis by considering each cell cluster independently (Figure 3A). We first determined which cell types change their gene expression patterns in response to *pitx* and *lhx1/5-1* gene knockdown. Taking full advantage of our multiplexed single-cell approach, we aggregated gene expression counts informed by cluster identity and sample of origin. This way we generated pseudo-bulk count tables for each cell type (Figure 3D-G, Supplementary Figure 3D-E) and broad group (Supplementary Figure 4) separately, computationally dissecting each cell type in each sample. We then used DESeq2 to perform differential gene expression (DGE) analysis including biological and technical replicates in each cell type. To assess the rate of response to the knockdown on each cell type, we examined the relationship between cluster size and the number of differential genes detected in each cell type (Figure 3D-E, Supplementary Figure 3D-E) and broad group (Supplementary Figure 4). The number of differentially regulated genes rises proportionally with cluster size, as it correlates with the volume of reads included in the analysis (Figure 3D-E). This pattern persists when examining broader group categories (Supplementary Figure 4). However, some exceptions lie outside of this distribution, with a much higher number of differentially regulated genes compared to cluster size (Figure 3D-E, Supplementary Figure 3D-E, Supplementary Figure 4). To model these effects, we performed linear regression and considered the last quantile as the cell types that respond more dynamically to the gene knockdown. These included cluster 45 (*tph*+ neurons), cluster 8 (late epidermal progenitors), cluster 3 (intestinal and DV muscle), and cluster 7 (*pgrn*+ parenchymal cells 1) for both *pitx*(*RNAi*) and *lhx1/5-1*(*RNAi*) animals compared to the control (Figure 3D-E), consistent with previous results on cell abundances (Figure 2B-E). Cluster 30 (*otf*+ neurons 2) was significant only in the *lhx1/5-1*(*RNAi*) condition (Fig. 3E), consistent with its high expression of *lhx1/5-1* but not of *pitx*. This same analysis performed at broad group resolution consistently shows that neurons are the broad group that respond more dynamically to both *pitx* and *lhx1/5-1* knockdown at the gene expression level, while the epidermal group responds specifically to ther knockdown of *lhx1/5-1* (Supplementary Figure 4).

To differentiate potential direct from indirect effects we explored the relationship between the numbers of differentially expressed genes per cluster with the expression of the two transcription factors in each cluster (Figure 3F-G). This analysis revealed that changes in the gene expression programme of cluster 45 (*tph*+ neurons) were likely directly linked with the knockdown, while the changes of gene expression in cluster 8 (late epidermal progenitors), cluster 3 (intestinal and DV muscle), and cluster 7 (*pgrn*+ parenchymal cells 1) must be due to secondary, indirect or compensatory, effects that lead to the complexity of this phenotype, as *pitx* was not expressed in those cell types (Figure 3F). Similarly, DGE analysis in *lhx1/5-1*(*RNAi*) samples revealed a strong and likely direct effect in cluster 30 (*otf*+ neurons 2, Figure 3G). This was recapitulated at broad group resolution, especially for *lhx1/5-1*(*RNAi*) where changes in the neuron broad type reflected a direct effect of gene expression loss and the epidermal broad type an indirect effect (Supplementary Figure 4). Altogether, this analysis shows that single-cell RNA-seq using biological and technical replicates can detect gene expression changes at the cell type level, disentangling possible direct vs indirect effects.

### Effects of serotonergic transcription factor gene knockdown on *tph*+ neurons

To estimate the direct effects of the gene knockdown, we looked at the cell types that express *pitx* and *lhx1/5-1* [24] (Figure 1D). We focused on the broad neuronal type and two specific neuronal clusters, cluster 45 (*tph*+ neurons) and 30 (*otf*+ neurons 2), as they show the highest changes in gene expression between the knockdowns and the control. We visualised the genes that are differentially regulated in *tph*+ neurons and *otf*+ neurons 2 with volcano plots and heatmaps, and analysed the Gene Ontology (GO) term enrichment of up- and down-regulated genes (Figure 4A-C, Supplementary Figure 4, Supplementary Files 7-8).

**Figure 4:**
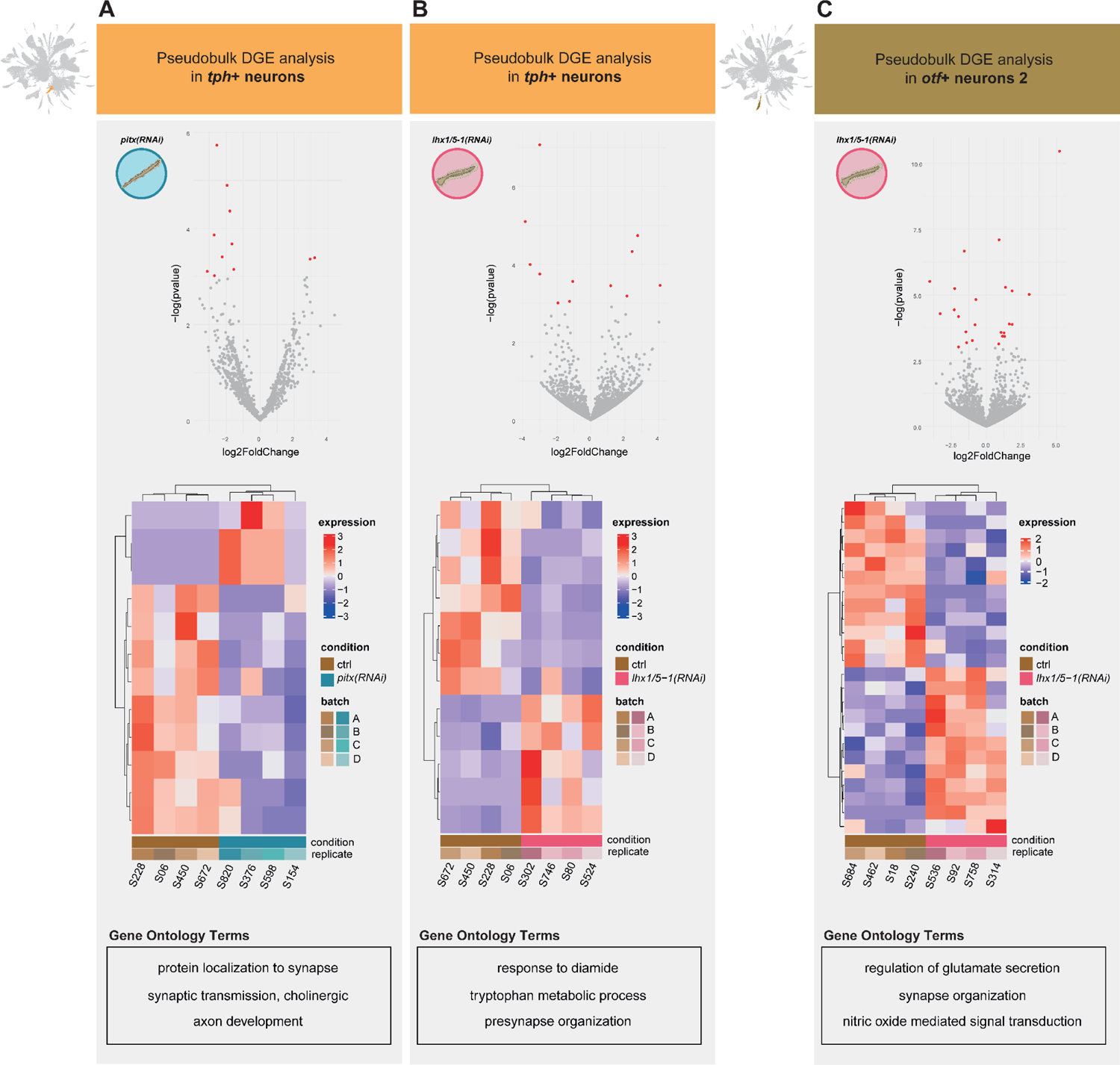
Knockdown effects on serotonergic cells. **A-C**: Volcano plot representation, heatmap representation and GO summary of the DGE analysis in *tph*+ neurons of *pitx*(*RNAi*) animals (A), *tph*+ neurons of *lhx1/5-1*(*RNAi*) (B) and *otf*+ neurons 2 of *lhx1/5-1*(*RNAi*) animals. Significance levels are based on p-values < 0.05.

Genes differentially expressed in *tph*+ neurons of *pitx(RNAi)* animals were predominantly dowregulated, consistent with a function of *pitx* as an activator TF. These genes were enriched in GO terms related to synaptic transmission and axon development, suggesting that the altered gene expression related to synapse organisation may disrupt the normal functioning of neuronal circuits involved in motor control (Figure 4A, Supplementary File 8). A key gene significantly differentially regulated in this analysis was the serotonin transporter *sert* (h1SMcG0015186, Supplementary File 7).

Genes differentially expressed in *tph*+ neurons of *lhx1/5-1*(*RNAi*) animals were enriched in GO terms related to tryptophan metabolic process, response to diamide and presynaptic organisation (Figure 4B, Supplementary File 8). Notably, there is evidence concerning the influence of diamide, a sulfhydryl oxidising agent, on serotonin (5-HT) transport [55]. In particular, diamide simulates 5-HT transport in a concentration-dependent manner, with fluoxetine – a selective inhibitor of serotonin reuptake – constraining this effect [55]. A key gene significantly differentially regulated in this analysis was the serotonin synthesis rate-limiting enzyme *tph* (h1SMcG0000075, Supplementary File 7). Given the loss of *tph*+ neuronal cells (Figure 2), GO terms related to tryptophan metabolism highlight these changes are related to the synthesis of serotonin. GO terms for presynaptic organisation suggest synaptic alterations that could contribute to locomotor defects, helping to connect the gene expression changes with functional consequences in neuronal communication at the phenotypic level.

Genes differentially expressed in *otf*+ neurons 2 of *lhx1/5-1*(*RNAi*) animals were enriched in GO terms related to regulation of glutamate secretion, nitric oxide mediated signal transduction and synapse organisation (Figure 4C, Supplementary File 8). Examples of genes significantly differentially regulated in this analysis were the glutamatergic synapse regulator *unc13c* (h1SMcG0017191), the nitric oxide synthesis gene *nos1ap* (h1SMcG0011739), and the sensory ion channel *trpa1* (h1SMcG0019963, Supplementary File 7). This suggests alterations in the processes that control the release of glutamate, a key excitatory neurotransmitter. Changes in the expression of these genes may impact the balance between excitatory and inhibitory neurotransmission in the affected neurons. Nitric oxide is involved in various physiological processes, including neurotransmission, and its dysregulation can have significant implications for neuronal function [56]. Enrichment of Nitric Oxide GO terms suggests a compensatory response or an adaptive mechanism in the broader context of neurotransmitter and metabolic alterations. We performed these same analyses in the neuronal broad type in order to have a broader view of the changes in gene expression while collecting more gene counts for the analysis (Supplementary Figure 5).

Overall, these results suggest that downregulation of *pitx* and *lhx1/5-1* in serotonergic neurons trigger changes in processes tied to the locomotor defects of knockdown animals, such as excitation/inhibition balance, synapsis organisation, and neurotransmitter homeostasis.

### Indirect effects of serotonergic neuron impairment in epidermis, muscle and parenchymal cells

To assess the indirect effects of gene knockdown, we explored cell types that do not express *pitx* and *lhx1/5-1* (Figure 4D). To gain some understanding of which cell types may respond to changes in serotonin signalling, we compiled a list of genes labelled with GOs related to serotonin (Supplementary File 9) and scored their aggregated gene expression in every cell cluster. This list contains several genes associated with serotonin metabolism (both synthesis and degradation), as well as serotonin receptors and other serotonin responsive genes. This score was highest in cell clusters associated with neuronal, muscle, and parenchymal cell types (Figure 5A), which match the clusters with changes in cell numbers and gene expression levels as described previously (Figure 2B-E, Figure 3D-G). We focused on three specific clusters – cluster 3 (intestinal and DV muscle), cluster 8 (late epidermal progenitors), and cluster 7 (*pgrn*+ parenchymal cells 1) – due to their variations in cell numbers and gene expression between knockdown and control animals. To examine the genes that are differentially regulated in these groups we generated volcano plots and heatmaps, and analysed the GO term enrichment of up- and down-regulated genes (Figure 5, Supplementary File 10). Furthermore, in alignment with prior findings of impaired ciliated epithelium in these RNAi animals (Currie and Pearson, 2013), we extended our focus to the epidermal broad group for *lhx1/5-1*(RNAi) in order to have a comprehensive perspective on its systemic phenotype (Supplementary Figure 6).

**Figure 5:**
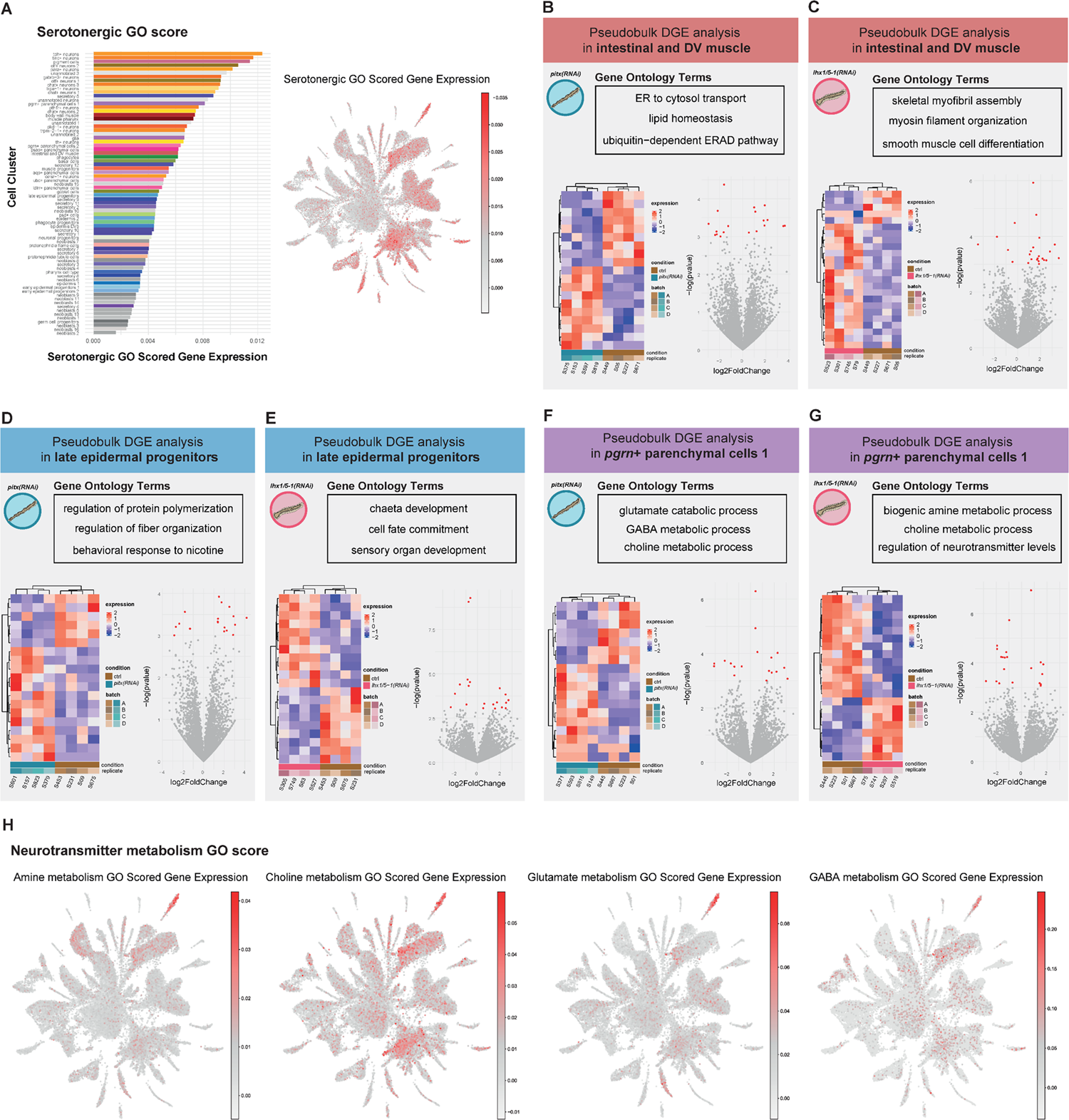
Widespread indirect effects of serotonergic neuron inhibition. **A**: Serotonergic GO score sorted barplot coloured by cell type and UMAP representation. **B-G**: Volcano plot representation, heatmap representation and GO summary of the DGE analysis in intestinal and DV muscle (B-C), late epidermal progenitors (D-E) and *pgrn*+ parenchymal cells 1 (F-G) of *pitx*(*RNAi*) animals (B, D, F) and *lhx1/5-1*(*RNAi*) animals (C, E, G). Significance levels are based on p-values < 0.05. **H**: Neurotransmitter metabolism GO scores.

**Figure 6:**
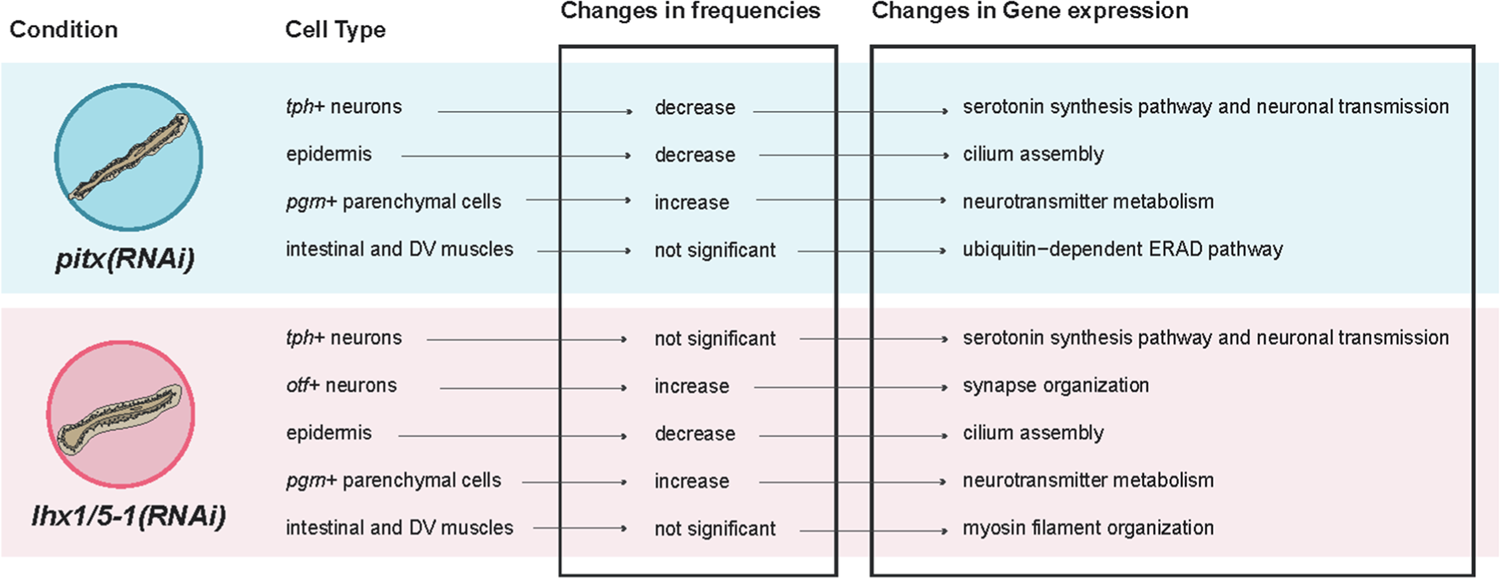
Summary of effects of serotonergic neuron inhibition

Some of the genes differentially expressed in intestinal and DV muscle of *pitx*(*RNAi*) animals were *trappc10* and *rps23* (h1SMcG0010209, h1SMcG0021056, Supplementary File 7), with GO terms related to endoplasmic reticulum (ER) and Golgi vesicular transport, as well as ubiquitination. This suggests that these specific muscle types could be facing potential damage related to cellular metabolism and protein folding (Figure 5B, Supplementary File 10).. On the other hand, genes differentially expressed in intestinal and DV muscle of *lhx1/5-1*(*RNAi*) animals were enriched in genes with GO terms directly related to myosin filament organisation and smooth muscle cell differentiation (Figure 5C, Supplementary File 10), as *unc-54* and *unc-15* (h1SMcG0000998, h1SMcG0021076, Supplementary File 7). The role of serotonin in muscle regulation, though insufficiently understood, shows indications of involvement in platyhelminth muscle activity [57, 58]. The observed close spatial relationships between peripheral serotonergic nerve elements and musculature underscore the complex influence of serotonin in these cell types. Further exploration is required to deepen our understanding of the indirect effects of serotonin transmission on the physiological function of muscles in planaria. Altogether, these analyses show that inhibition of *pitx* and *lhx1/5-1* results in molecular effects in the muscle, including genes involved in myofiber assembly.

Summary of RNAi effects of effects observed in *pitx*(*RNAi*) animals and *lhx1/5-1*(*RNAi*) animals in widespread tissues, including *tph+* neurons, epidermis, *pgrn*+ parenchymal cells and intestinal and DV muscle Genes showing differential expression in late epidermal progenitors of *pitx*(*RNAi*) and *lhx1/5-1*(*RNAi*) animals were enriched in GO terms associated with the regulation of protein polymerization, fibre organisation, and sensory organ development (Figure 5D-E). This aligns with our results as well as previous literature detailing the loss of gliding locomotion and the disruption of metachronal synchronicity (wave-like sequential coordination) in the beating cilia of planarian epithelium in the knockdown animals [24]. Examples of genes exhibiting significant dysregulation in *pitx(RNAi)* samples include *ddx54* and *unc-22* (h1SMcG0016859, h1SMcG0009472, Supplementary File 7). Conversely, in *lhx-1/5-1(RNAi)* samples, examples of dysregulated genes are *tfap4* and *fcgbp* (h1SMcG0016956, h1SMcG0011389, Supplementary File 7). We obtained similar results when we consider the epidermal broad group as a whole (Supplementary Figure 4). Overall, in this specific cell type and experimental group, a distinct and pronounced downregulation pattern is evident in genes annotated with GO terms associated with cilia movement, cilium assembly, and cilium organisation (Supplementary Figure 4). Altogether, these analyses show that inhibition of *pitx* and *lhx1/5-1* results in molecular effects in the epidermis of the worm, aligning with the observed cilia and locomotion defects.

Differentially expressed genes of *pgrn*+ parenchymal cells 1 in both knock-downs were enriched in GO terms associated with neurotransmitter metabolism. Specifically, we observed downregulation of genes related to amine, glutamate, and choline catabolism in both experimental groups (Figure 5F-G, Supplementary File 10). This suggests a function similar to that of glial cells in neurotransmitter recycling, responding to the loss of serotonergic neuronal transmission. Examples of genes significantly differentially regulated in *pitx*(*RNAi*) were the enzymes involved in the catabolism of GABA *aldh5a1* (h1SMcG0021567) and choline *dmgdh* (h1SMcG0015771), Supplementary File 7). Similarly, genes significantly differentially regulated in *lhx1/5-1*(*RNAi*) included the amino acid metabolism gene *abat* (h1SMcG0002319) and the GABA metabolism gene *pah* (h1SMcG0016738). Crucially, most differentially regulated genes involved in neurotransmitter catabolism are downregulated, indicating a possible compensatory effect.

These results indicate that the planarian parenchymal cell type *pgrn+* cells plays an important role in neurotransmitter metabolism, and that inhibition of serotonin function triggers possible compensatory effects in these cells. The role of planarian parenchymal cells is enigmatic, as there are very few molecular studies. Classical microscopy studies have shown that these types have extensions and make gap junctions with other types [59]. Recent studies have shown that these cells are phagocytic [60], play important roles in extracellular matrix function [61] and are important for regeneration [47]. Importantly, single cell studies [17, 19] have shown that these cells cluster together with planarian glial cells [47, 48]. Our results suggest that *pgrn*+ parenchymal cells may have roles in neurotransmitter catabolism. To gain support for this notion, we compiled a list of genes labelled with GOs related to amine, choline, GABA and glutamate metabolism (Supplementary File 9) and scored their aggregated gene expression in every cell (Figure 5H). This analysis showed that parenchymal cells and glial cells have high scores for these neurotransmitter metabolism gene programmes, strengthening the notion that parenchymal types have roles similar to those of glial cells. Altogether, these analyses show that upon serotonin function inhibition, parenchymal cells regulate genes involved in neurotransmitter recycling, suggesting a glial-like function of these planarian cell types.

## Discussion

In this work we used single-cell transcriptomics and genetic perturbation to study the role of serotonergic neurons with single-cell resolution at the whole organism level. Our results unveil effects in neurons but also which are the major non-neuronal cell types that respond to their inhibition. Neuron physiology sits at the interface between molecular and systems biology analyses. On one hand, detailed molecular approaches are needed to shed light into neuronal function. On the other hand, the widespread impact that neuron perturbation can have in other tissues necessitates a holistic approach like that of systems biology. At its inception, systems biology was argued as insufficient in deducing the interactions of molecular or cellular components from standalone omic observations [62]. This was based on the fact that omic techniques at the time (i) provided a unique static snapshot of the molecular features of an organism, and (ii) lacked resolution to access the cell, which is the fundamental unit of biological systems. While the omic study of dynamic scenarios is now possible thanks to the integration with genetic perturbation and differential assays [63], only recently gene expression transcriptomic studies have reached cellular resolution. However, several aspects of single-cell technologies have slowed down their widespread use in the context of dynamic scenarios, such as the effects of cell dissociation on the studied system [40–42], the affordability of replicates [35], and the lack of consensus in the appropriate statistical models for differential analysis [29, 33–35]. Our combined use of ACME and SPLiT-seq [43, 44, 51] in this work is crucial for addressing some of these problems. ACME helps to avoid the stress of enzymatic dissociation, while SPLiT-seq enables multiplexing of several samples and replicates in a single experiment, minimising batch effects and reducing cost.

Neurons serve as a key example of the complexity of the systems problem and the need for both holistic and reductionist approaches. Neurons function by interacting with each other and modulating organism-level organs and tissues. Studying neurons in isolation cannot reveal the systemic complexity of their regulation, while bulk organism-level approaches often cannot distinguish direct effects in neurons from those in the tissues they regulate. In our study, ACME and SPLiT-seq were used to analyse serotonergic neurons in the planarian *Schmidtea mediterranea*. By inhibiting the expression of *pitx* and *lhx1/5-1*, two transcription factors expressed in *tph*+ serotonergic neurons [24, 26, 53], we computationally dissect individual cell populations from a transcriptomic perspective, revealing changes in gene expression and cell abundance, and providing a holistic view of the system.

Our results uncovered genetic changes in serotonergic neurons themselves, affecting transcripts directly involved in serotonin metabolism, likely reflecting direct effects of the knockdown. These would have been challenging to elucidate from bulk RNA-seq analyses, given the tiny fraction of the serotonergic neuronal population in planarians. Additionally, we revealed cellular and molecular changes in cell types that do not express *pitx* or *lhx-1/5-1*, suggesting indirect effects from the inhibition of serotonergic neurons. These included effects in the epidermis and muscle, aligning with the described systemic effects of serotonin inhibition in planarians [24, 26, 53].

An unexpected consequence of serotonin inhibition in planarians is its impact on several parenchymal cell populations. These cell types were traditionally studied through microscopy studies in the 60s-90s [59, 64] but received minimal attention in subsequent molecular research. A 1961 study by Pedersen emphasised the striking ultrastructural resemblance of these cell types to mammalian glia [59]. In later years, parenchymal cells in planarians gained increased attention, particularly following organism-level cell type atlas studies [17, 19]. Notably, a planarian cell type labelled as glia has been explored [47, 48], with cell type atlases and lineage reconstruction studies agreeing in clustering this planarian glial type with parenchymal cells [17, 19]. This discovery has sparked controversy, as this association could be due to transcriptional similarity rather than a lineage relationship [49]. Despite the debate, the association between planarian glial and parenchymal cells has been consistently observed in multiple studies by different groups, utilising diverse dissociation techniques and single-cell transcriptomic platforms [16, 17, 19].

In our study, we reveal that serotonergic inhibition leads to an increase in the numbers of parenchymal cells and a downregulation of neurotransmitter breakdown enzymes. This suggests a potential glial-like role in neurotransmitter catabolism for these parenchymal cell types. Importantly, ultrastructural studies have demonstrated that parenchymal cells establish gap junction contacts not only with neurons but also with other cell types, including planarian neoblasts [64]. Molecular investigations have further indicated that parenchymal cells exhibit phagocytic behaviour, and inhibiting parenchymal cell transcripts has repercussions in both stem cells and planarian regeneration [47, 49, 61]. These findings open up novel avenues for exploring the function of parenchymal cells and their enigmatic relationship with glial cells.

Notably, these findings emerge from a study that marries the system-level approach with perturbation assays in order to extract information at cellular level. This showcases that ACME and SPLiT-seq are valuable methods to study complex phenotypes as they allow combining reductionistic gene manipulations with a holistic organism-level view. In time, we are confident that this approach will be scalable to more samples such as time points and multi-gene perturbations, and will extend to other single-cell omic techniques such as genome architecture and chromatin accessibility. This will greatly enhance our capability of studying the dynamics of cell behaviour and interactions in complex living systems.

## Methods

### Animal culture

All experiments were conducted using the asexual strain derived from the clonal line Berlin-1 of *Schmidtea mediterranea* [65]. Throughout the study, the animals were housed at a constant temperature of 20°C and were cultured in 1× Montjuic water composed of 1.6 mM NaCl, 1.0 mM CaCl2, 1.0 mM MgSO4, 0.1 mM MgCl2, 0.1 mM KCl, and 1.2 mM NaHCO3 in ultrapure water. The pH of the solution was adjusted to 7.5 using 1 M HCl. Regular feeding with freshly frozen calf liver was conducted every four to five days to sustain the planarians. Planarians were starved for a period of at least one week before any experimental procedure was initiated.

### Systemic RNA interference

RNA interference (RNAi) experiments were conducted with double stranded RNA (dsRNA) obtained by *in vitro* synthesis. To initiate this process, transcripts corresponding to the gene of interest were specifically selected from version 6 of the planarian “Dresden” transcriptome, which is publicly accessible on Planmine [66]. To design primers for the selected genes, we used Primer3 [67, 68].

To obtain cDNA we performed RT reactions on isolated *Schmidtea mediterranea* total RNA. Planarians were homogenised in TRIzol to isolate total RNA. Following incubation in TRIzol, we added chloroform to segregate the upper aqueous phase containing RNA. Subsequently, the RNA present in the aqueous phase was precipitated using isopropanol and dissolved in nuclease-free water to undergo further purification steps. Then, the Thermo Scientific Maxima Reverse Transcriptase was used to synthesise complementary DNA (cDNA). We incubated the extracted total RNA with deoxyribonucleotide triphosphates (dNTPs) and oligo(dT)20V at 65°C for a duration of 5 minutes, followed by immediate placement of the sample on ice for a minimum of 1 minute to facilitate primer annealing. Subsequently, a mixture of 5x Salt Buffer, Ribolock, and Maxima RT (200 U/μl) was added to the reaction, and the cDNA synthesis was carried out at 50°C for 30 minutes. The reaction was concluded with an additional incubation at 85°C for 10 minutes, which terminated the reverse transcription process. The final cDNA obtained from the experiments was appropriately diluted at a ratio of 1:5 using nuclease-free water.

dsRNA synthesis was carried out as previously described [65]. Essentially, the primary PCR mix was prepared, comprising 2 μL of cDNA, 2 μL of 10x Standard Taq Reaction Buffer (NEB), 0.4 μL of dNTPs (2.5 μM), 0.2 μL of Hot Start Taq DNA Polymerase (NEB), 4 μL of Primer Forward (2.5 μM), 4 μL of Primer Reverse (2.5 μM), and 7.4 μL of nuclease-free water. The primer sequences employed were as follows: ggccgcggGATGATATGGGACCCGACAC (lhx1/5-1-F), gccccggccGGATTCCATTCCAACCATTG (lhx1/5-1-R), ggccgcggGTCATTCTCCATCGGCTCAT (pitx-F) and gccccggccTGACAACATTGGCTGTCGAT (pitx-R). Both primer pairs were designed to encompass linkers for Universal T7 primers: ggccgcgg (linker-F) and gccccggcc (linker-R). The PCR reactions underwent thermal cycling, employing the following parameters: an initial denaturation at 94°C for 30 seconds, followed by 35 cycles of denaturation at 94°C for 20 seconds, annealing at 55°C for 20 seconds, and extension at 68°C for 30 seconds. The final extension step was carried out at 68°C for 5 minutes. Subsequently, the PCR products were subjected to electrophoresis in a 1% agarose gel to separate the amplified bands. Under UV light, the target bands were excised from the gel and placed in individual Eppendorf tubes containing 50 μL of nuclease-free water. Following a brief centrifugation of 1 minute at maximum speed, the supernatant containing the cDNA was collected.

The secondary PCR mix contained 3 μL of the cDNA, 2 μL of dNTPs (2.5 μM), 10 μL of 10x Standard Taq Reaction Buffer, 1 μL of Hot Start Taq DNA Polymerase, 82 μL of nuclease-free water, 1 μL of Universal T7-F5’ primer (25 μM, gagaattctaatacgactcactatagggccgcgg), and 1 μL of Universal T7-R3’ primer (25 μM, agggatcctaatacgactcactataggccccggc). The secondary PCR reactions were carried out in a thermocycler, utilising the following parameters: an initial denaturation at 94 °C for 30 seconds, followed by 5 cycles of denaturation at 94 °C for 20 seconds, annealing at 50 °C for 20 seconds, and extension at 68 °C for 30 seconds. This was followed by 35 cycles of denaturation at 94 °C for 20 seconds, annealing at 65 °C for 20 seconds, and extension at 68 °C for 30 seconds. A final extension step was performed at 68 °C for 5 minutes. The final product underwent purification via SPRI (Solid Phase Reversible Immobilisation) size selection in adherence to the manufacturer’s protocol and eluted in 20 μL of nuclease-free water.

By incorporating T7 promoters at both the 5’ and 3’ ends of the PCR product, the synthesis of both the sense and antisense strands of the dsRNA was facilitated through the utilisation of T7 polymerase. A precipitation protocol was employed to achieve the annealing of the complementary nucleotide chains. For each reaction, a mixture was prepared containing 1 μg of purified cDNA, 12.5 μL of 2x Express Buffer (T7 RiboMAX; Promega), 2.5 μL of Express Mix (T7 RiboMAX; Promega), and an appropriate volume of nuclease-free water to achieve a final reaction volume of up to 25 μL. The mixture was incubated for 4 hours at 37°C. Subsequently, 2.5 μL of DNase (1 U/μL, T7 RiboMAX; Promega) was added to the reaction and further incubated for 30 minutes at 37°C. To halt the reaction, 375 μL of Stop Solution (containing 1M NH4OAc, 10 mM EDTA, and 0.2% SDS) was added.

To purify the resulting double-stranded RNA (dsRNA), Phenol:Chloroform extraction was performed. For each reaction, 1 μL of GlycoBlue (to improve pellet visualisation) and 400 μL of Acid-Phenol:Chloroform (pH 4.5; Thermo Fisher) were added, and the mixture was thoroughly vortexed. After centrifugation for 5 minutes, the aqueous top phase was transferred to new tubes. Subsequently, 400 μL of chloroform was added, followed by another round of centrifugation for 5 minutes, and the top phase was collected. To precipitate the dsRNA pellets, 1 mL of cold ethanol was added to each sample, and the mixture was vortexed and then centrifuged for 15 minutes. The resulting pellets were washed in 1 mL of 70% ethanol and centrifuged for 10 minutes. The supernatants were discarded, and the pellets were allowed to air dry for 5 minutes at 37°C. Finally, the pellets were resuspended in 10-20 μL of nuclease-free water. All centrifugations throughout the process were performed at 4°C and maximum speed. As a quality check, 0.5 μL of the purified dsRNA was run on a 1% agarose gel. The concentration of the purified dsRNA was measured using a Nanodrop instrument.

To serve as controls, planarians were fed with dsRNA specific to GFP (Green Fluorescent Protein), a non-endogenous gene in planarians. For the GFP gene, a DNA miniprep of EGFP was amplified while being present in a pAGW vector. Throughout the experimental period, the planarians were fed dsRNA twice a week for a total duration of three weeks, amounting to five feedings. Each feeding involved the administration of a pellet composed of calf liver puree and low melting agarose (Thermo Fisher Scientific) at a ratio of 1:1, with a final volume of 50μL, and dsRNA at a final concentration of 1μg/μL.

### ACME cell dissociation and fixation

For each sample, approximately 20 planarians were collected and added to a 15mL Falcon tube, resulting in a final biomass volume ranging from 100–300μL. Planarian water was removed using a Pasteur pipette, and subsequently, 200μL of a 7.5% N-acetyl cysteine solution in 1× PBS was added to eliminate planarian mucus and safeguard the RNA from degradation.

Next, the ACME solution, made of DNase/RNase free water, methanol, glacial acetic acid, and glycerol in a ratio of 13:3:2:2, was introduced to each tube, reaching a final volume of 10 mL. The sample was dissociated as previously described [43]. To enhance the proportion of singlets vs aggregates, multiple filtration steps at (with a 50 nm and 40 nm filter meshes and 40 nm filter tip) were incorporated during the incubation in the ACME solution [51].

Following the incubation, the samples were centrifuged at 1000g for 5 minutes at 4 °C to remove the ACME solution. Subsequently, the cells were washed with 7mL of 1× PBS 1% BSA buffer to eliminate any remaining ACME solution. An additional centrifugation step was performed to remove the supernatant, and the final pellet was resuspended in 900μL of 1× PBS 1% BSA buffer. To preserve the cells through cryopreservation, 100μL of DMSO was added to each tube before storing them at − 80 °C.

### Flow cytometry

The dissociated samples were diluted 1:3 in 1× PBS 1% BSA buffer with a final volume of 150 μL and labelled with 0.6μL of a 1mg/mL stock solution of Alexa Fluor 488-Conjugated Concanavalin A (Invitrogen) for cytoplasm staining and with 1.5μL of a 5mM stock solution of DRAQ5 diluted 1:10 (eBioscience) for nuclear staining. This labelling process was carried out in the dark at 4°C for 30– 45 minutes. Subsequently, the stained samples were subjected to visualisation using a CytoFlex S Flow Cytometer (Beckman Coulter). During the visualisation process, a red laser (with a 780/60 nm filter) was employed for detecting DRAQ5, while a yellow-green laser (with a 525/40 nm filter) was used for detecting Concanavalin A.

### SPLiT-seq single-cell RNA sequencing

Single-cell RNA sequencing (scRNA-seq) libraries were prepared using a modified version of the SPLiT-seq protocol [44], which was further optimised based on the procedure outlined by García-Castro in 2021 [43].

#### Combinatorial Barcoding

The first round of barcoding was accomplished through in-cell reverse transcription (RT). In this process, anchored poly (dT) oligos with an extended stretch of thymidine were employed to minimise internal mispriming events (Supplementary File 1). Each reaction contained 5000 singlet events, with a volume of 8μL per well. Subsequently, a second and third round of barcoding was achieved through ligation reactions, following the methodology outlined by García-Castro in 2021. A notable modification from the original protocol was the inclusion of polyethylene glycol (PEG 8000; Promega) at a concentration of 10% w/v per reaction. This adjustment served the dual purpose of augmenting reagent concentration and facilitating cell precipitation during the pooling steps.

#### Fluorescent Activated Cell Sorting (FACS)

Following barcoding, cells were resuspended in 400-500 μL of buffer and underwent labelling with 1μL/mL of a 1mg/mL stock solution of Alexa Fluor 488-Conjugated Concanavalin A (Invitrogen) for cytoplasm staining, and 0.5μL/mL of a 5mM stock solution of DRAQ5 (eBioscience) for nuclear staining. This staining procedure was carried out in the dark at 4°C for a duration of 30–45 minutes. Subsequently, the labelled samples were sorted utilising a BD FACS Aria III (BD Biosciences) instrument. To ensure the prevention of RNAse contamination during the cell sorting process, the FACS instrument underwent thorough cleaning with bleach and pre-cooling prior to sorting. To maintain optimal conditions, both the injection and collection chambers were consistently maintained at 4 °C throughout the sorting process. Sorting was performed using the BD FACSDiva Software, configured in 4-Way Purity mode, employing an 85μm nozzle and moderate-pressure separation (45 Psi).

#### Cell lysis

Gated events characterised as DRAQ5+ and ConA+ singlets were collected and placed into 1.5 mL Eppendorf tubes, each containing 50μL of lysis buffer composed of Tris pH 8.0 (20 mM), NaCl (400 mM), EDTA pH 8.0 (100 mM), and SDS (4.4%). Each tube was capped at a maximum of 25,000 events. Following this, the volume in each tube was adjusted to 100μL, and 10μL of Proteinase K (20 mg/mL) was added to each sub-library. Subsequently, these tubes were subjected to an incubation period at 55 °C for a duration of 2 h, with manual agitation performed every 15 min to ensure proper mixing. Ultimately, the resulting lysates were stored at − 80 °C to preserve their integrity for downstream analyses.

#### Streptavidin-Biotin purification

To purify cDNA from genomic DNA, we utilised 44 μL of Dynabeads™ MyOne™ Streptavidin C1 (Invitrogen) per lysate. This purification method employs magnetic beads that contain streptavidin, which efficiently binds to the biotin moiety present at the 3’-end of the third barcode. We adhered to the manufacturer’s protocol for Dynabeads nucleic acid purification.

#### Template-switch

The template-switch reaction was conducted following the procedure previously described [43]. Additionally, to enhance the efficiency of this reaction, we incorporated polyethylene glycol (PEG 8000; Promega) at a concentration of 10% w/v in each reaction, as previously mentioned. This inclusion of polyethylene glycol aids in promoting optimal reaction conditions acting as a molecular crowding agent and thereby facilitates the successful transformation of ssDNA into dsDNA.

#### Quantitative cDNA amplification

We amplified the cDNA using 2× Kapa HiFi HotStart ReadyMix (Roche), a high-fidelity polymerase enzyme. To monitor the amplification progress, 20× EvaGreen (Biotium) was employed as the fluorescent dye, enabling real-time quantitative PCR (qPCR) analysis. The amplification process was terminated prior to reaching the exponential plateau phase, employing a strategic measure aimed at minimising the occurrence of PCR duplicates.

#### Size selection

The qPCR reactions underwent purification using SPRI size selection with Kapa Pure Beads (Roche) at specific ratios of 0.8× and 0.7× to eliminate fragments smaller than 300bp. This process was conducted following the manufacturer’s protocol. By selecting the appropriate ratios, undesired short fragments were removed, ensuring the enrichment of longer DNA fragments, which are essential for high-quality sequencing results.

To determine the concentration of the resulting libraries, quantification was performed using Qubit (Thermo Fisher). Additionally, the fragment distribution of the libraries was assessed using an Agilent 2100 Bioanalyzer, adhering to the Agilent High Sensitivity DNA Kit Guide.

#### Tagmentation and library preparation

The sub-libraries were subjected to tagmentation using the Nextera DNA Library Preparation Kit from Illumina. The input material for the tagmentation was measured to 1ng/5μL and the tagmentation reaction was carried out for 5 minutes. To halt the tagmentation process, the enzyme’s activity was immediately neutralised using the Monarch PCR & DNA Cleanup Kit (NEB). Subsequently, the samples were eluted in a final volume of 20μL of Elution buffer.

For the fourth and final round of barcoding and library preparation, PCR amplification was employed with the utilisation of tagmentation master primer i7 and library-specific tagmentation primers i5 as detailed by García-Castro in 2021. The resulting products were then subjected to size selection with Kappa Pure Beads (Roche) at specific ratios of 0.7× and 0.6×, effectively removing undesired fragments and obtaining a suitable library size distribution. The fragment distribution of the libraries was re-assessed using an Agilent 2100 Bioanalyzer (Agilent) to ensure optimal fragment lengths. To determine the concentrations of the libraries accurately, quantification was performed using Qubit (Thermo Fisher).

### Data analysis

#### Sequencing and quality control

Upon sequencing, the generated data underwent quality control and preprocessing steps, including read alignment, barcode demultiplexing, and removal of low-quality or duplicate reads. These procedures ensure the generation of high-quality data for downstream analyses. Our libraries were sequenced on a NovaSeq 6000 platform (Illumina) by Novogene, with 150 bp length, paired-end reads. The reads were provided without any quality verification apart from a basic check. Therefore, initial quality checks with FastQC software were performed. The CutAdapt v2.1 tool [69] was adopted in order to remove Illumina universal adaptors together with short and low-quality reads from read 1. Similarly, short and low-quality reads, terminal Ns and the nextera adapter sequence were removed from read 2. Makepairs was used to retain only paired reads.

#### Read mapping, barcode extraction, and matrix generation

We mapped the reads to the new version of the *Schmidtea mediterranea* genome [70]. Each of the sequenced sub-libraries was processed individually. To extract and revise the barcodes, we employed the SPLiTseq toolbox, which incorporates many components from Drop-seq_tools-2.3.05. Subsequently, each sub-library was mapped using STAR (--quantMode GeneCounts) [71], and further reordered and merged using Picard v2.20.5 SortSam and MergeBamAlignment.

For mapping location tagging, we utilised Drop-seq tools-2.4.0 TagReadWithInterval and TagReadWithGeneFunction. To generate the expression matrices for each library individually, we employed Drop-seq tools-2.4.0 with the following settings: READ_MQ=0, EDIT_DISTANCE=1, MIN_NUM_GENES_PER_CELL=50, and LOCUS_FUNCTION_LIST=INTRONIC. These steps enabled the generation of expression profiles for each library.

#### Cell clustering and identification of cell types

The cell per gene matrices were uploaded into a Python Jupyter Notebook, using the toolkits Scanpy, pandas and NumPy. Each library was normalised, log transformed and variable genes detected. Unsupervised clustering methods, such as Principal Component Analysis (PCA) and Uniform Manifold Approximation and Projection for Dimension Reduction (UMAP), were employed to group cells based on their gene expression profiles. PCA was run for 150 principal components, and an elbow plot generated to check for the variance ratio among these PCs. Neighbours were calculated considering the first 95 PCs, the UMAP visualisation was generated (min_dist = 0.75, spread = 1.25) and clustering performed with the Leiden algorithm (resolution = 1, 2, 3, and 4). Markers were extracted by ranking genes for characterising groups (method=’wilcoxon’ and method=’logreg’, Supplementary File 2). By comparing the expression patterns of known marker genes, cell types were assigned to each cluster.

#### Differential Gene Expression (DGE) analysis

We created a pseudo-bulk matrix leveraging the cell annotation data derived from our single-cell analysis (e.g. clustering and broad cell types) along with additional cell information like sample characteristics (e.g. type, replicates). This was achieved by aggregating, for a given gene X, cell type Y, experiment I, and replicate J, all the counts of gene X from cells belonging to the same cluster Y and under identical conditions (experiment I and replicate J). This process effectively generated a matrix with genes arranged in rows and ‘pseudo-samples’ (combinations of cell type, experiment, and replicate) in columns. These pseudo-samples encompassed various combinations of cell types and conditions, such as RNA interference treatment and replicates (e.g. biological, technical, library etc.).

We performed DGE analysis for a specific cell type y as follows: (i) we extracted the relevant pseudo-samples of cell type y from the pseudo-bulk matrix.; (ii) we also filtered the matrix from (i) to keep genes identified as “high-confidence” in the latest *Schmidtea* genome annotation [70]; (iii) we filtered out genes with less than two counts in at least one replicate; (iv) we ran DESeq2 [54] inside a custom wrapper with a contrast of “condition 1” (RNAi of either *lhx1/5-1* or *pitx*) relative to “condition 2” (*control RNAi*). Genes were identified as differentially expressed if having a p-value below 0.05 (negative binomial test).

#### Statistical analysis

In order to assess differences in cell type abundances between experimental and control conditions, we employed the Fisher’s Exact Test. To perform the test, contingency tables were constructed, with rows representing the experimental and control conditions and columns representing the presence or absence of each cell type. We adhered to a significance level of p < 0.01.

## Availability of Data and Materials

The datasets supporting the conclusions of this article are available in: Code: https://github.com/scbe-lab/planarian-serotonergic-rnai GEO: GSE256032

## Competing Interests

The authors declare that they have no competing interests.

## Funding

Research at the Solana lab at Oxford Brookes University is supported by MRC grants (MR/S007849/1 and MR/W017539/1), a BBSRC Grant (BB/V014447/1) and a Leverhulme Trust grant (RPG-2019-332) to JS. EE was supported by Nigel Groome studentship from Oxford Brookes University. DRF was supported by an Oxford Interdisciplinary Biosciences DTP studentship.

## Acknowledgments

We thank Robert Hedley and Vasiliki Tsioligka at the Flow Cytometry Facility at the Dunn School of Pathology (University of Oxford) and Vincent Mason for technical assistance. We thank Jochen Rink and Luca Pandolfini for early access to the new *Schmidtea mediterranea* genome and annotation. We thank all other members of the Stem Cell Biology and Evolution team for discussion and support.

## Authors’ Contributions

JS and EE conceived the study and designed the experiments. DR-F prepared the two biological replicates of the RNAi samples assisted by EE. EE generated one technical replica of the cell dissociations and DR-F prepared the other one. EE performed single-cell transcriptomic experiments. EE conducted the pre-processing of the reads and the mapping, and curated the cell clustering supervised by JS. EE executed differential expression analysis in pseudo-bulk supervised by AP-P and A.P-P re-analysed the available bulk RNA-seq data. EE performed the statistical analysis. An earlier round of phenotypic screening was performed by EE, assisted by HGC.

## Supplementary Figure Legends

**Supplementary Figure 1.**
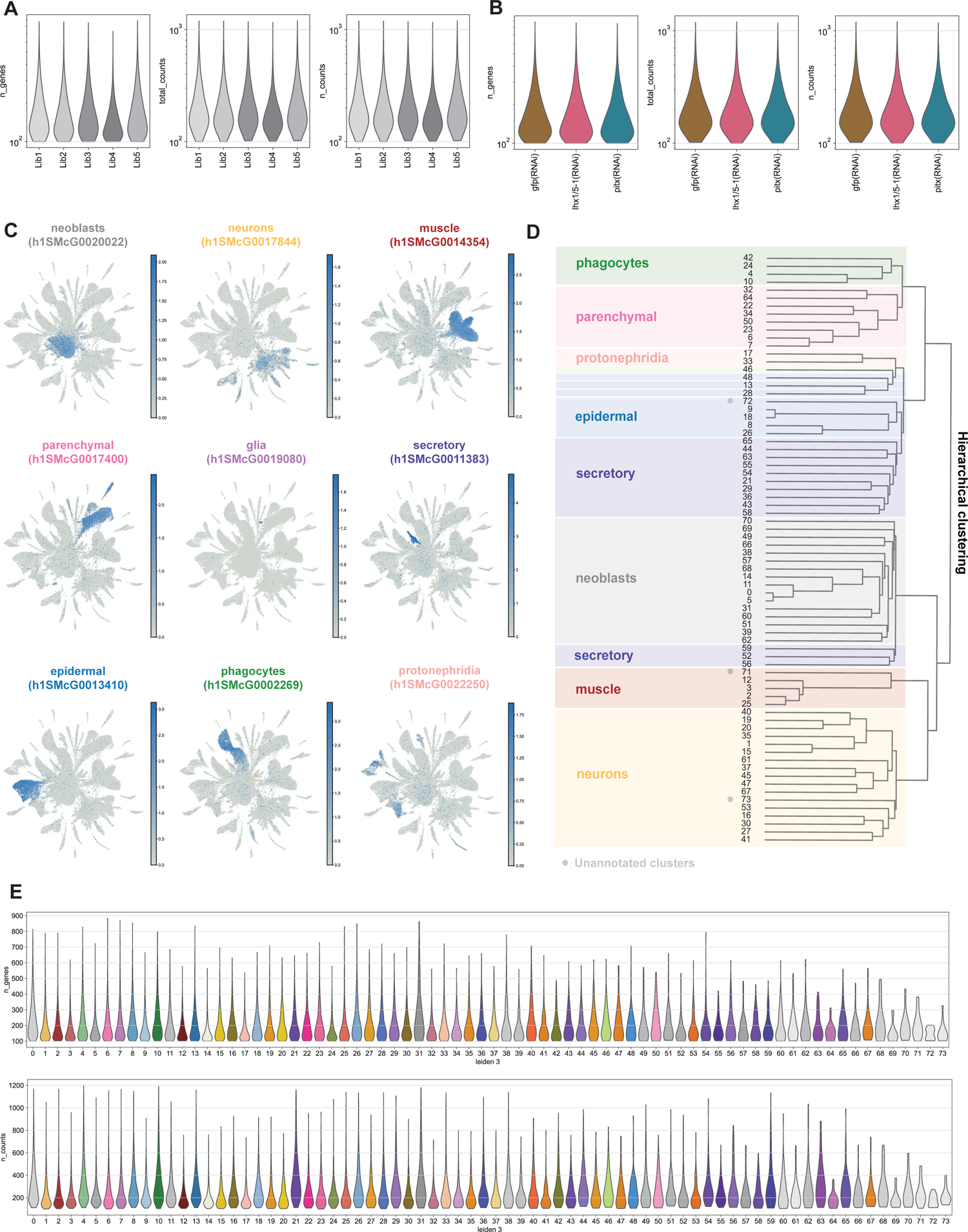
Serotonergic dataset metrics. A: Violin plots showing the number of genes and UMI counts detected per sub-library. B: Violin plots showing the number of genes and UMI counts detected per sample. C: UMAP features plots of *Schmidtea mediterranea* cell type diagnostic marker genes. D: Hierarchical clustering dendrogram of cell clusters to define broad group identities. E: Violin plots showing the number of genes detected per cell in each cluster (upper). Violin plots showing the number of UMI counts detected per cell in each cluster (lower).

**Supplementary Figure 2.**
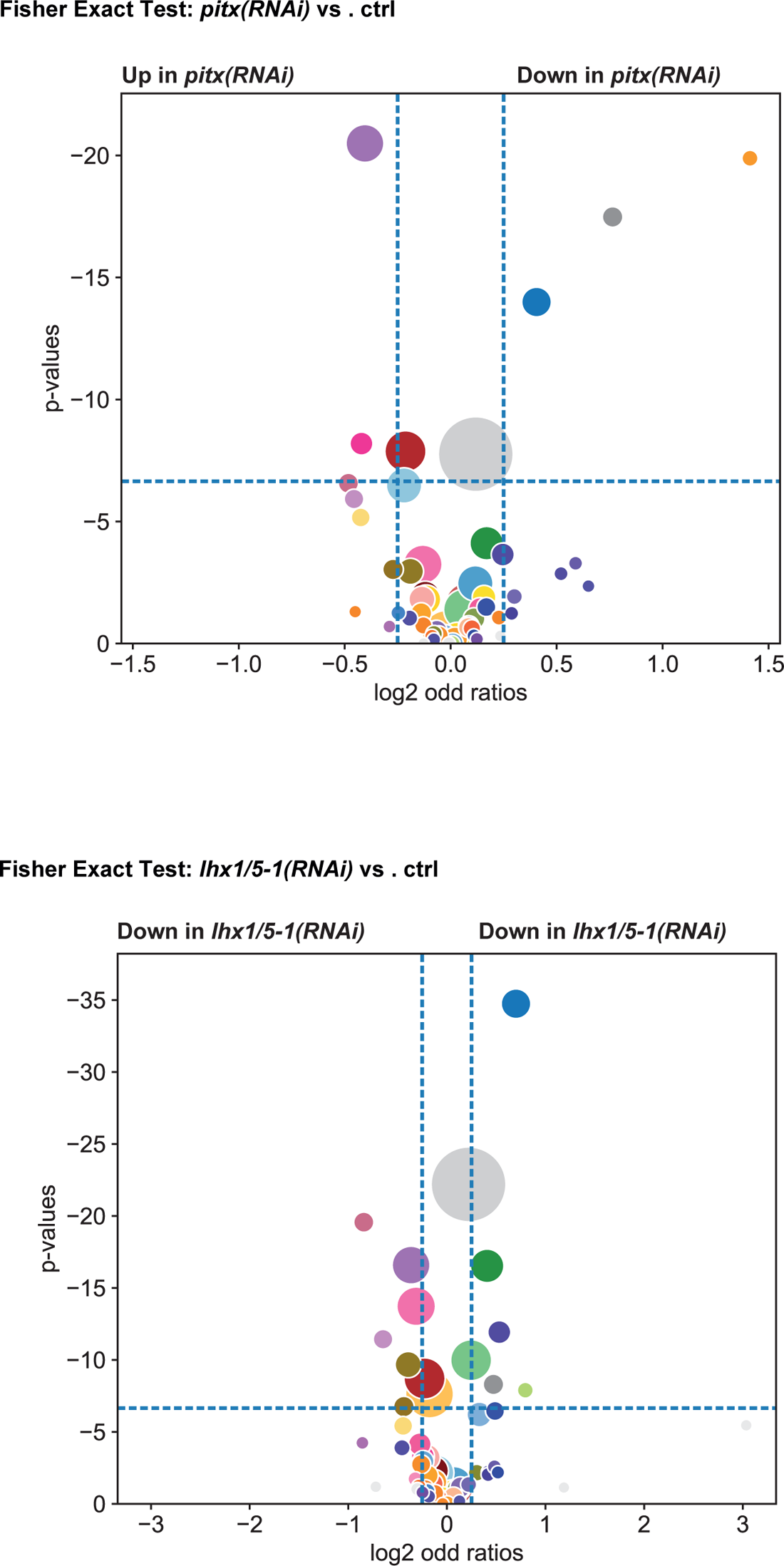
Volcano-plot representation of the results of the Fisher’s Exact Test examining differences in cell frequencies between *pitx(RNAi)* and controls animals (upper), and *lhx1/5-1(RNAi)* and controls animals (lower).

**Supplementary Figure 3.**
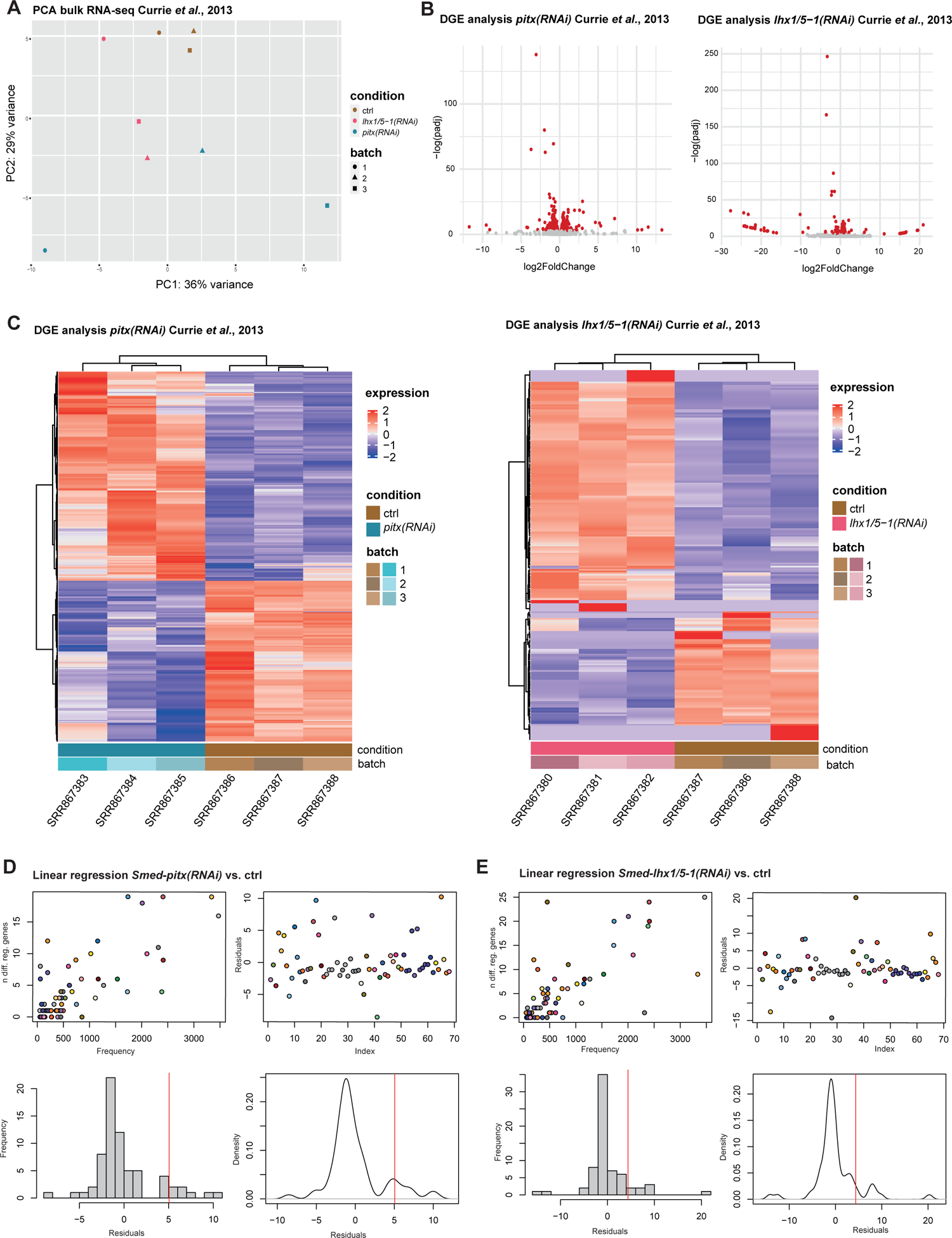
Differential gene expression analysis with DEseq2. A: Principal Component Analysis of Currie and Pearson (2023) bulk RNA-seq displaying variability among replicates for both *pitx*(*RNAi*) and *lhx1/5-1*(*RNAi*). B-C: Volcano and Heatmaps displaying differentially regulated genes for both *pitx*(*RNAi*) and *lhx1/5-1*(*RNAi*). D-E: Linear regression of scatter plot distribution showing the number of differentially regulated genes per cluster compared to the log-transformed cell count within each cluster for both *pitx*(*RNAi*) and *lhx1/5-1*(*RNAi*) in this study.

**Supplementary Figure 4.**
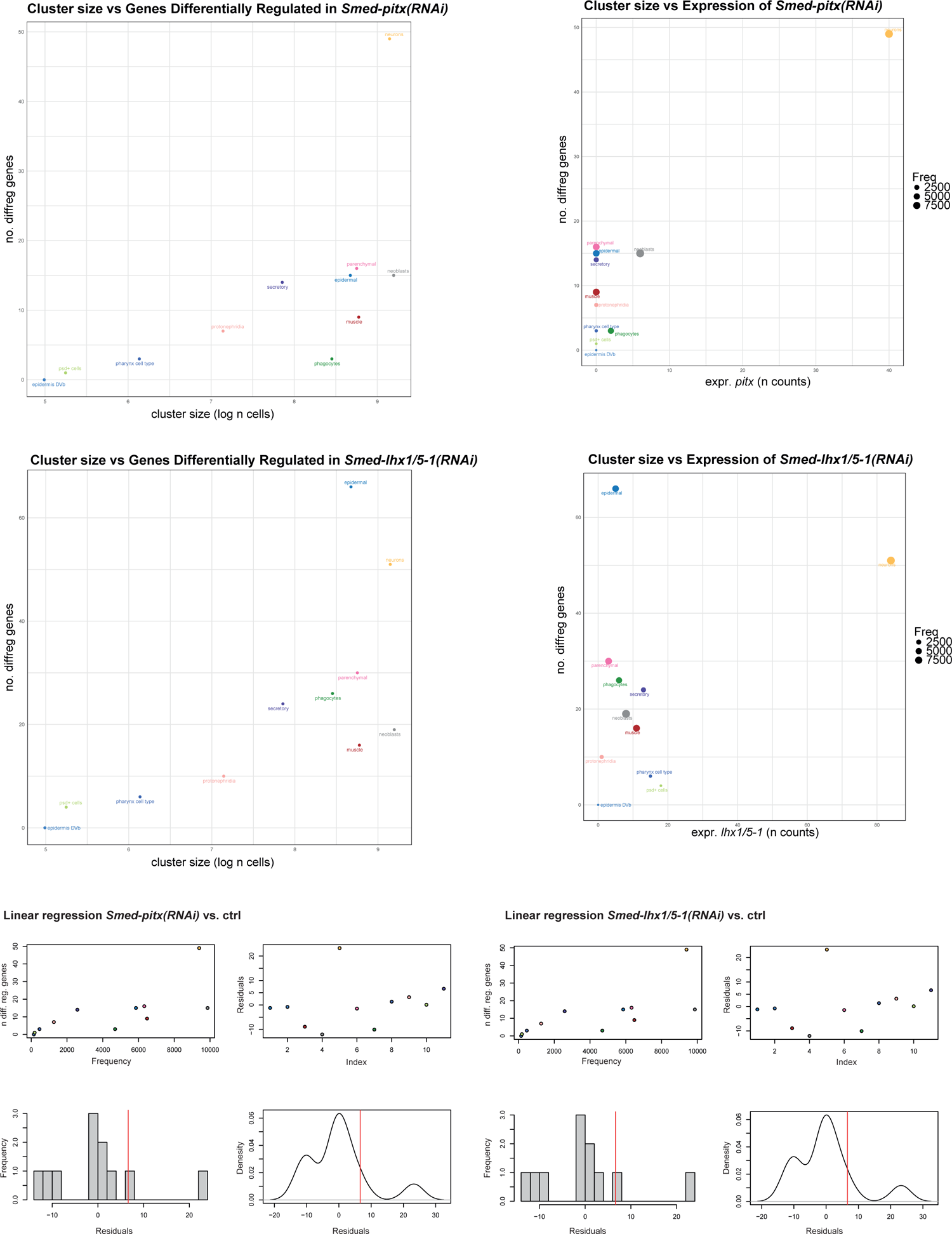
Scatter plot showing the number of differentially regulated genes per broad group compared to the log-transformed cell count within each broad group in both *pitx*(*RNAi*) (up) and *lhx1/5-1*(*RNAi*) (down) DGE analysis. Scatter plot showing the number of differentially regulated genes per broad group compared to the expression of *pitx* (right) and *lhx1/5-1* (left). Linear regression of scatter plot distribution showing the number of differentially regulated genes per cluster compared to the log-transformed cell count within each cluster for both *pitx(RNAi)* and *lhx1/5-1(RNAi)* respectively.

**Supplementary Figure 5.**
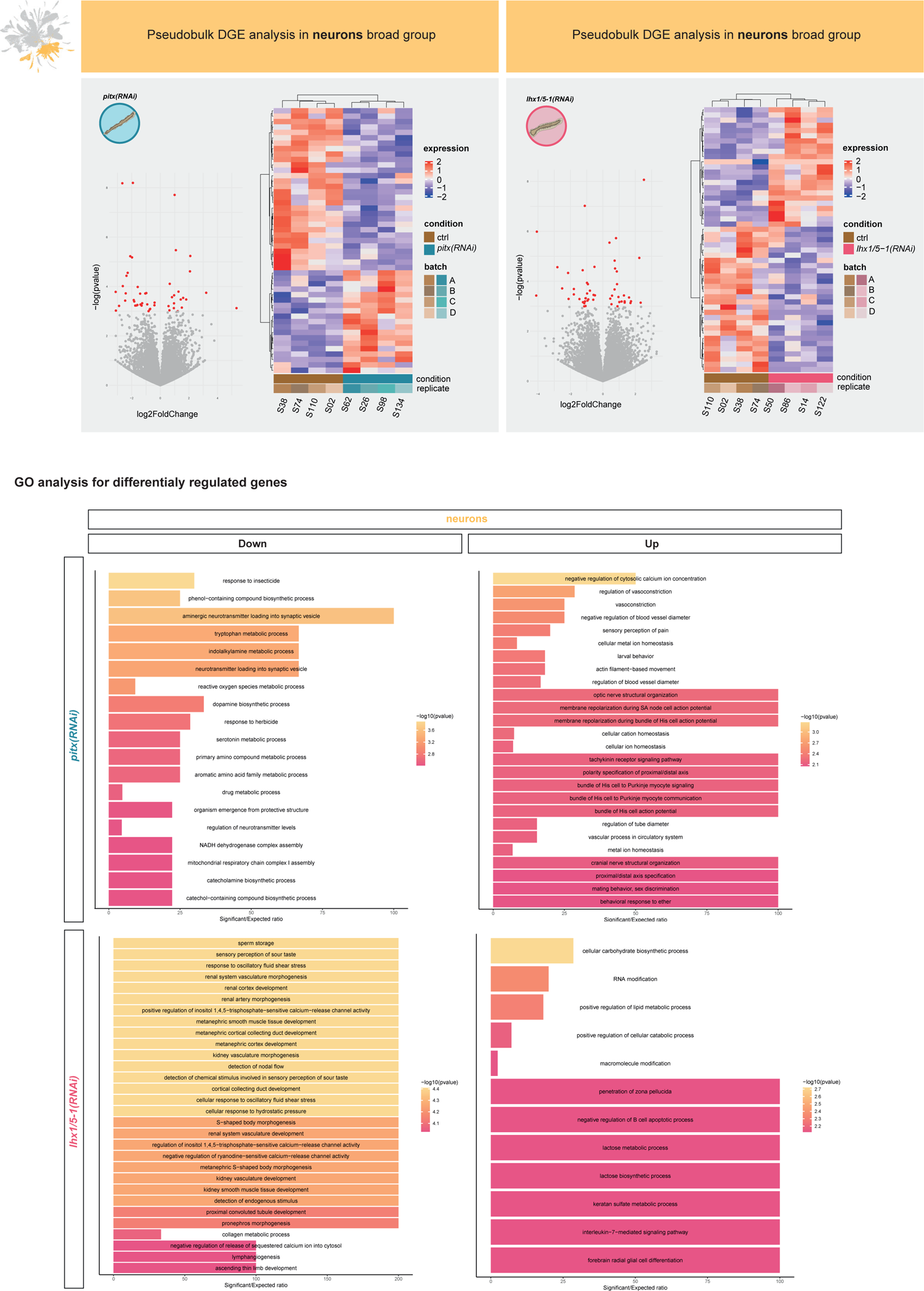
Direct effects of the loss of gene function. Volcano plot representation, heatmap representation and GO analysis summarising DGE analysis in the neuron broad group of *pitx*(*RNAi*) animals and *lhx1/5-1*(*RNAi*) animals.

**Supplemental Figure 6.**
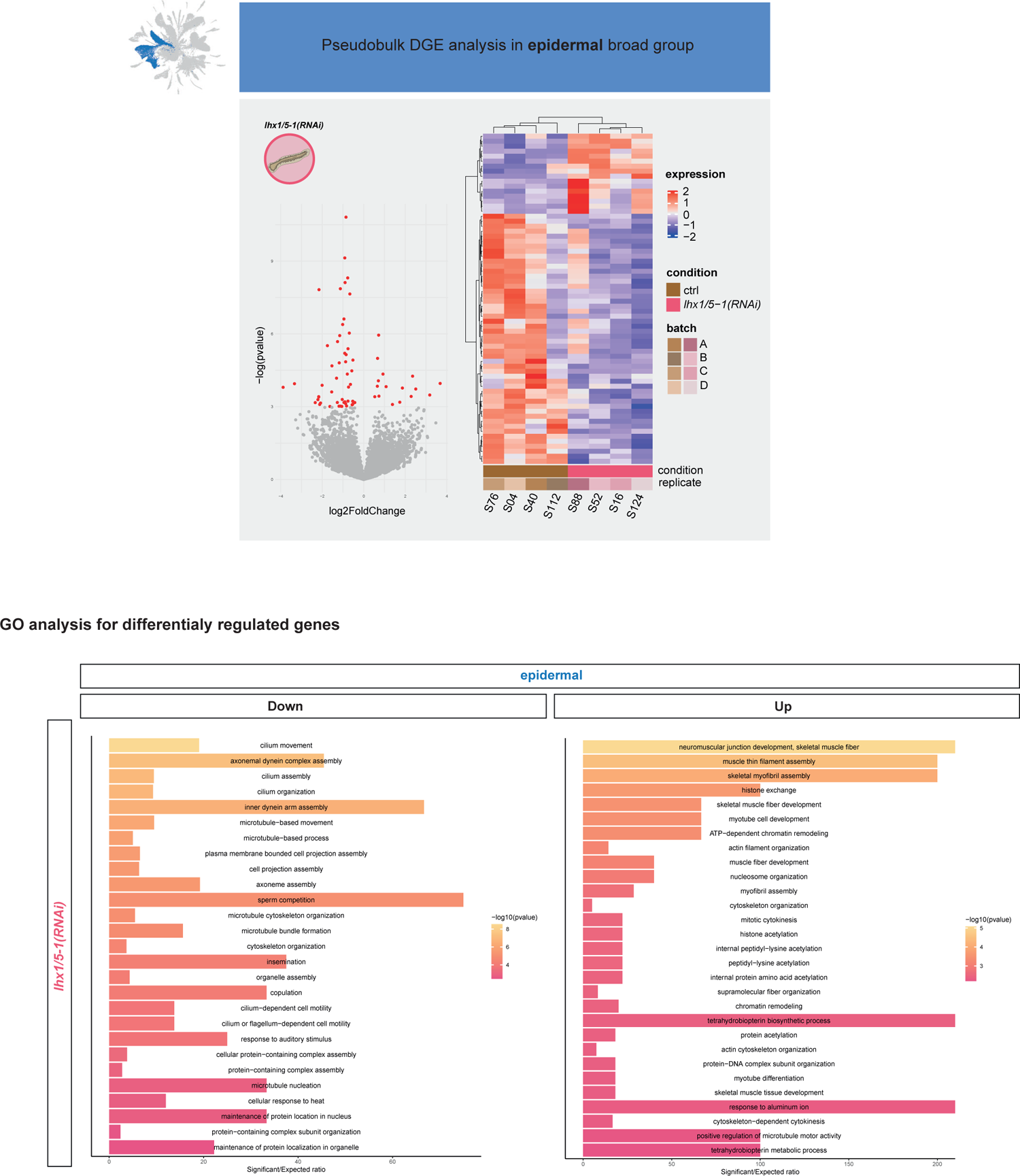
Indirect effects of the loss of gene function. Volcano plot representation, heatmap representation and GO analysis summarising DGE analysis in epidermal broad groups of *pitx*(*RNAi*) animals and *lhx1/5-1*(*RNAi*) animals.

## Supplementary File names

**Supplementary File 1.** Oligonucleotides for 96×96×96×5 Combinatorial Barcoding

**Supplementary File 2.** Cluster Wilcoxon and Logistic Regression markers

**Supplementary File 4.** Top cluster markers UMAP visualisations

**Supplementary File 5.** Neuronal markers UMAP visualisations

**Supplementary File 6.** Bulk DEseq2 Differential Gene Expression re-analyses of Currie and Person, 2013.

**Supplementary File 7.** Single-cell DEseq2 Differential Gene Expression analyses

**Supplementary File 8.** GOs of Differentially Expressed Genes in *tph*+ neurons and *otf*+ neurons

**Supplementary File 9.** GOs and gene IDs related to serotonin, amine, choline, GABA, and glutamate metabolism.

**Supplementary File 10.** GOs of Differentially Expressed Genes in *pgrn*+ cells late epidermal and dv muscle

## Notes

https://github.com/scbe-lab/planarian-serotonergic-rnai

### Competing Interest Statement

The authors have declared no competing interest.

