## Supplementary material for "Multiplex single-cell analysis of serotonergic neuron function in planarians reveals widespread effects in diverse cell types": Emili et al 2024 Serotonin Supplementary File 4 20240225 vs1.pdf

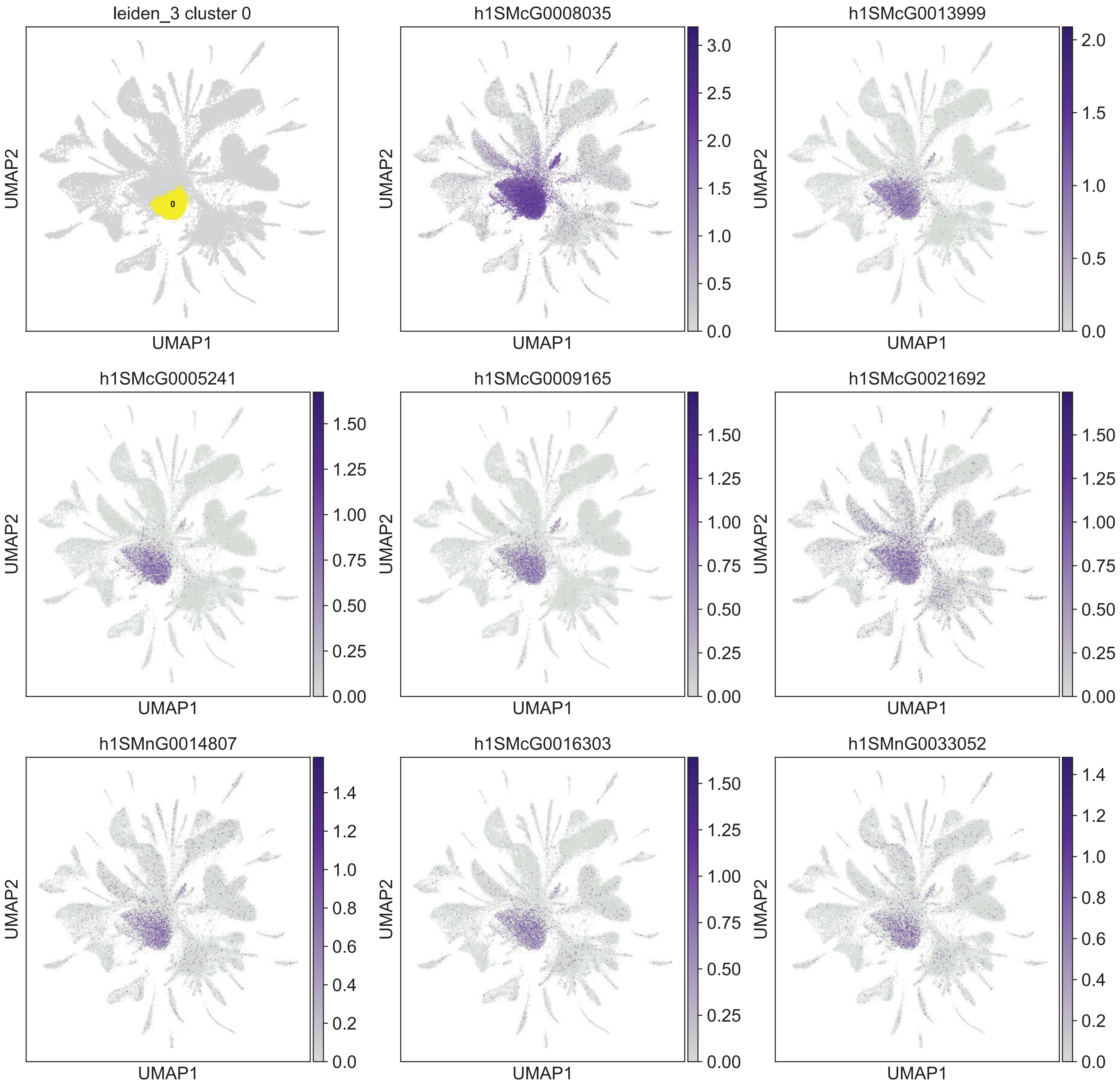

leiden_3 cluster 0
0
h1SMcG0008035
h1SMcG0013999
h1SMcG0005241
h1SMcG0009165
h1SMcG0021692
h1SMnG0014807
h1SMcG0016303
h1SMnG0033052
UMAP1
UMAP2

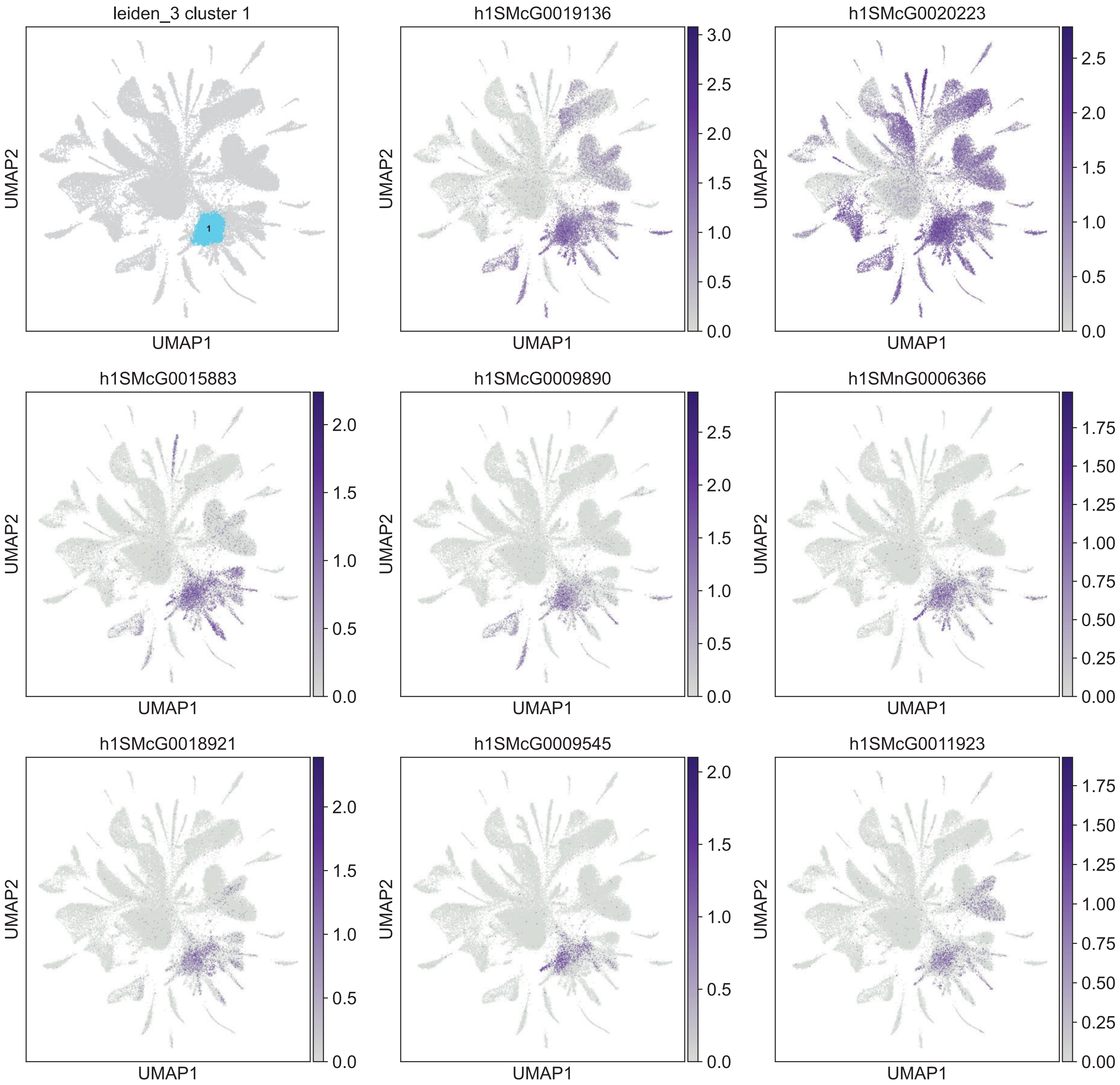
leiden_3 cluster 1
1
h1SMcG0019136
h1SMcG0020223
h1SMcG0015883
h1SMcG0009890
h1SMnG0006366
h1SMcG0018921
h1SMcG0009545
h1SMcG0011923
UMAP1
UMAP2
0.0
0.5
1.0
1.5
2.0
2.5
3.0
0.00
0.25
0.50
0.75
1.00
1.25
1.50
1.75

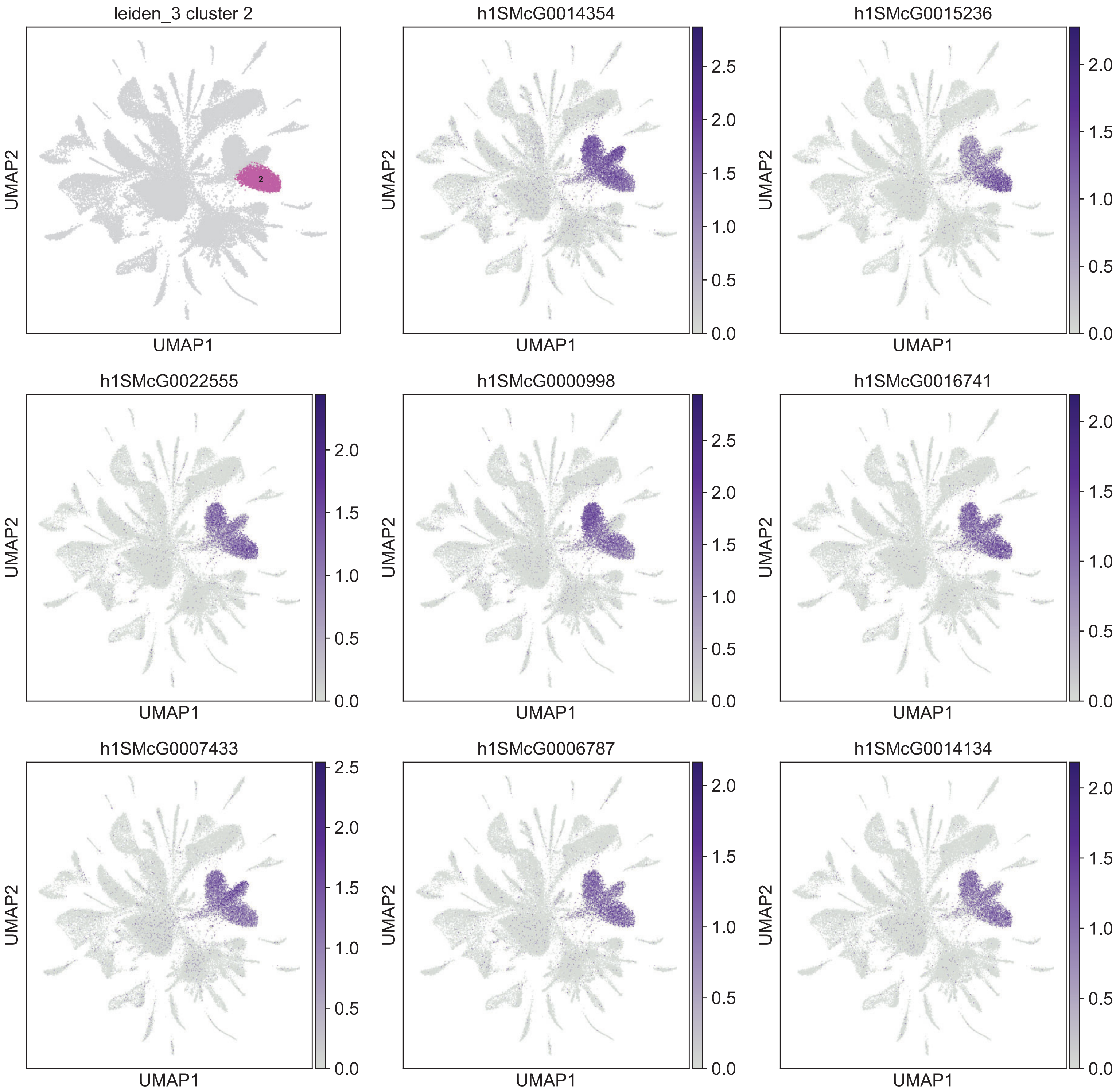

leiden_3 cluster 2
h1SMcG0014354
h1SMcG0015236
h1SMcG0022555
h1SMcG0000998
h1SMcG0016741
h1SMcG0007433
h1SMcG0006787
h1SMcG0014134
UMAP1
UMAP2

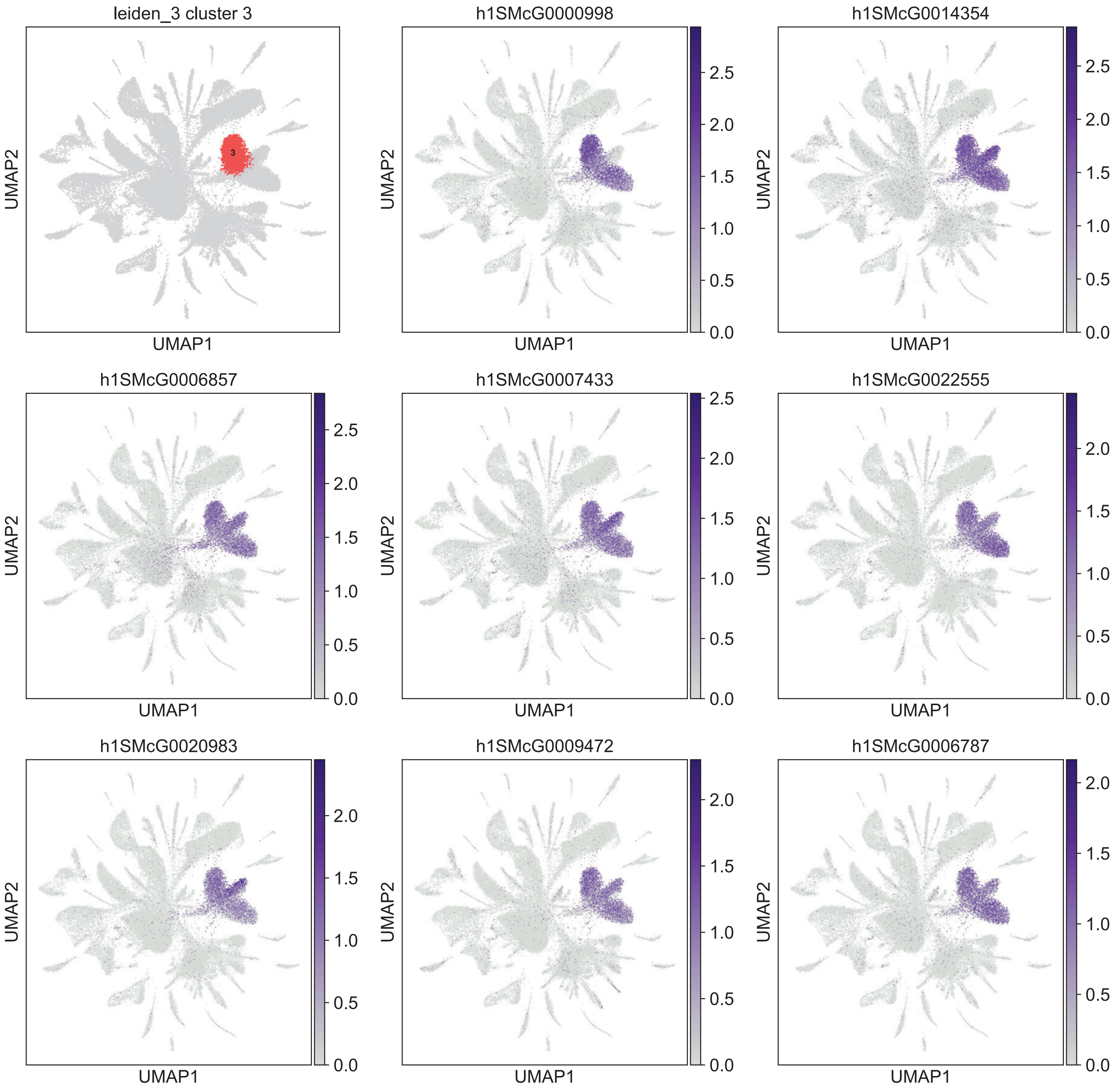

leiden_3 cluster 3
h1SMcG0000998
h1SMcG0014354
h1SMcG0006857
h1SMcG0007433
h1SMcG0022555
h1SMcG0020983
h1SMcG0009472
h1SMcG0006787
UMAP1
UMAP2

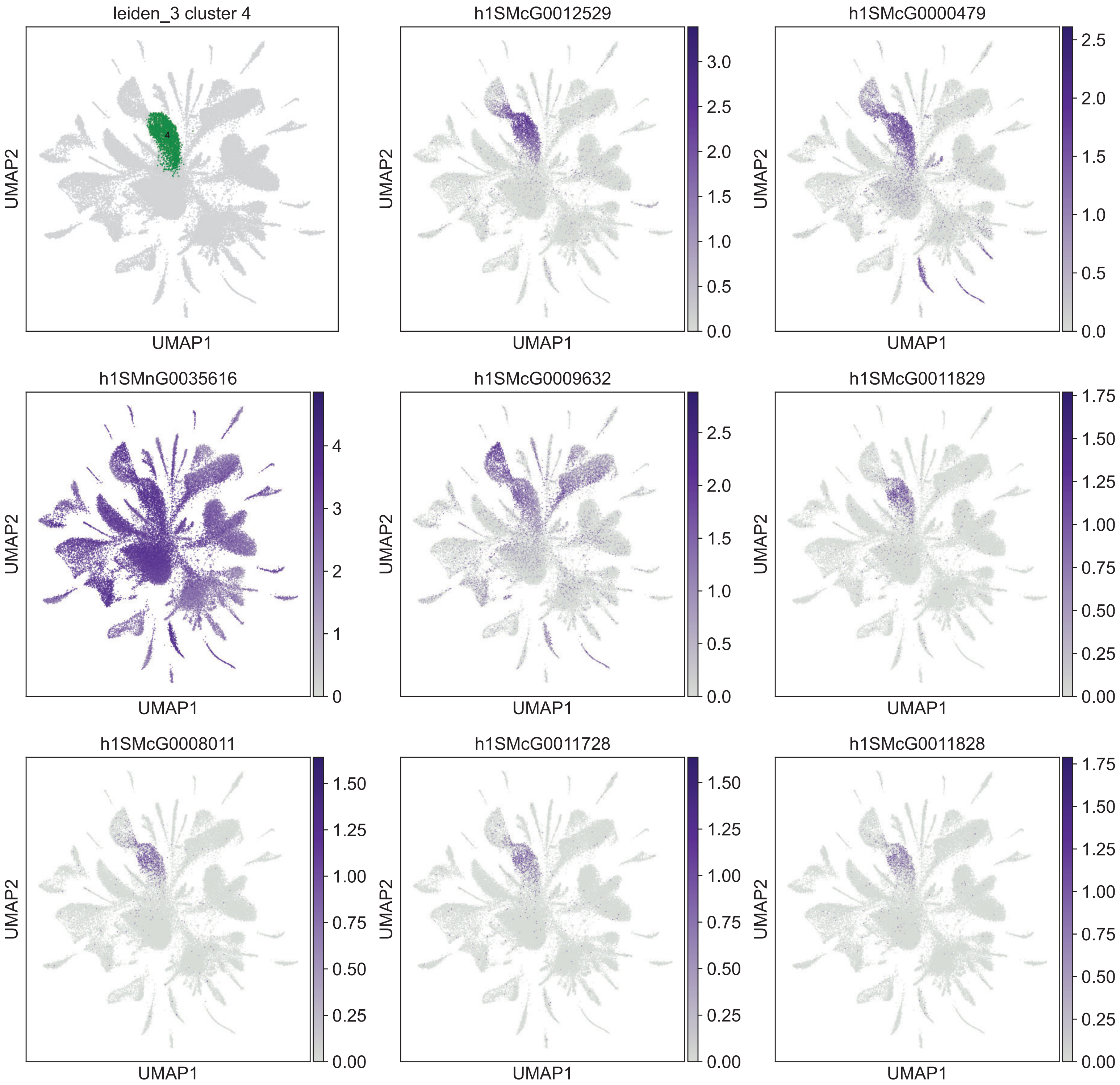
leiden_3 cluster 4
h1SMcG0012529
h1SMcG0000479
h1SMnG0035616
h1SMcG0009632
h1SMcG0011829
h1SMcG0008011
h1SMcG0011728
h1SMcG0011828
UMAP1
UMAP2

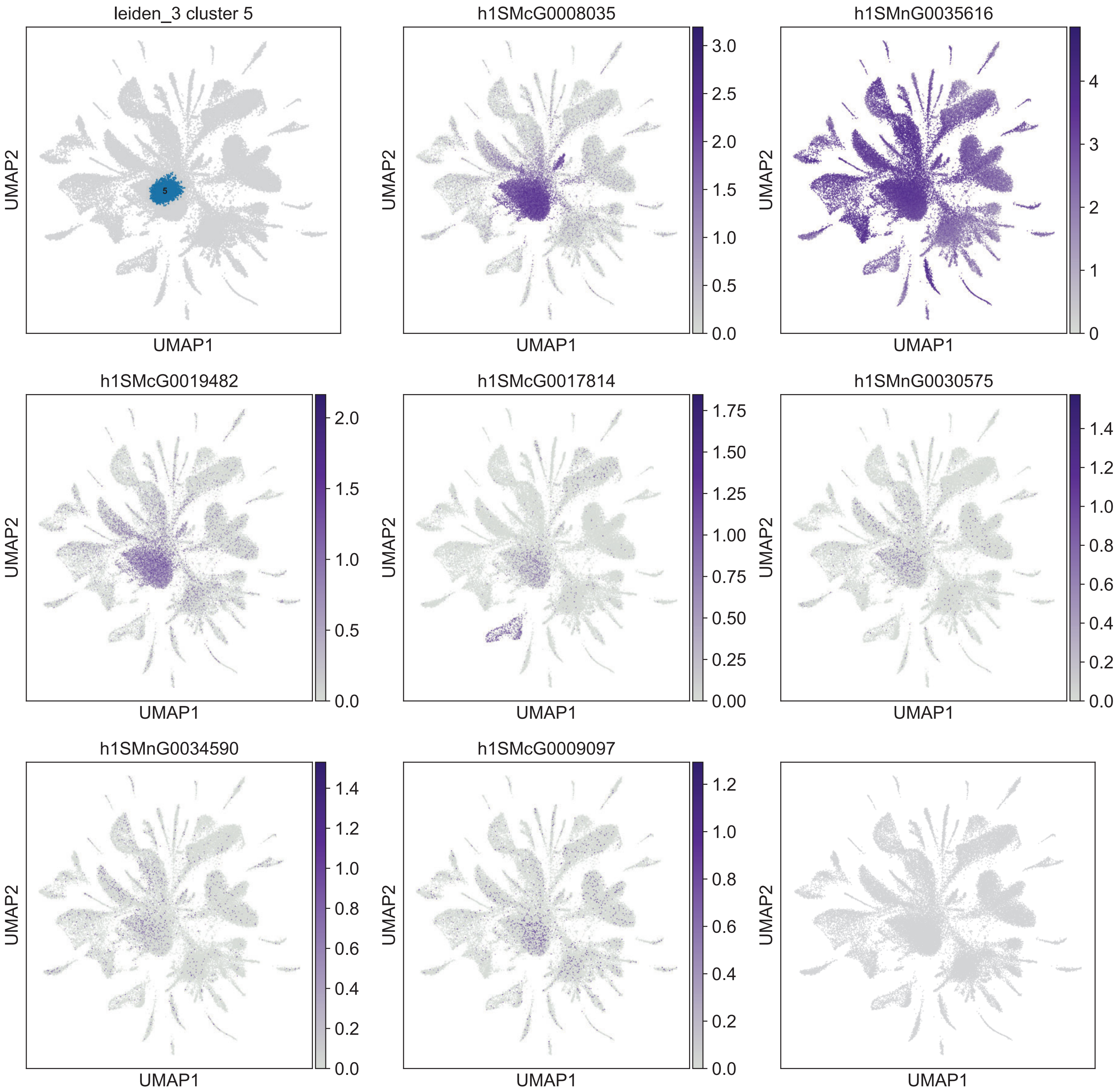
leiden_3 cluster 5
5
h1SMcG0008035
h1SMnG0035616
h1SMcG0019482
h1SMcG0017814
h1SMnG0030575
h1SMnG0034590
h1SMcG0009097
UMAP1
UMAP2

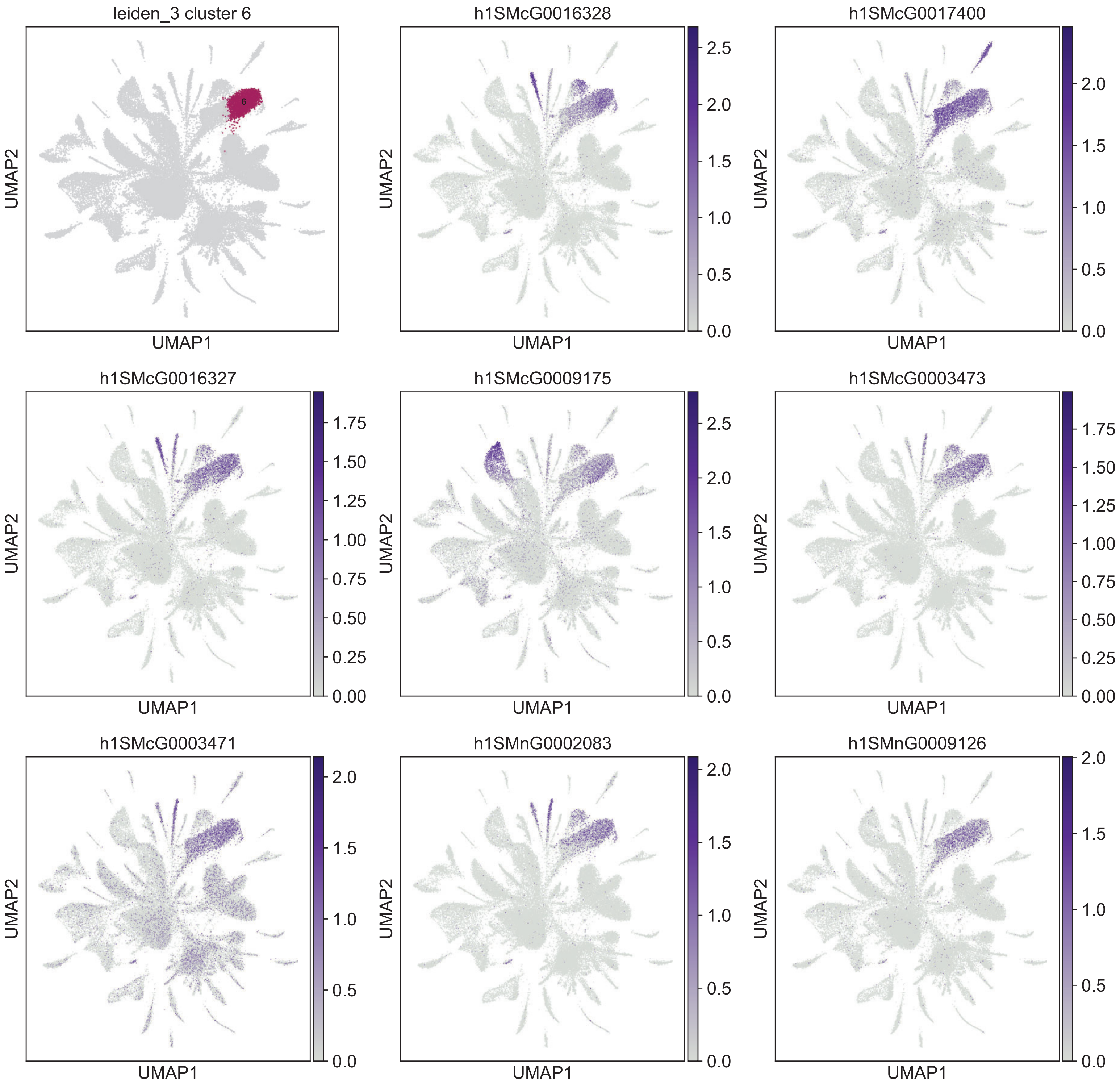
leiden_3 cluster 6
6
h1SMcG0016328
h1SMcG0017400
h1SMcG0016327
h1SMcG0009175
h1SMcG0003473
h1SMcG0003471
h1SMnG0002083
h1SMnG0009126
UMAP1
UMAP2

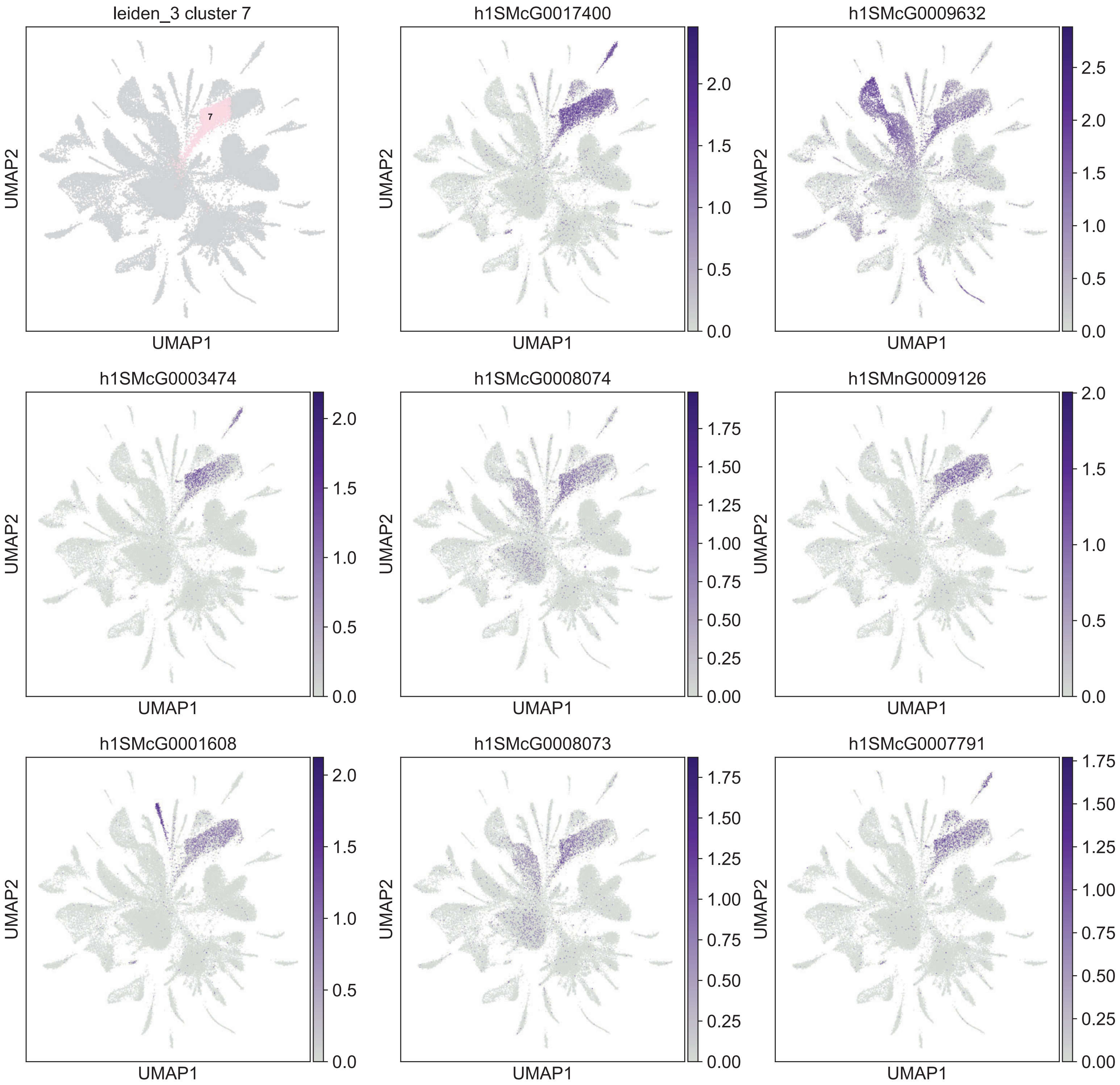

leiden_3 cluster 7
7
UMAP2
UMAP1
h1SMcG0017400
h1SMcG0009632
h1SMcG0003474
h1SMcG0008074
h1SMnG0009126
h1SMcG0001608
h1SMcG0008073
h1SMcG0007791

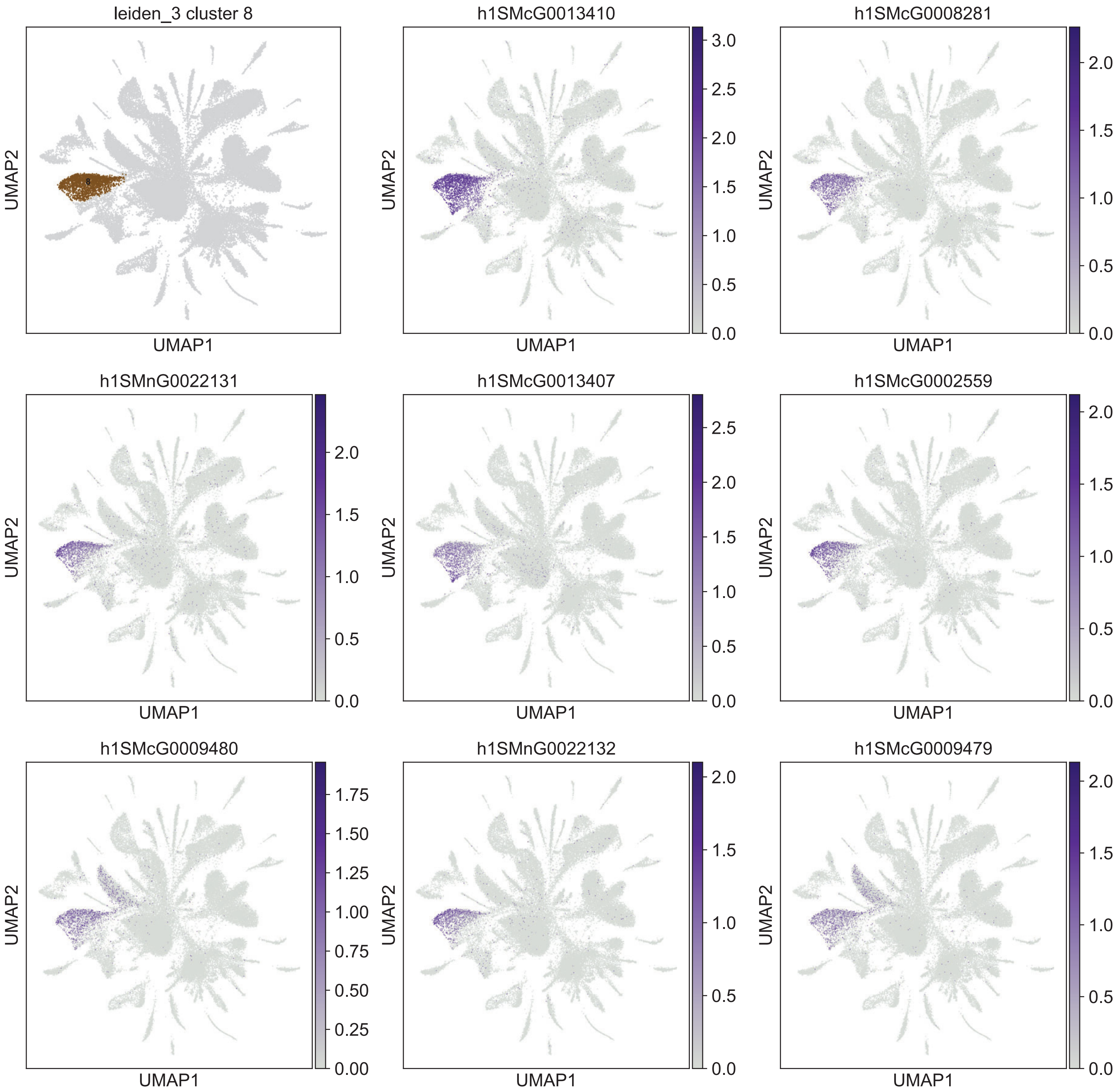
leiden_3 cluster 8
h1SMcG0013410
h1SMcG0008281
h1SMnG0022131
h1SMcG0013407
h1SMcG0002559
h1SMcG0009480
h1SMnG0022132
h1SMcG0009479
UMAP1
UMAP2
0.0
0.5
1.0
1.5
2.0
2.5
3.0
0.00
0.25
0.50
0.75
1.00
1.25
1.50
1.75

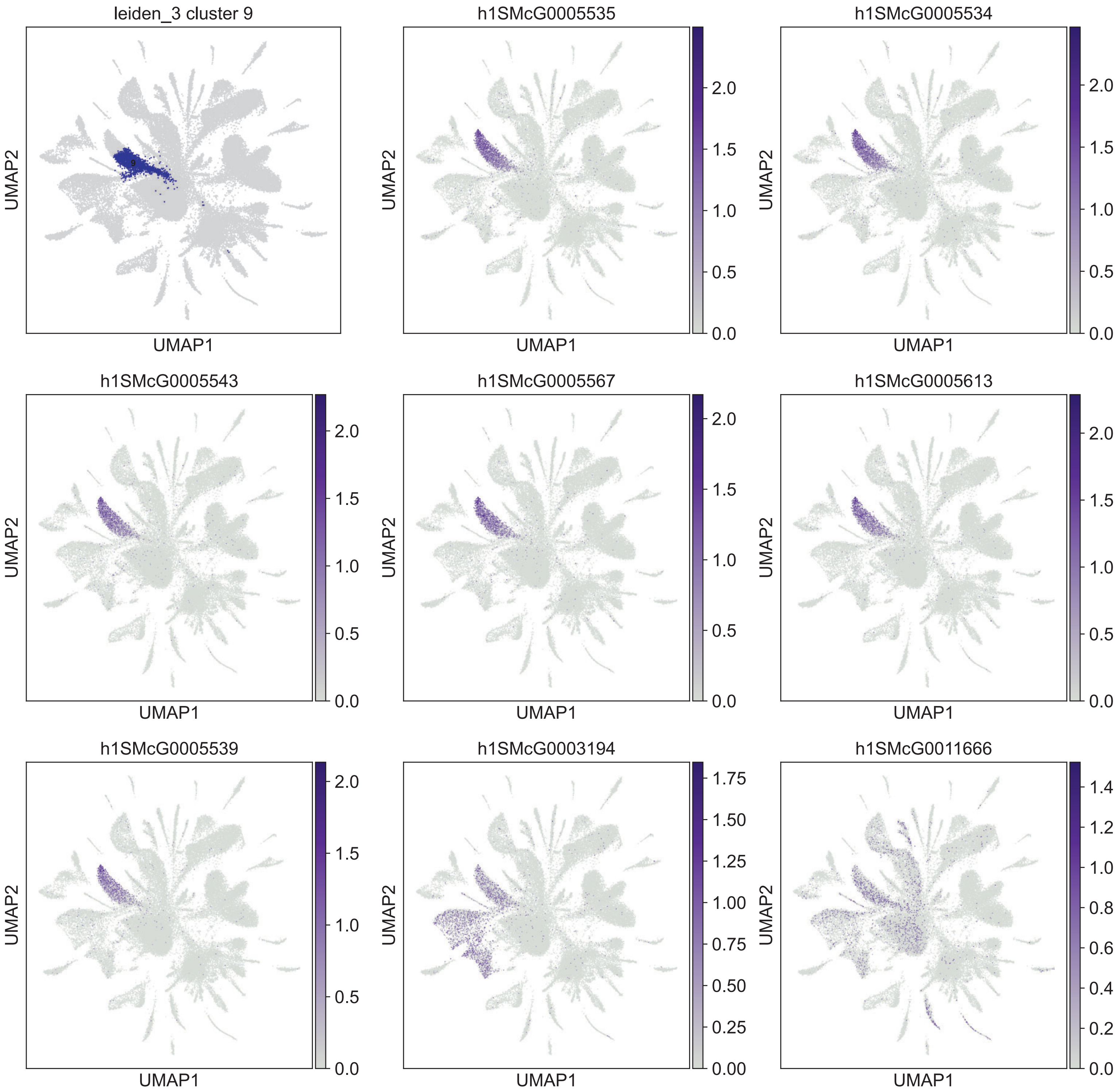
leiden_3 cluster 9
h1SMcG0005535
h1SMcG0005534
h1SMcG0005543
h1SMcG0005567
h1SMcG0005613
h1SMcG0005539
h1SMcG0003194
h1SMcG0011666
UMAP1
UMAP2

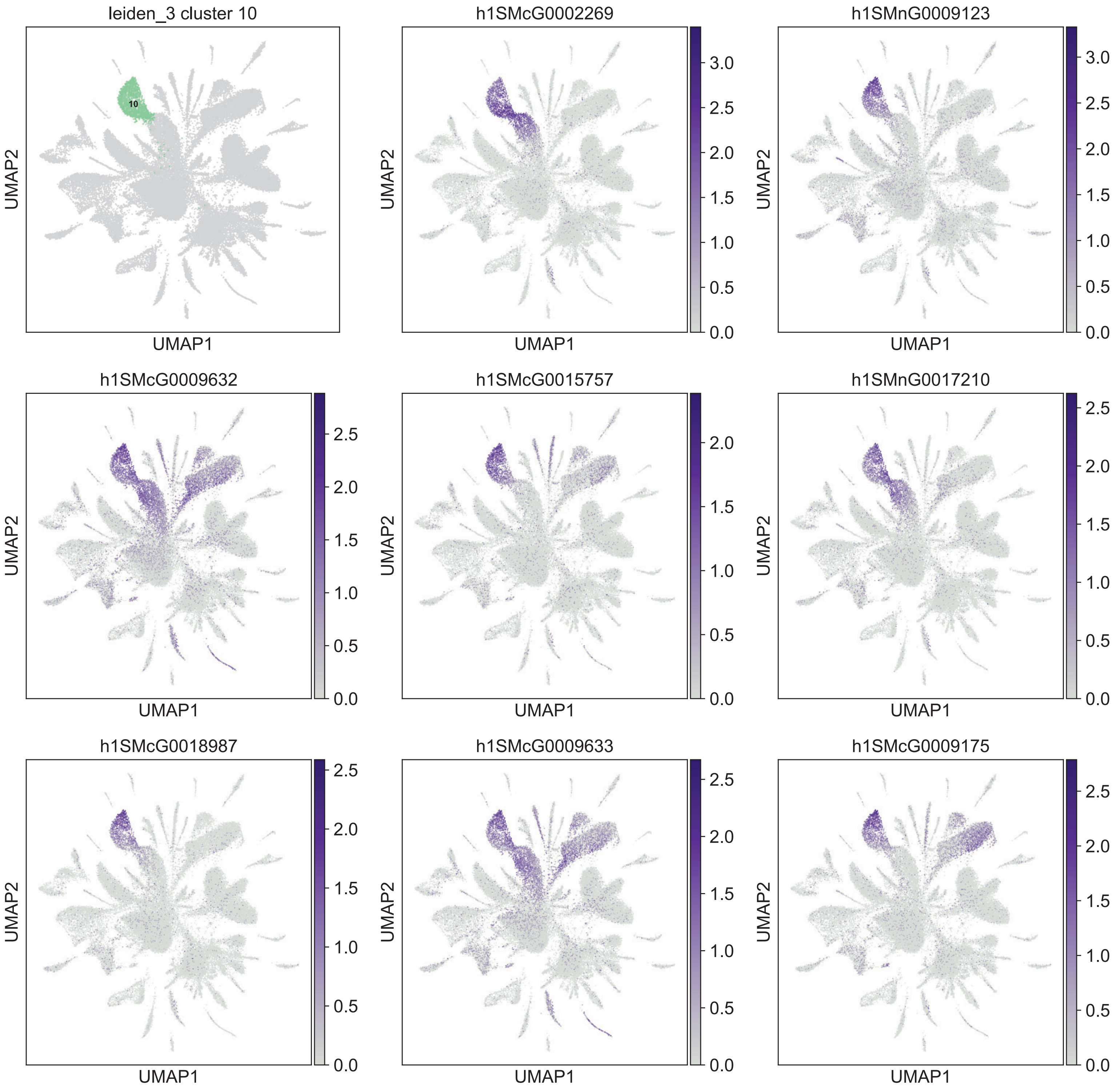
leiden_3 cluster 10
10
h1SMcG0002269
h1SMnG0009123
h1SMcG0009632
h1SMcG0015757
h1SMnG0017210
h1SMcG0018987
h1SMcG0009633
h1SMcG0009175
UMAP1
UMAP2

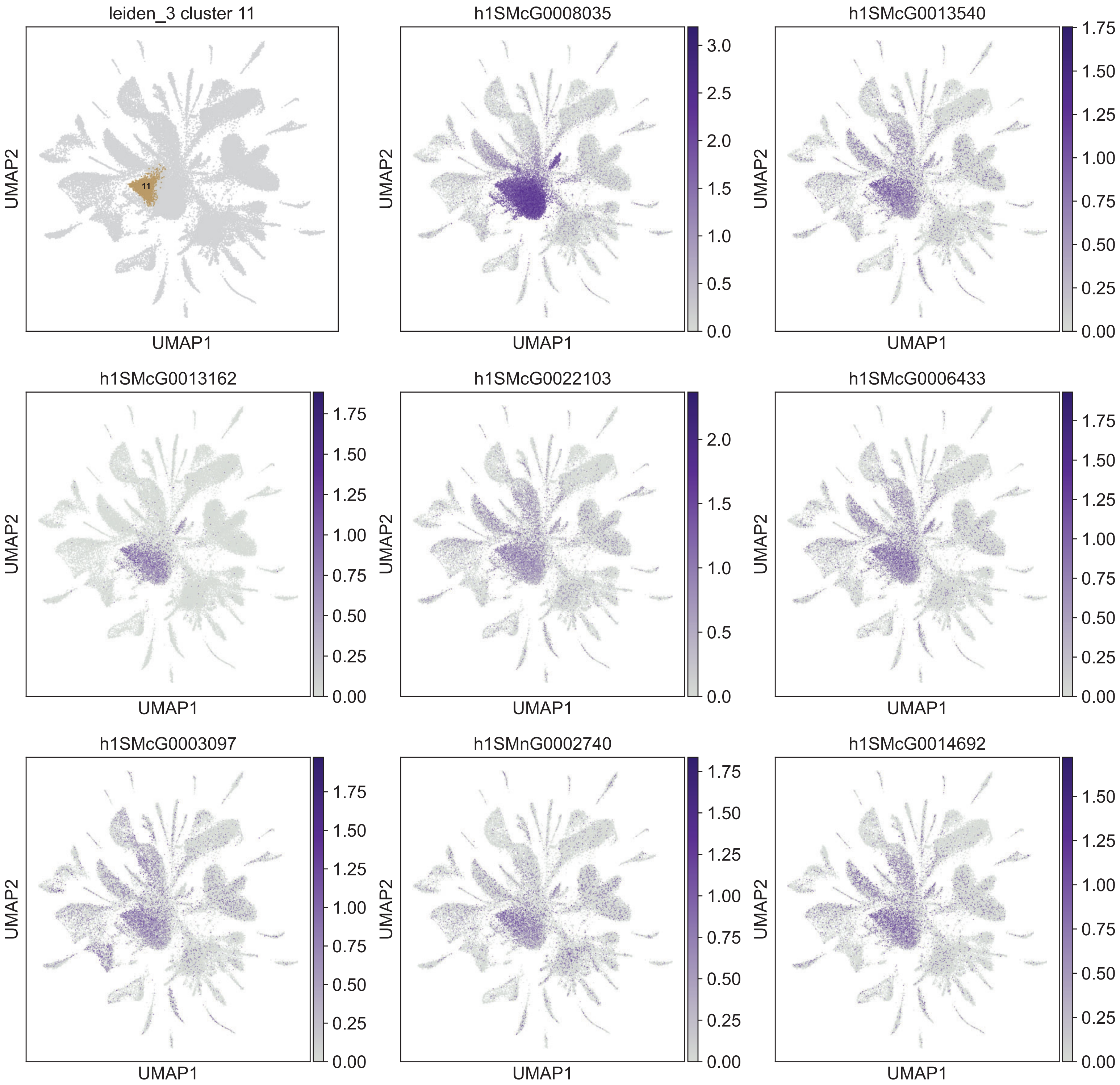

leiden_3 cluster 11
11
h1SMcG0008035
h1SMcG0013540
h1SMcG0013162
h1SMcG0022103
h1SMcG0006433
h1SMcG0003097
h1SMnG0002740
h1SMcG0014692
UMAP1
UMAP2

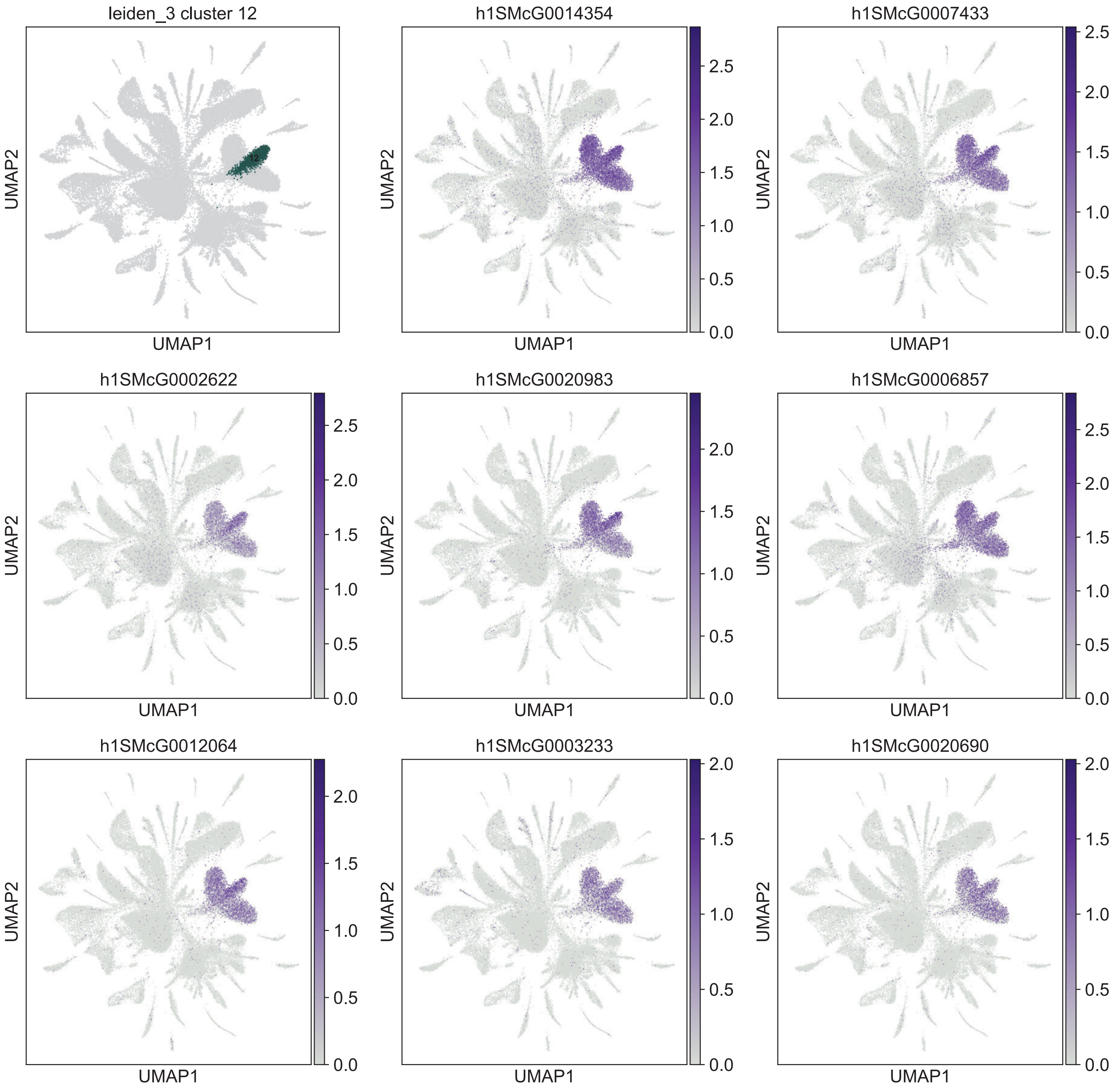

leiden_3 cluster 12
h1SMcG0014354
h1SMcG0007433
h1SMcG0002622
h1SMcG0020983
h1SMcG0006857
h1SMcG0012064
h1SMcG0003233
h1SMcG0020690
UMAP1
UMAP2

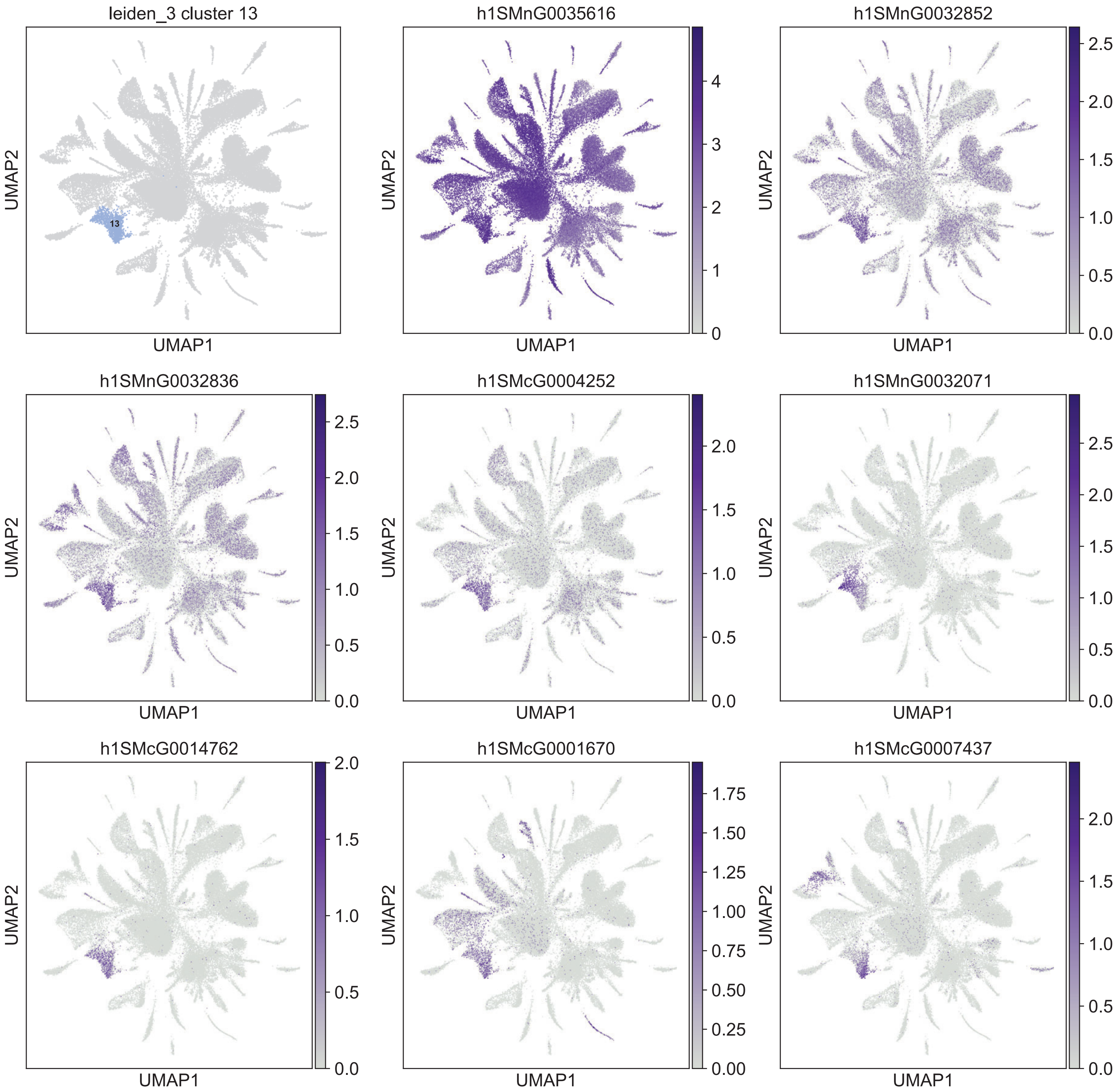
leiden_3 cluster 13
13
h1SMnG0035616
h1SMnG0032852
h1SMnG0032836
h1SMcG0004252
h1SMnG0032071
h1SMcG0014762
h1SMcG0001670
h1SMcG0007437
UMAP1
UMAP2

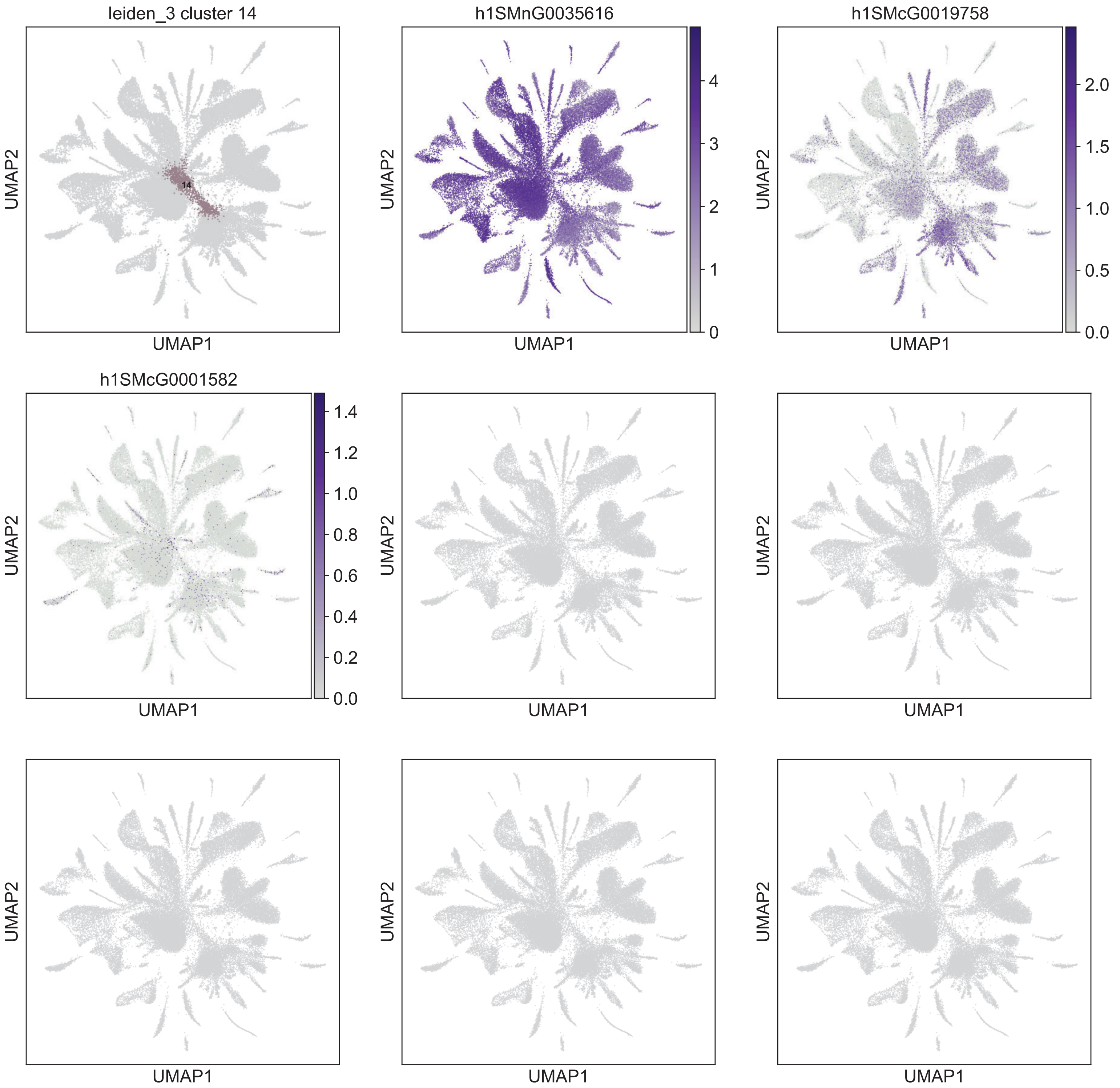
leiden_3 cluster 14
14
UMAP2
UMAP1
h1SMnG0035616
0
1
2
3
4
UMAP2
UMAP1
h1SMcG0019758
0.0
0.5
1.0
1.5
2.0
UMAP2
UMAP1
h1SMcG0001582
0.0
0.2
0.4
0.6
0.8
1.0
1.2
1.4
UMAP2
UMAP1
UMAP2
UMAP1
UMAP2
UMAP1
UMAP2
UMAP1
UMAP2
UMAP1
UMAP2
UMAP1

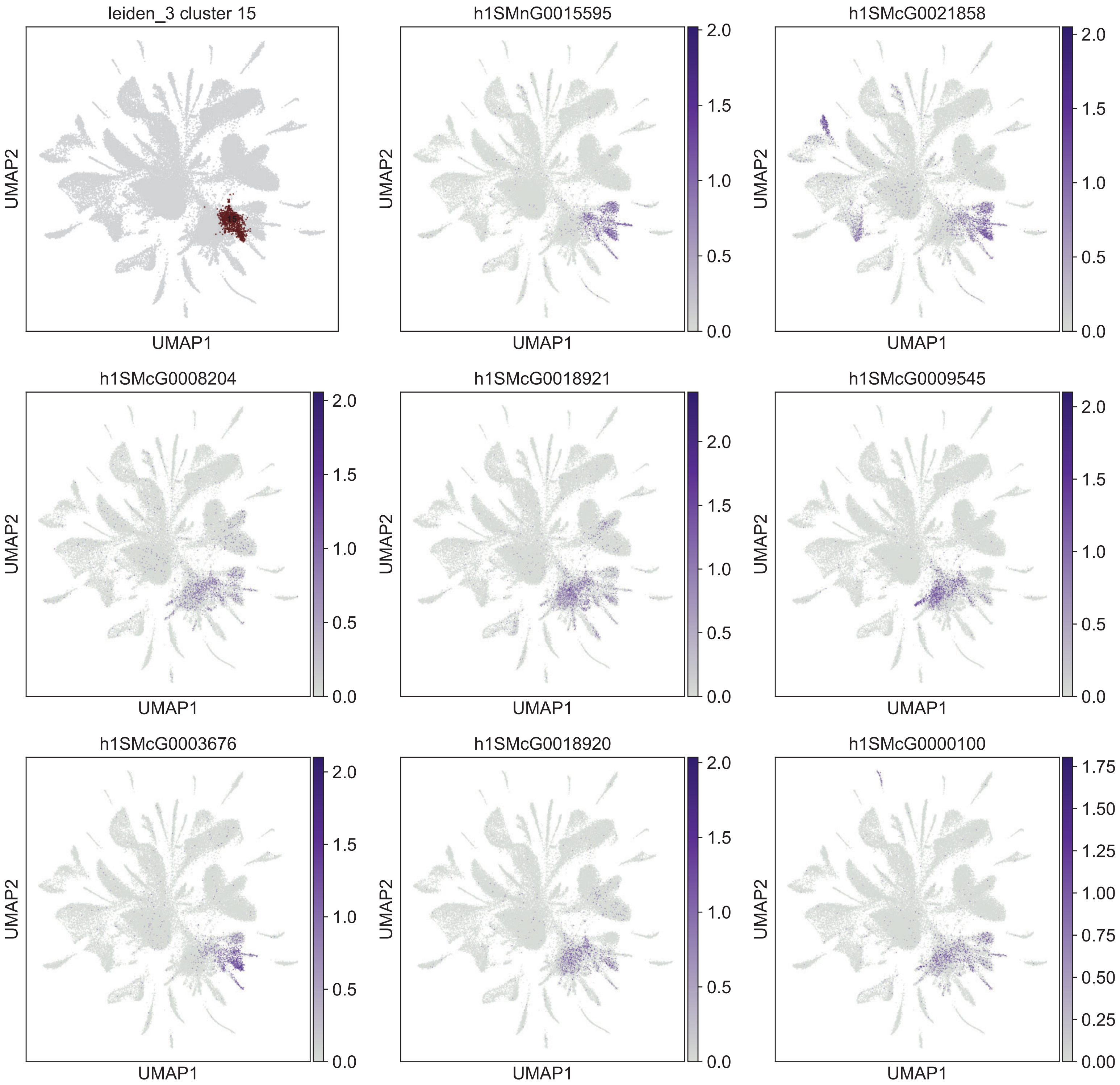
leiden_3 cluster 15
h1SMnG0015595
h1SMcG0021858
h1SMcG0008204
h1SMcG0018921
h1SMcG0009545
h1SMcG0003676
h1SMcG0018920
h1SMcG0000100
UMAP1
UMAP2

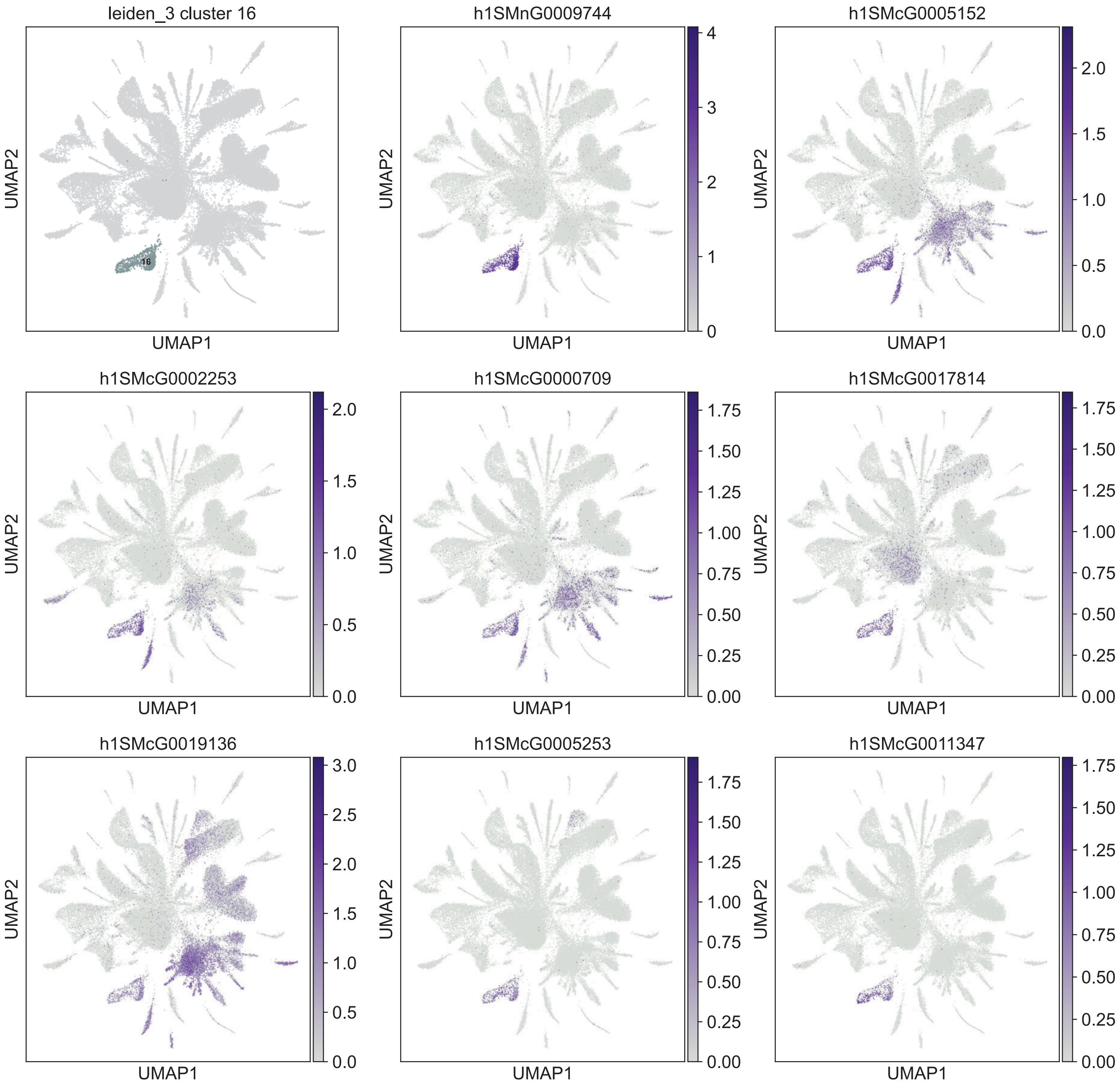

leiden_3 cluster 16
16
h1SMnG0009744
h1SMcG0005152
h1SMcG0002253
h1SMcG0000709
h1SMcG0017814
h1SMcG0019136
h1SMcG0005253
h1SMcG0011347
UMAP1
UMAP2

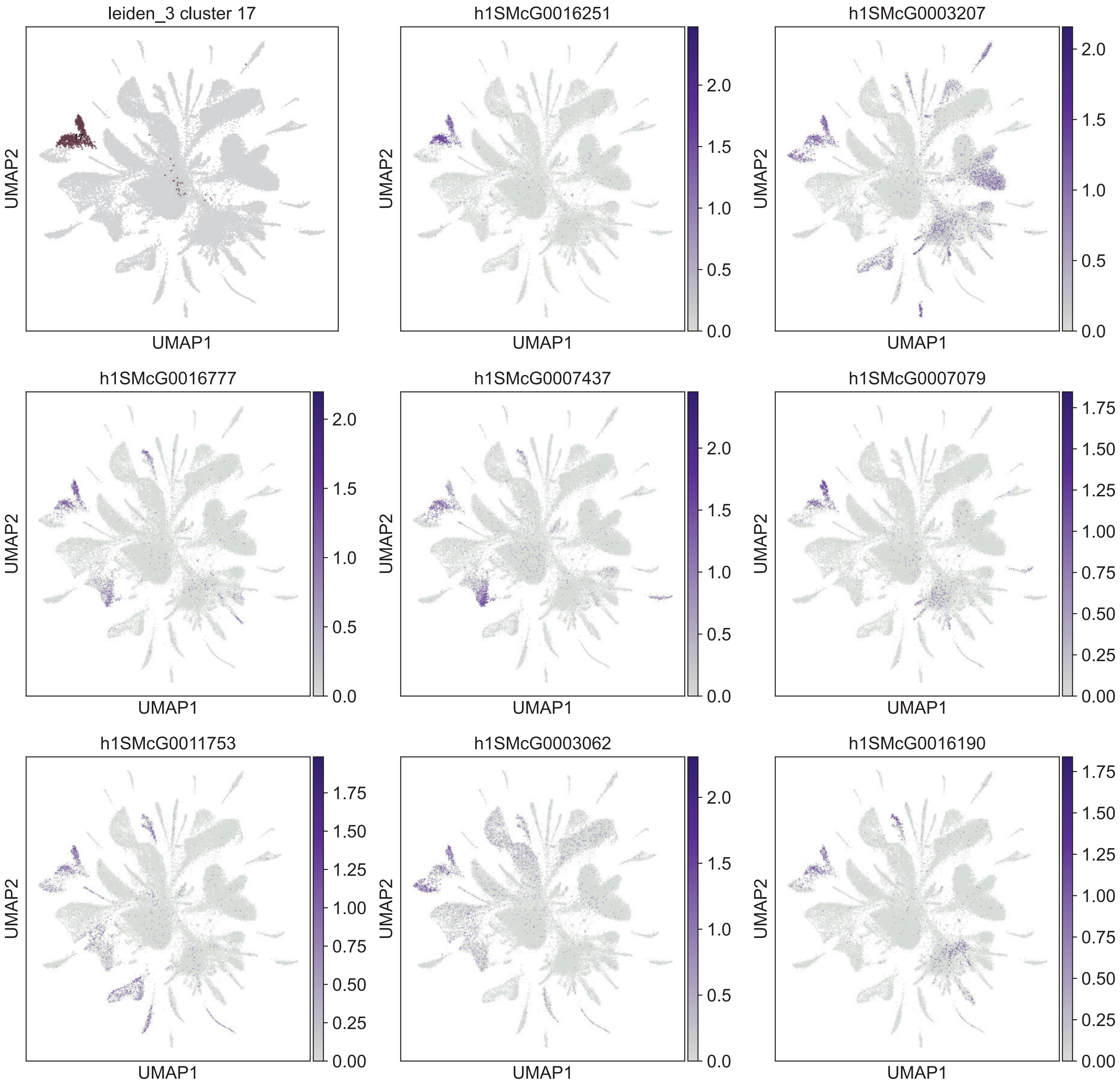
leiden_3 cluster 17
17
UMAP1
UMAP2
h1SMcG0016251
h1SMcG0003207
h1SMcG0016777
h1SMcG0007437
h1SMcG0007079
h1SMcG0011753
h1SMcG0003062
h1SMcG0016190
0.0
0.5
1.0
1.5
2.0
0.00
0.25
0.50
0.75
1.00
1.25
1.50
1.75

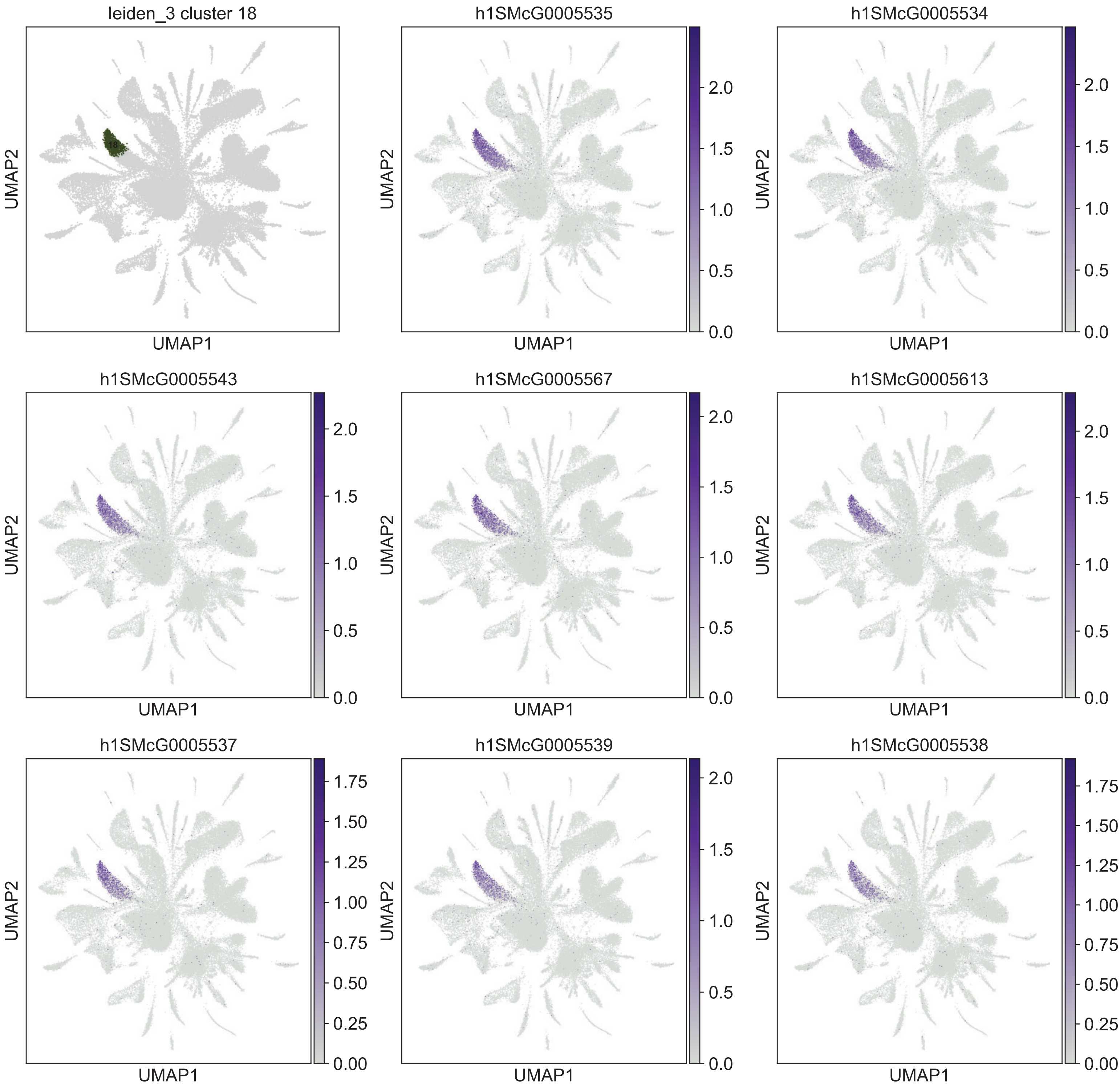
leiden_3 cluster 18
h1SMcG0005535
h1SMcG0005534
h1SMcG0005543
h1SMcG0005567
h1SMcG0005613
h1SMcG0005537
h1SMcG0005539
h1SMcG0005538
UMAP1
UMAP2

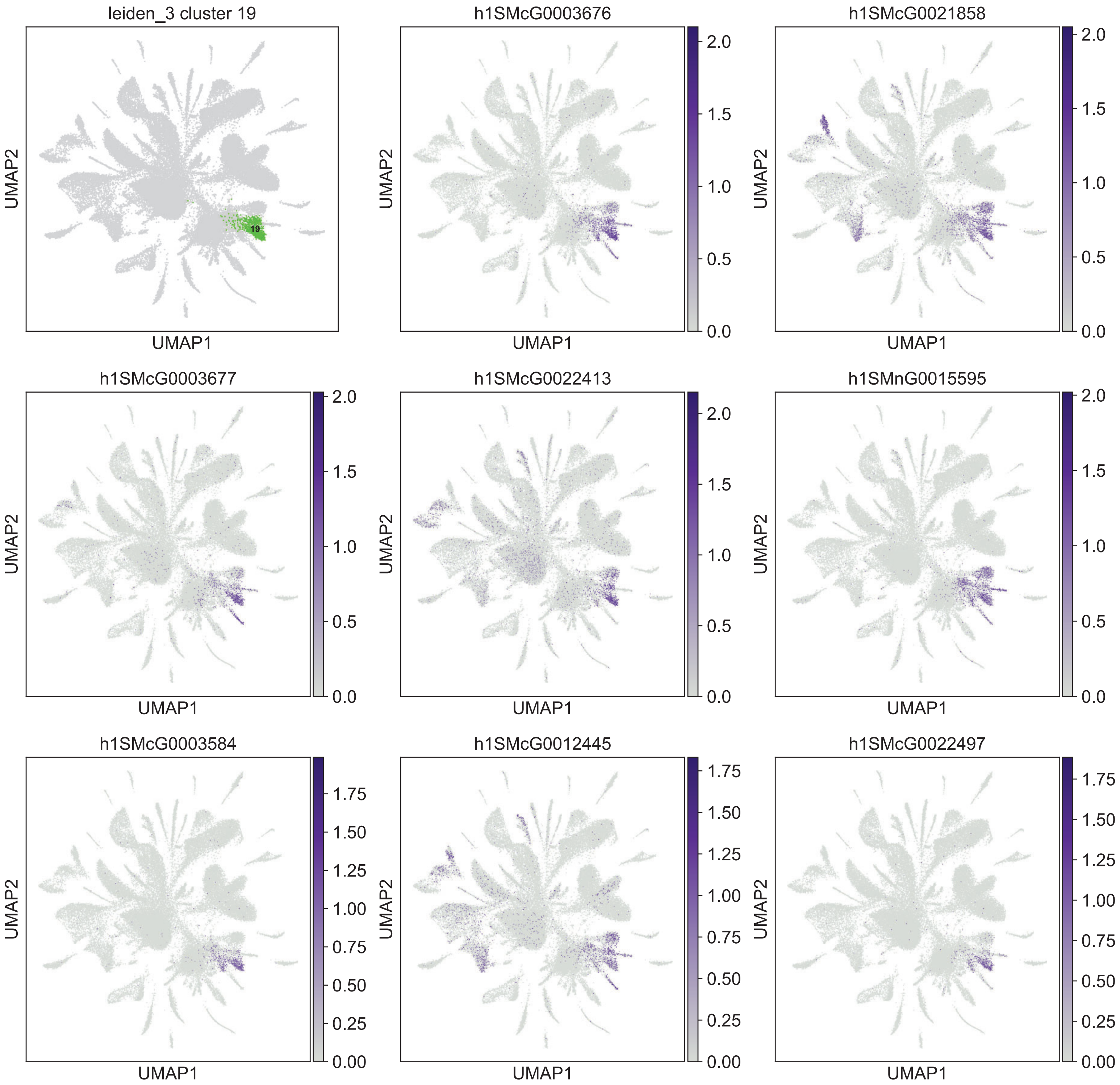
leiden_3 cluster 19
19
h1SMcG0003676
h1SMcG0021858
h1SMcG0003677
h1SMcG0022413
h1SMnG0015595
h1SMcG0003584
h1SMcG0012445
h1SMcG0022497
UMAP1
UMAP2
0.0
0.5
1.0
1.5
2.0
0.00
0.25
0.50
0.75
1.00
1.25
1.50
1.75

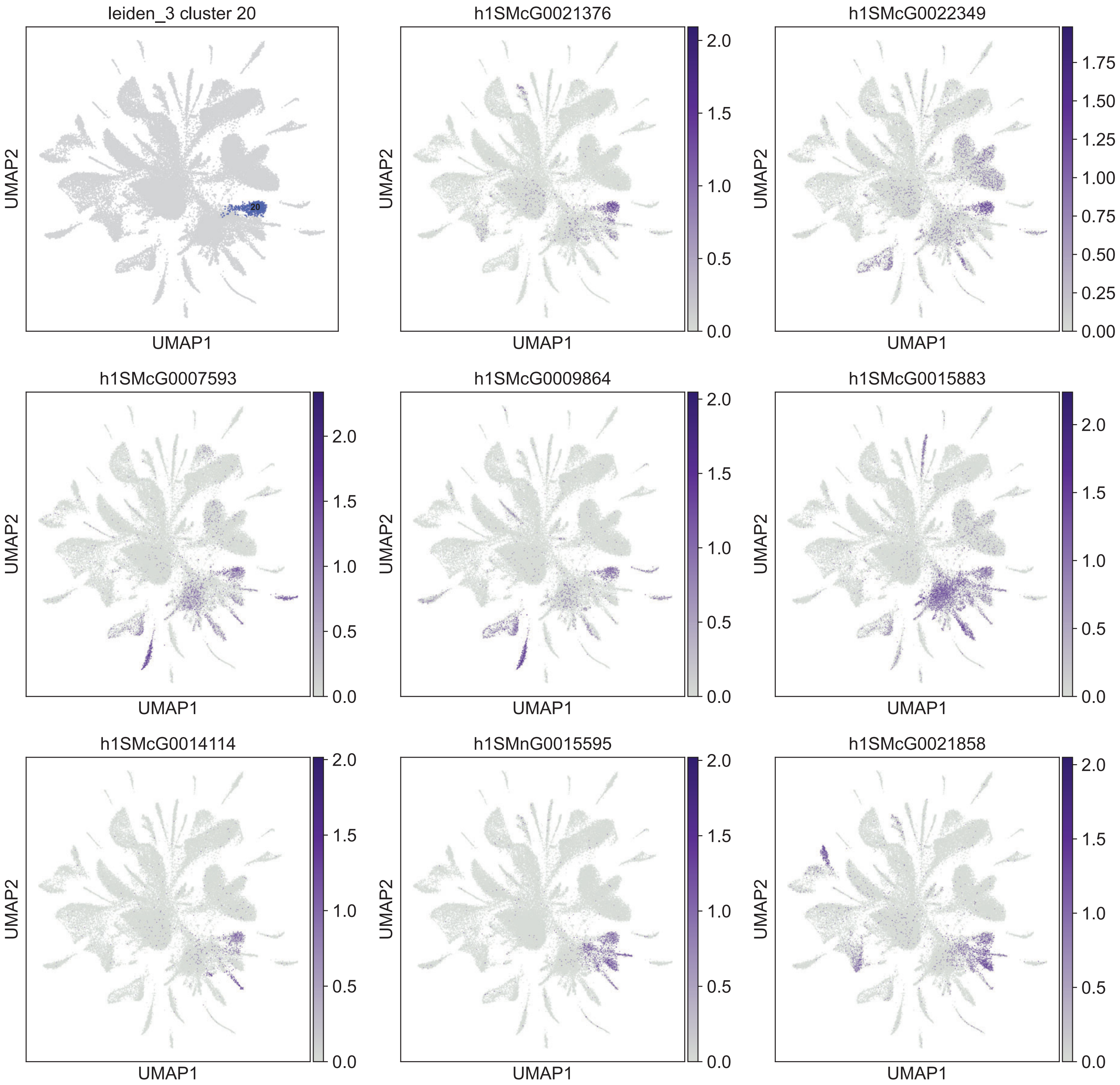
leiden_3 cluster 20
20
h1SMcG0021376
h1SMcG0022349
h1SMcG0007593
h1SMcG0009864
h1SMcG0015883
h1SMcG0014114
h1SMnG0015595
h1SMcG0021858
UMAP1
UMAP2
0.0
0.5
1.0
1.5
2.0
0.00
0.25
0.50
0.75
1.00
1.25
1.50
1.75

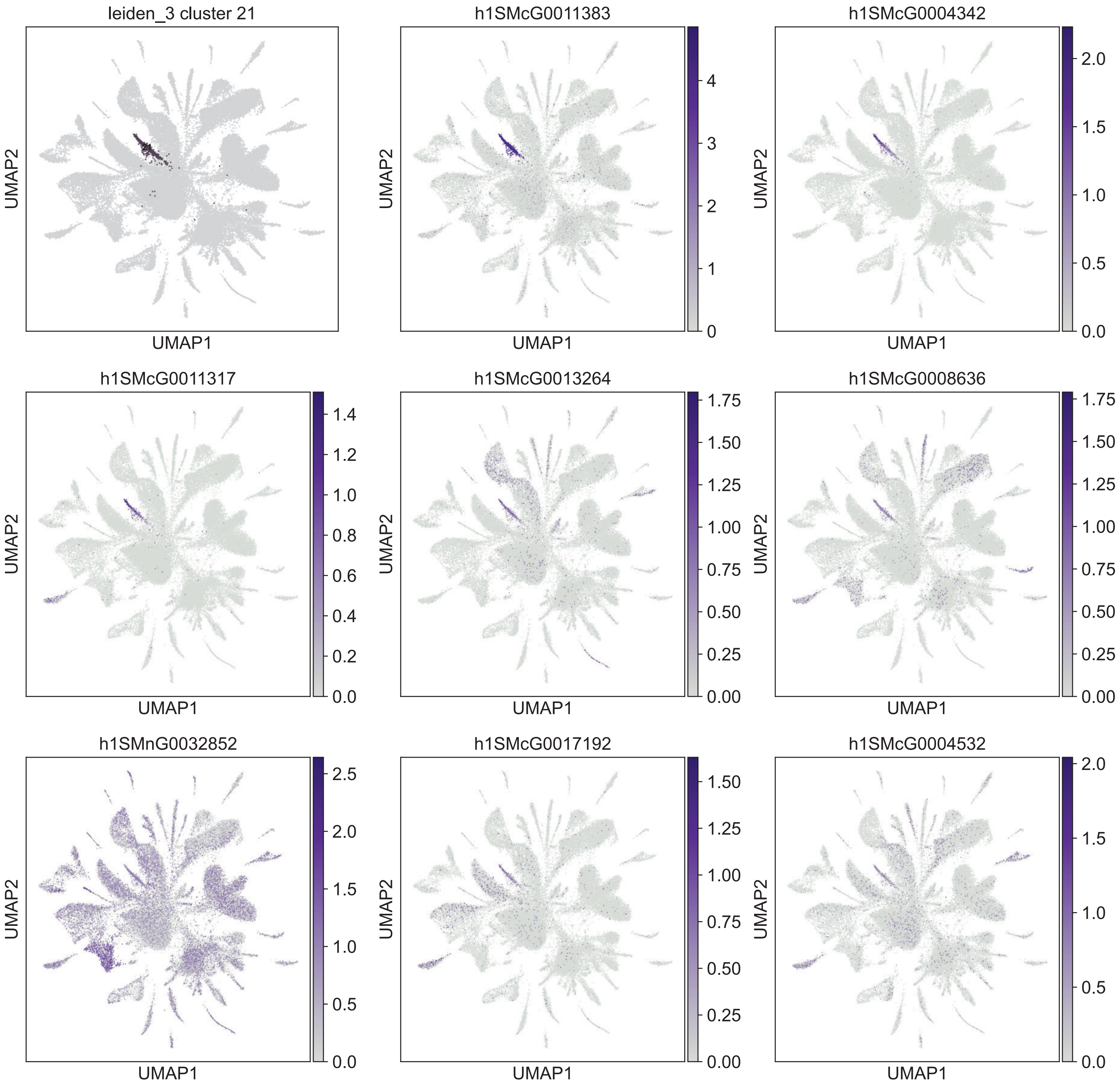

leiden_3 cluster 21
h1SMcG0011383
h1SMcG0004342
h1SMcG0011317
h1SMcG0013264
h1SMcG0008636
h1SMnG0032852
h1SMcG0017192
h1SMcG0004532
UMAP1
UMAP2

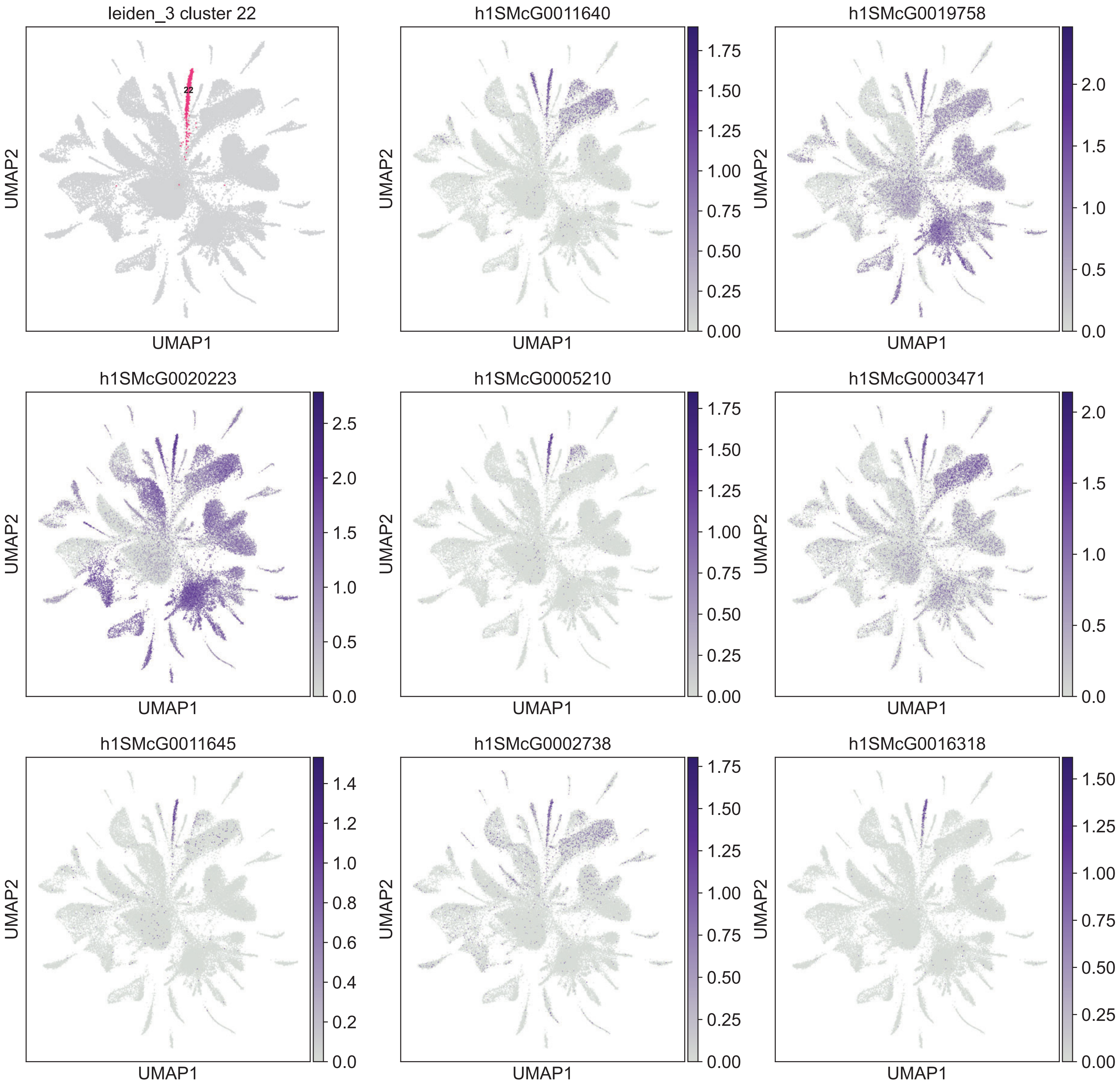

leiden_3 cluster 22
22
h1SMcG0011640
h1SMcG0019758
h1SMcG0020223
h1SMcG0005210
h1SMcG0003471
h1SMcG0011645
h1SMcG0002738
h1SMcG0016318
UMAP1
UMAP2

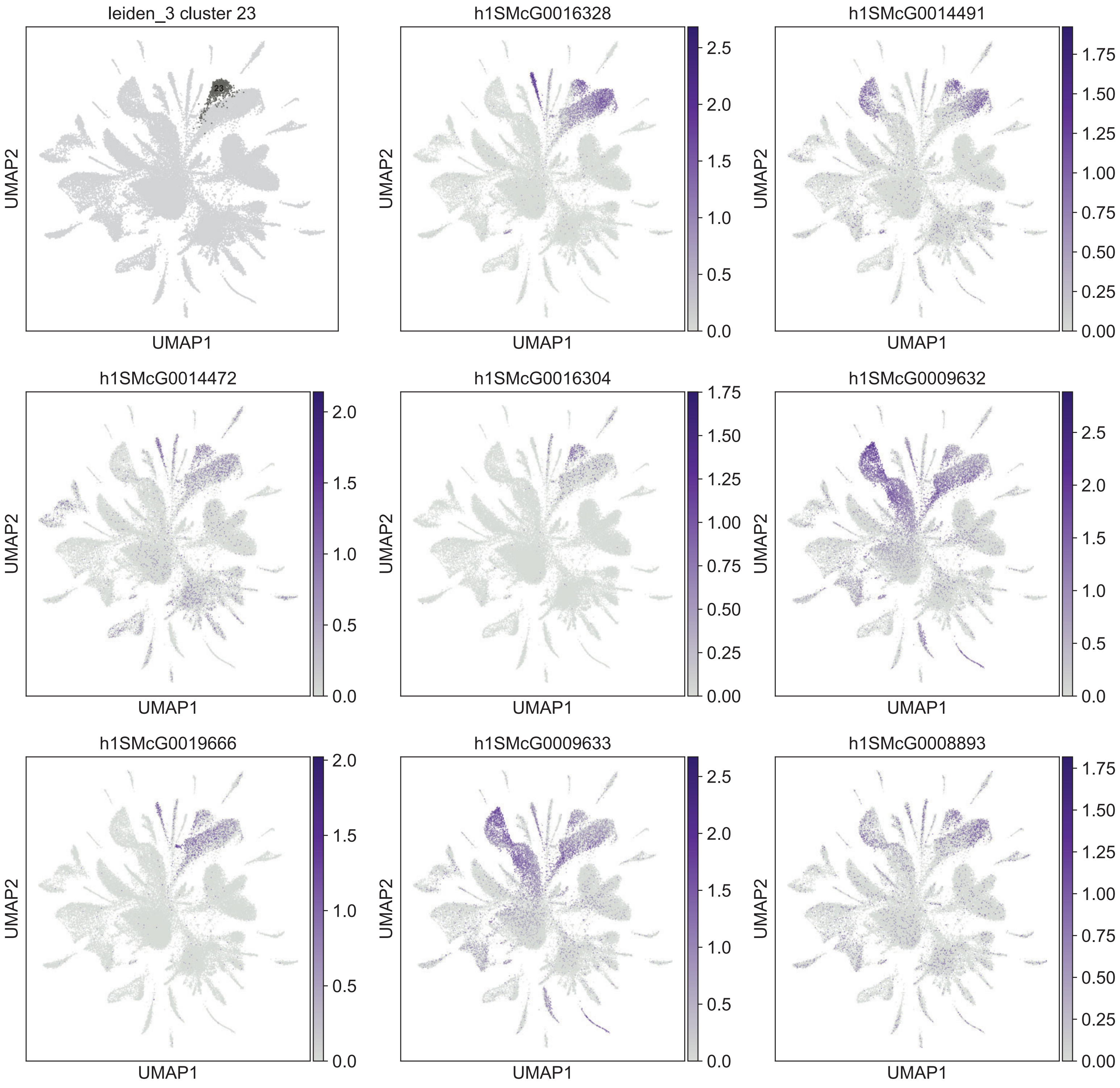

leiden_3 cluster 23
23
UMAP2
UMAP1
h1SMcG0016328
UMAP2
UMAP1
0.0
0.5
1.0
1.5
2.0
2.5
h1SMcG0014491
UMAP2
UMAP1
0.00
0.25
0.50
0.75
1.00
1.25
1.50
1.75
h1SMcG0014472
UMAP2
UMAP1
0.0
0.5
1.0
1.5
2.0
h1SMcG0016304
UMAP2
UMAP1
0.00
0.25
0.50
0.75
1.00
1.25
1.50
1.75
h1SMcG0009632
UMAP2
UMAP1
0.0
0.5
1.0
1.5
2.0
2.5
h1SMcG0019666
UMAP2
UMAP1
0.0
0.5
1.0
1.5
2.0
h1SMcG0009633
UMAP2
UMAP1
0.0
0.5
1.0
1.5
2.0
2.5
h1SMcG0008893
UMAP2
UMAP1
0.00
0.25
0.50
0.75
1.00
1.25
1.50
1.75

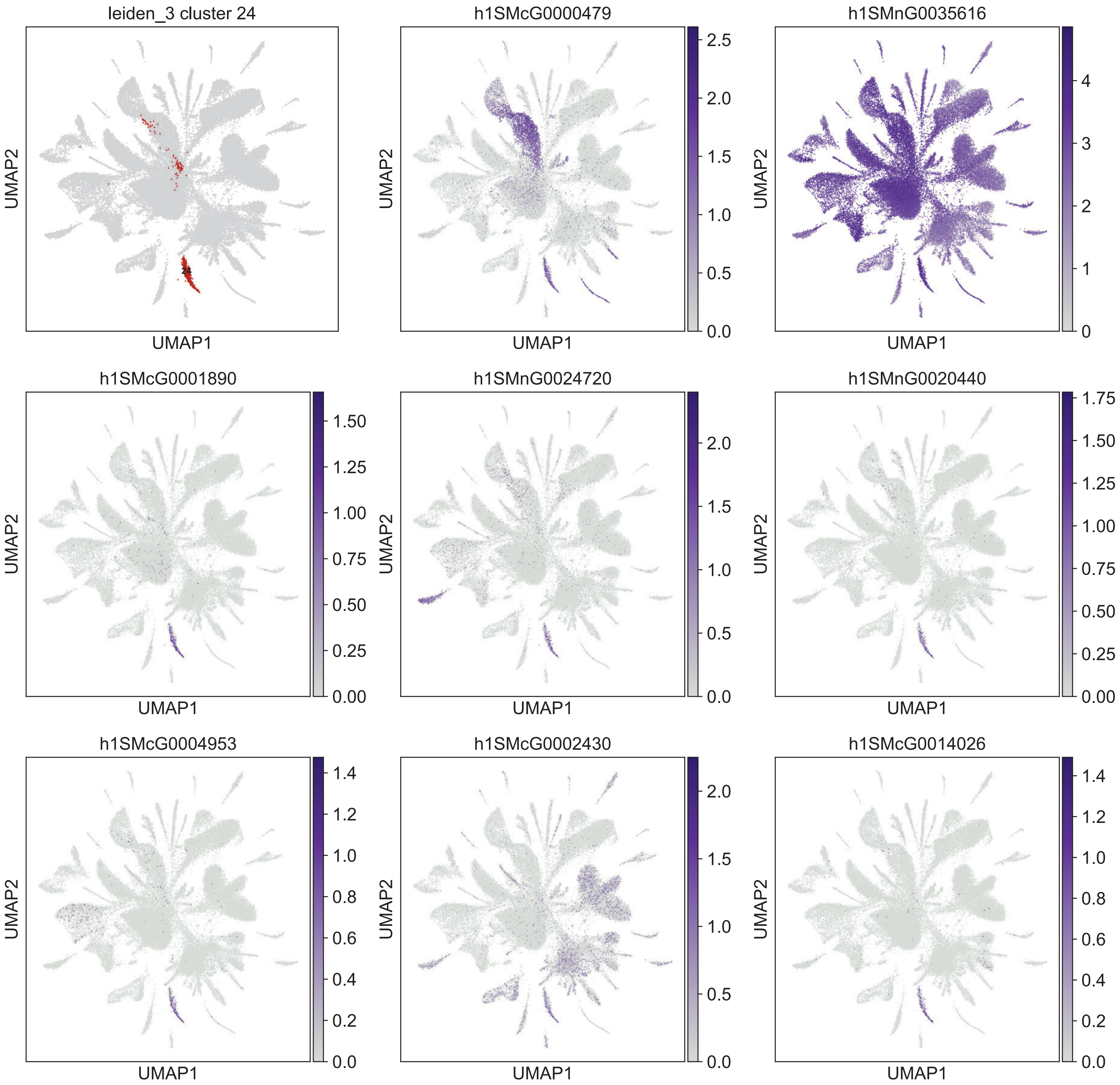

leiden_3 cluster 24
24
h1SMcG0000479
h1SMnG0035616
h1SMcG0001890
h1SMnG0024720
h1SMnG0020440
h1SMcG0004953
h1SMcG0002430
h1SMcG0014026
UMAP1
UMAP2

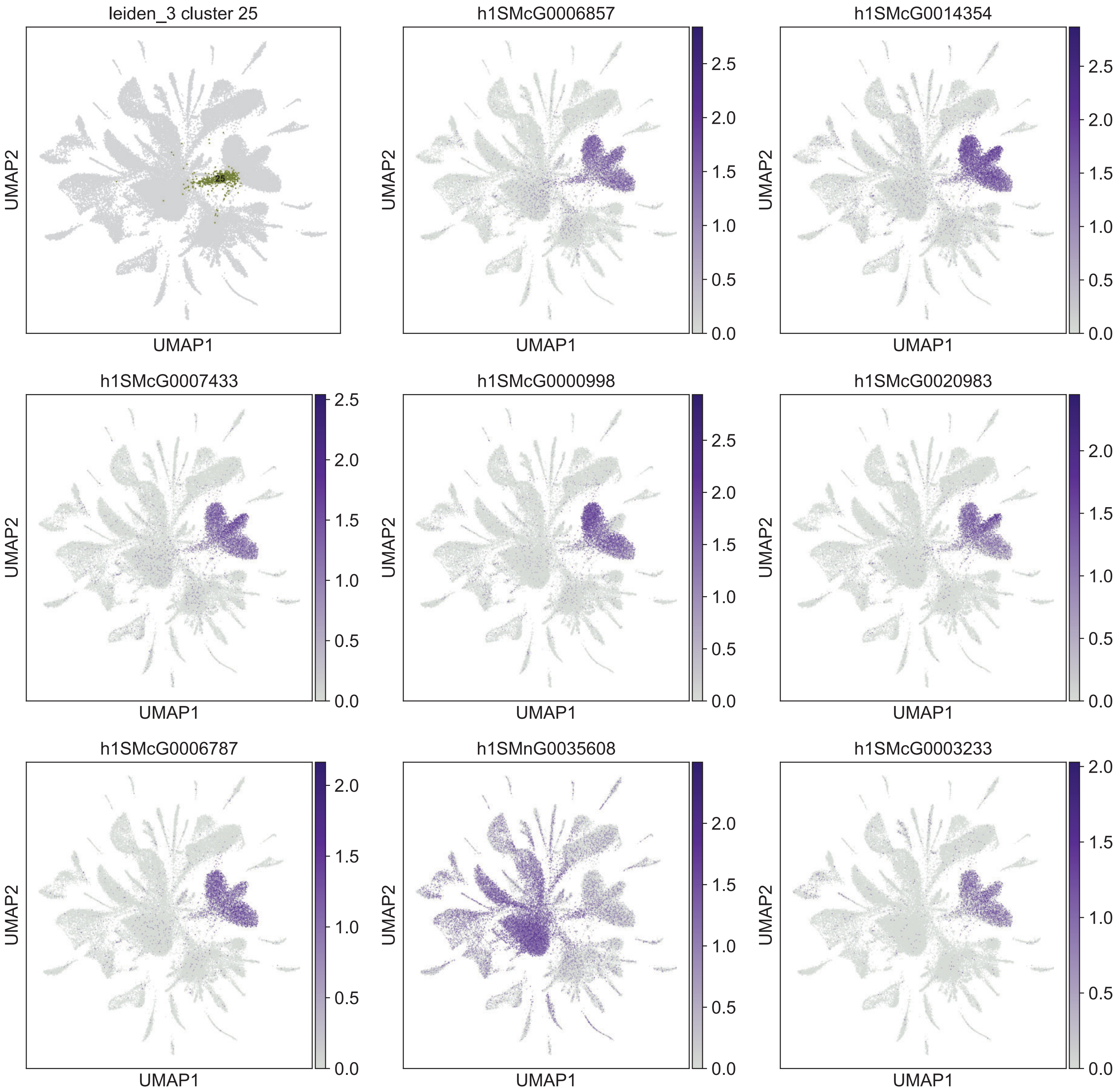
leiden_3 cluster 25
25
h1SMcG0006857
h1SMcG0014354
h1SMcG0007433
h1SMcG0000998
h1SMcG0020983
h1SMcG0006787
h1SMnG0035608
h1SMcG0003233
UMAP1
UMAP2

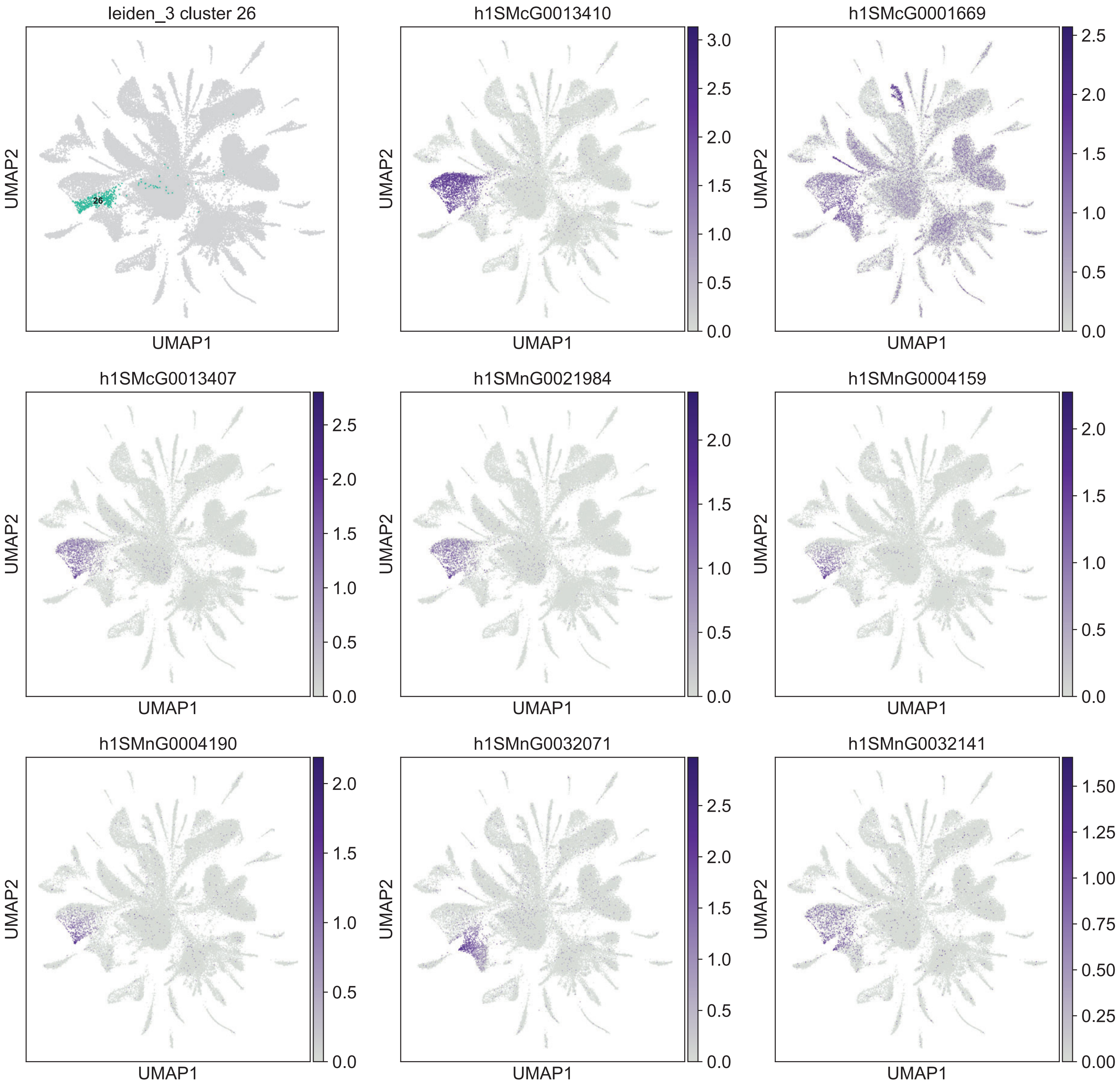
leiden_3 cluster 26
26
h1SMcG0013410
h1SMcG0001669
h1SMcG0013407
h1SMnG0021984
h1SMnG0004159
h1SMnG0004190
h1SMnG0032071
h1SMnG0032141
UMAP1
UMAP2

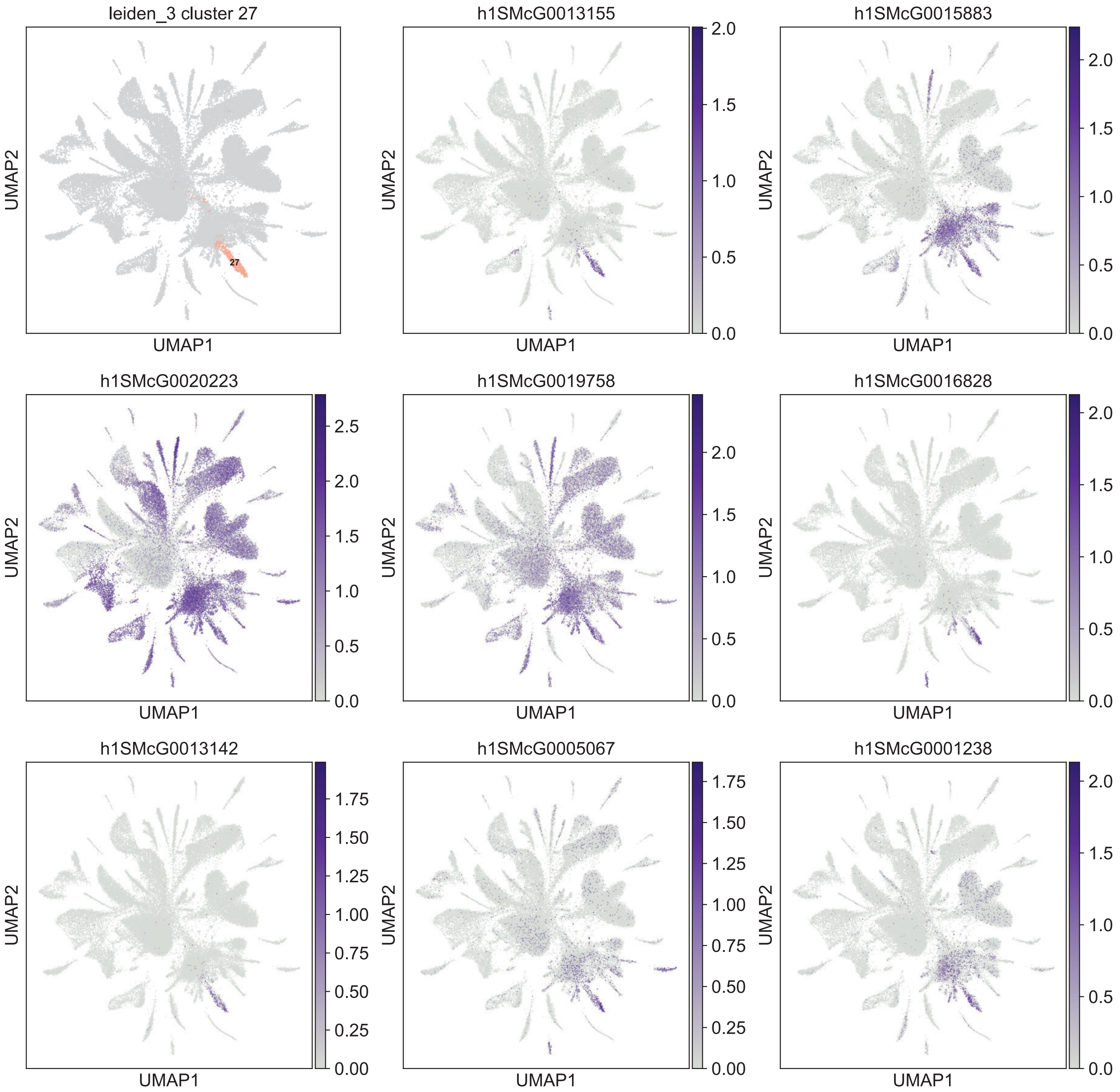

leiden_3 cluster 27
27
h1SMcG0013155
h1SMcG0015883
h1SMcG0020223
h1SMcG0019758
h1SMcG0016828
h1SMcG0013142
h1SMcG0005067
h1SMcG0001238
UMAP1
UMAP2

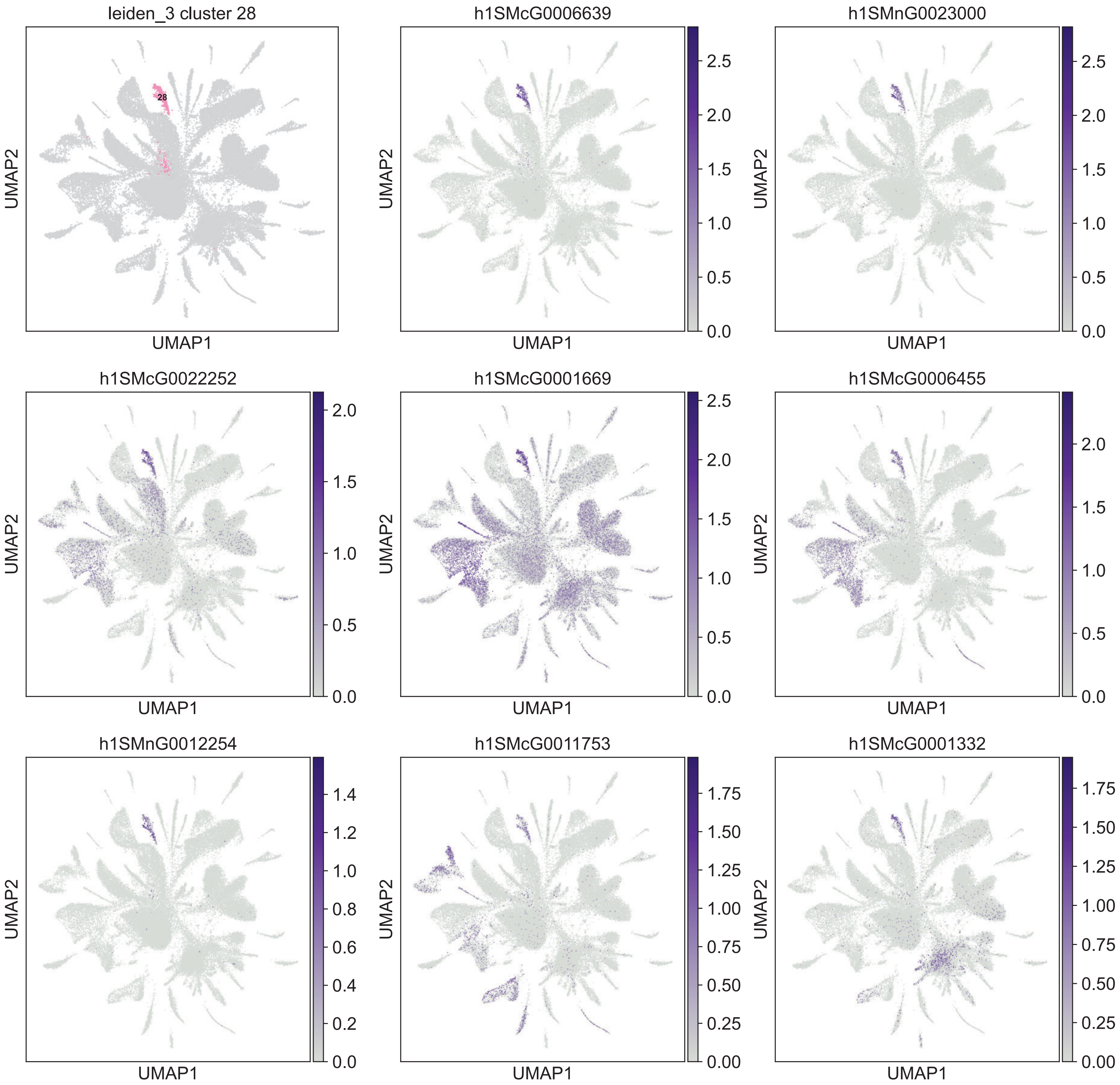

leiden_3 cluster 28
28
h1SMcG0006639
h1SMnG0023000
h1SMcG0022252
h1SMcG0001669
h1SMcG0006455
h1SMnG0012254
h1SMcG0011753
h1SMcG0001332
UMAP1
UMAP2

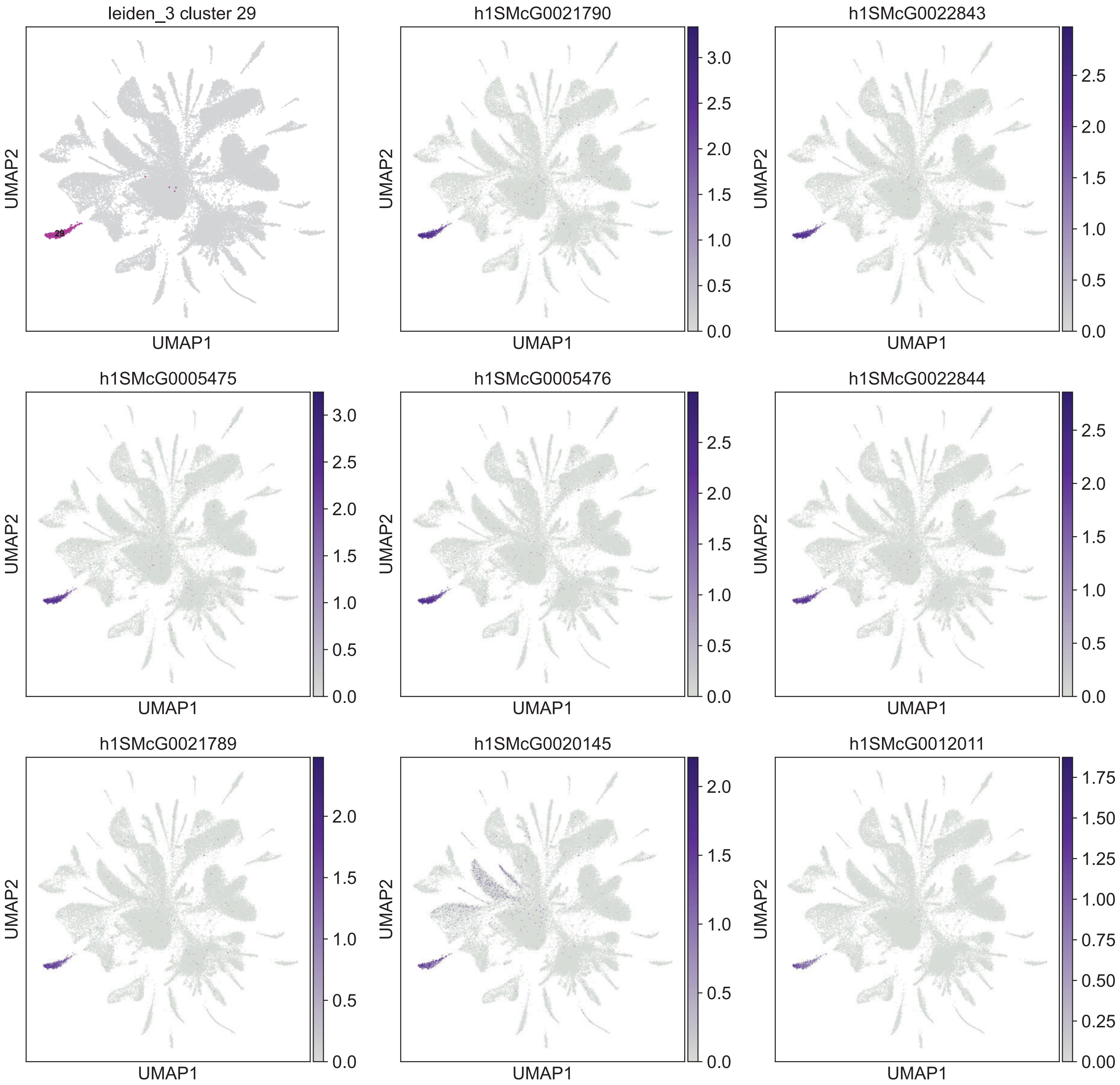

leiden_3 cluster 29
29
h1SMcG0021790
h1SMcG0022843
h1SMcG0005475
h1SMcG0005476
h1SMcG0022844
h1SMcG0021789
h1SMcG0020145
h1SMcG0012011
UMAP1
UMAP2
0.0
0.5
1.0
1.5
2.0
2.5
3.0
0.00
0.25
0.50
0.75
1.00
1.25
1.50
1.75

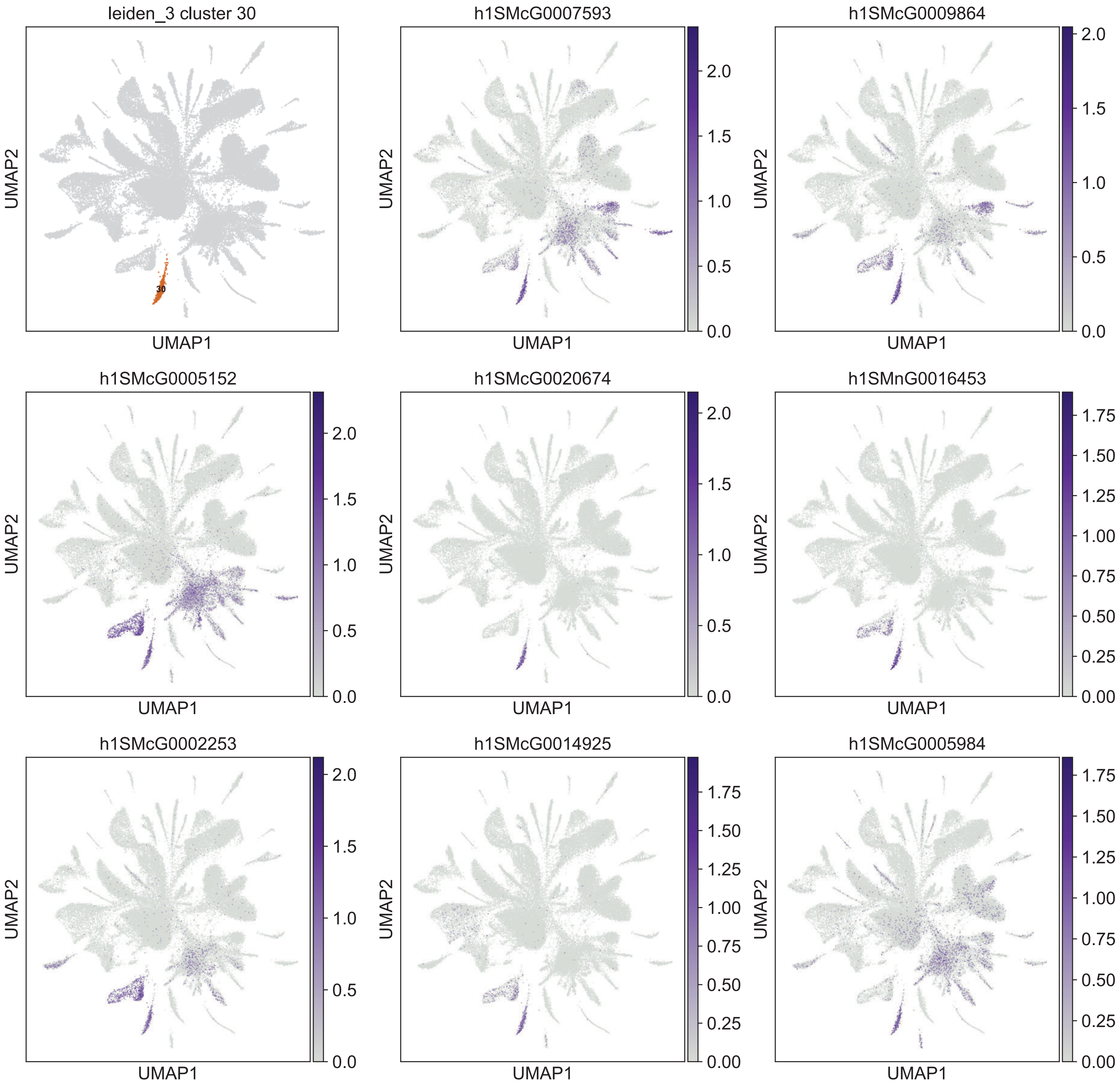
leiden_3 cluster 30
h1SMcG0007593
h1SMcG0009864
h1SMcG0005152
h1SMcG0020674
h1SMnG0016453
h1SMcG0002253
h1SMcG0014925
h1SMcG0005984
UMAP1
UMAP2
30

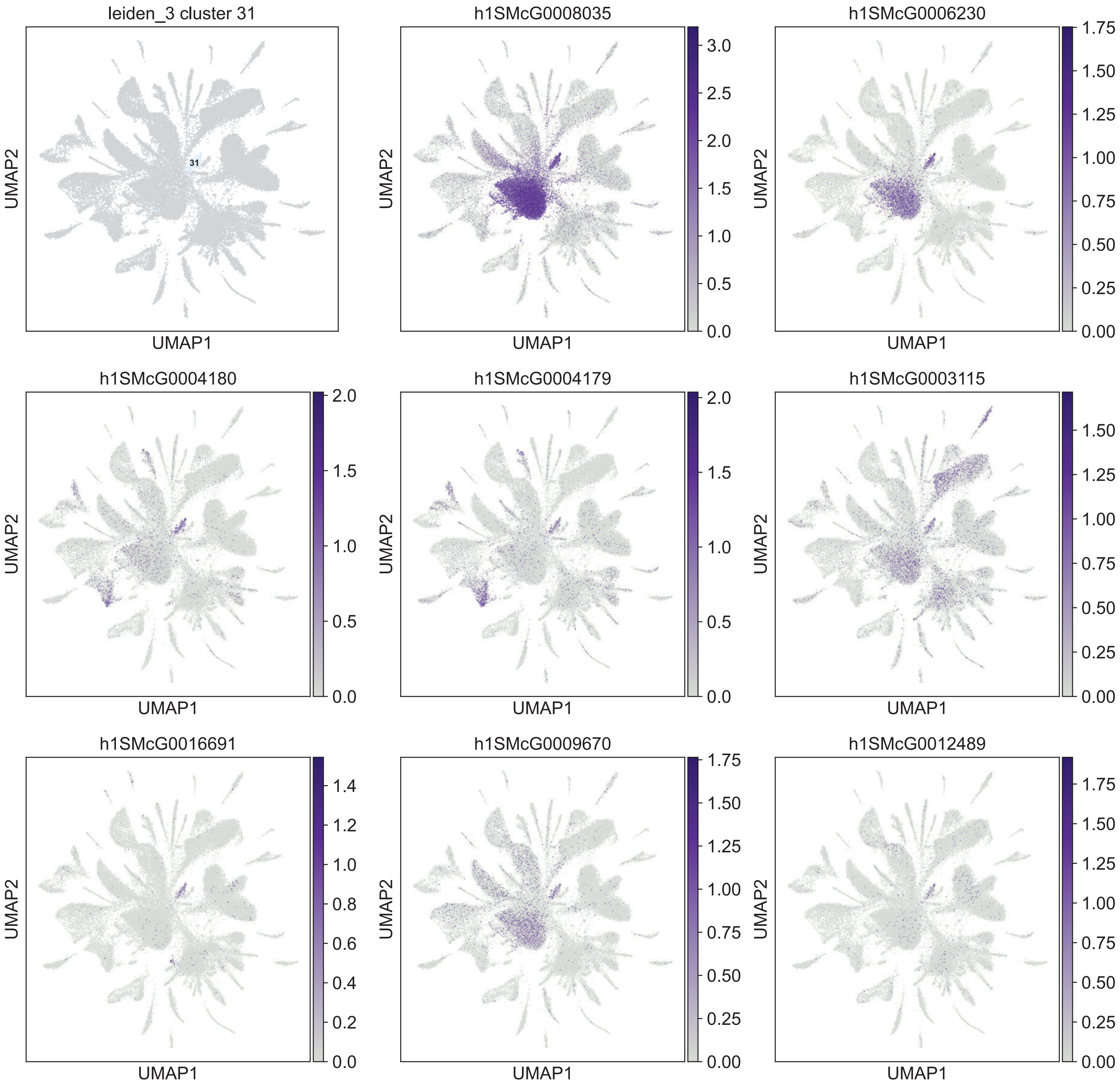

leiden_3 cluster 31
31
h1SMcG0008035
h1SMcG0006230
h1SMcG0004180
h1SMcG0004179
h1SMcG0003115
h1SMcG0016691
h1SMcG0009670
h1SMcG0012489
UMAP1
UMAP2

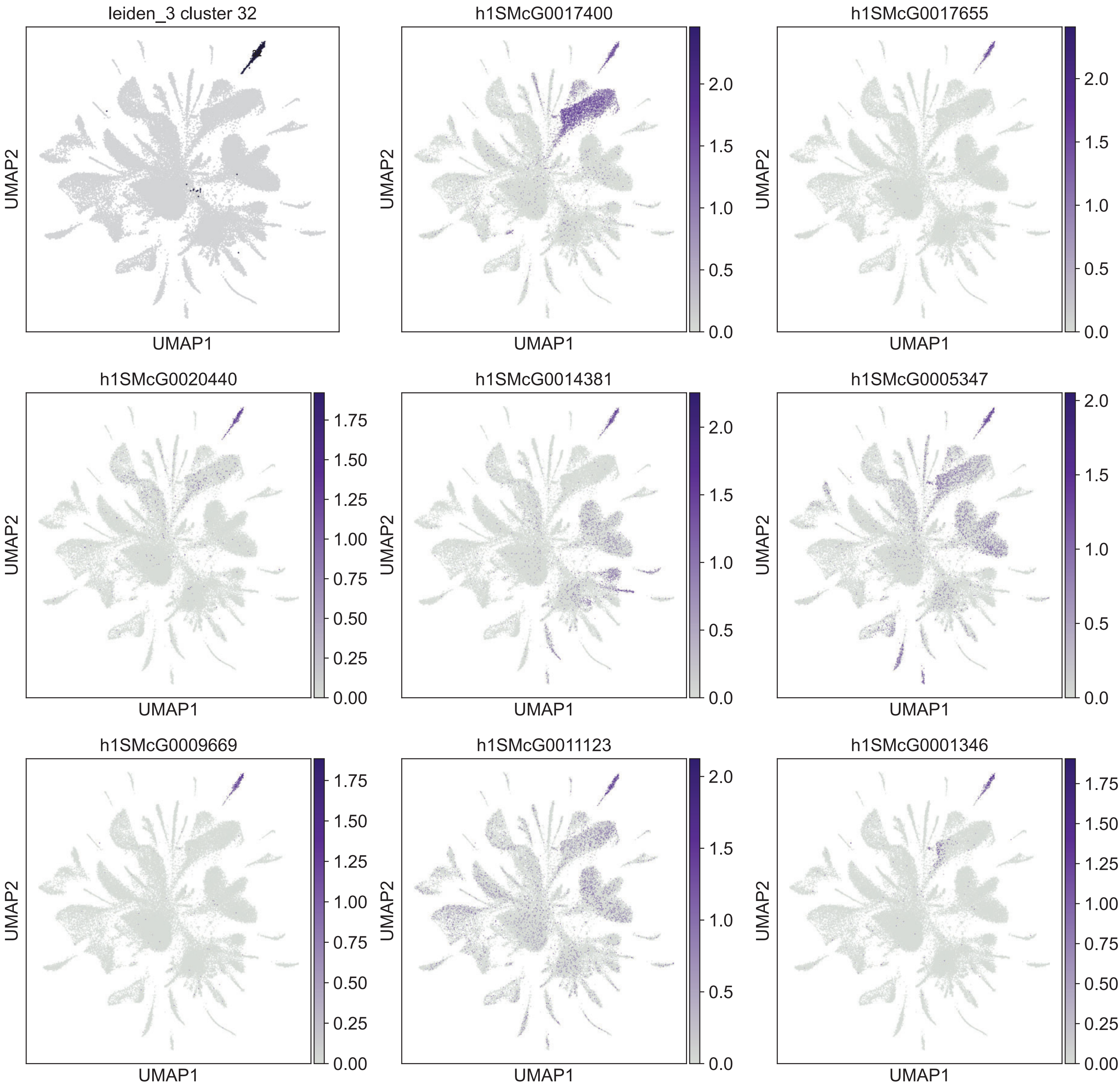

leiden_3 cluster 32
h1SMcG0017400
h1SMcG0017655
h1SMcG0020440
h1SMcG0014381
h1SMcG0005347
h1SMcG0009669
h1SMcG0011123
h1SMcG0001346
UMAP1
UMAP2

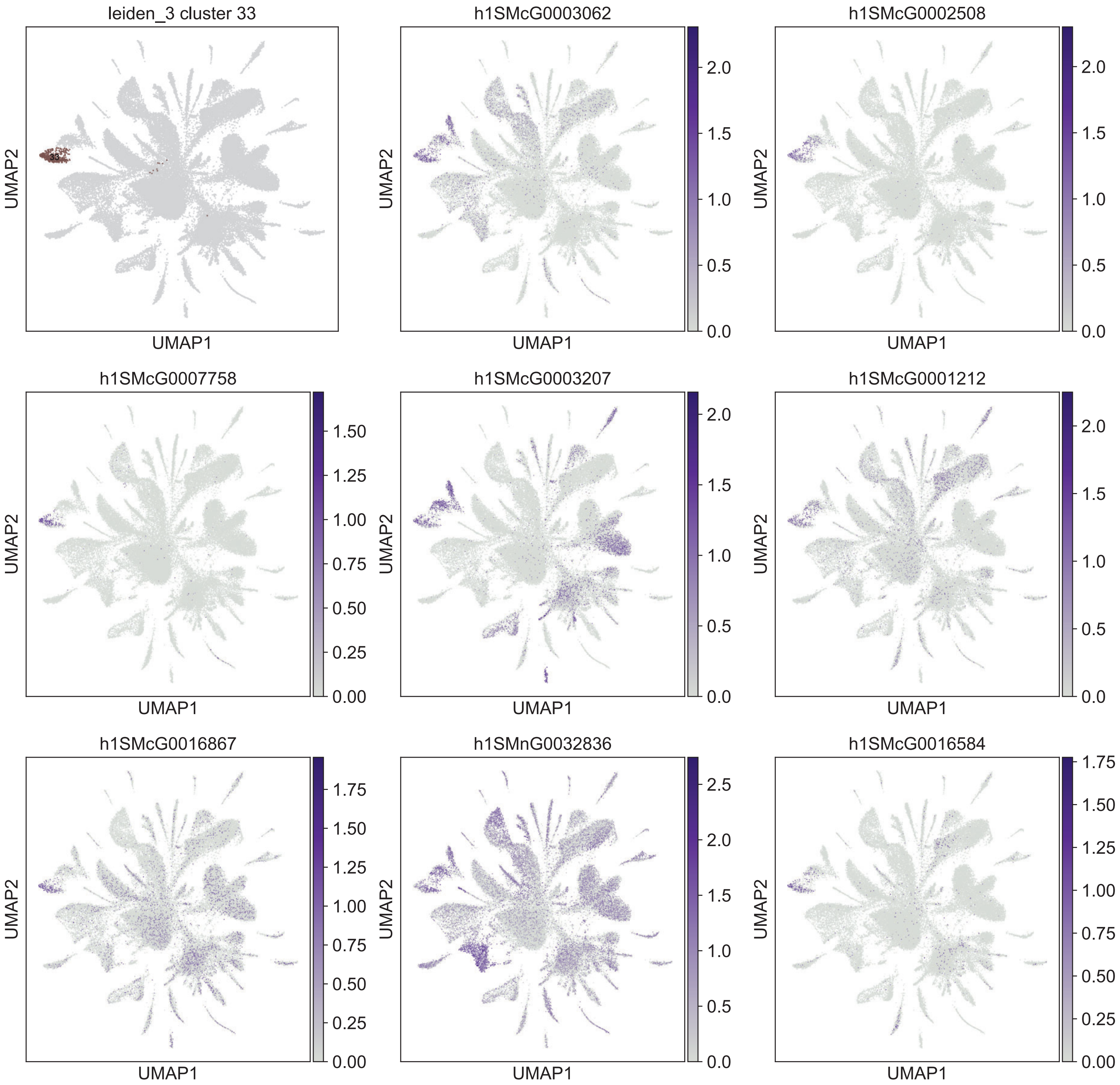
leiden_3 cluster 33
33
h1SMcG0003062
h1SMcG0002508
h1SMcG0007758
h1SMcG0003207
h1SMcG0001212
h1SMcG0016867
h1SMnG0032836
h1SMcG0016584
UMAP1
UMAP2
0.0
0.5
1.0
1.5
2.0
2.5
0.00
0.25
0.50
0.75
1.00
1.25
1.50
1.75

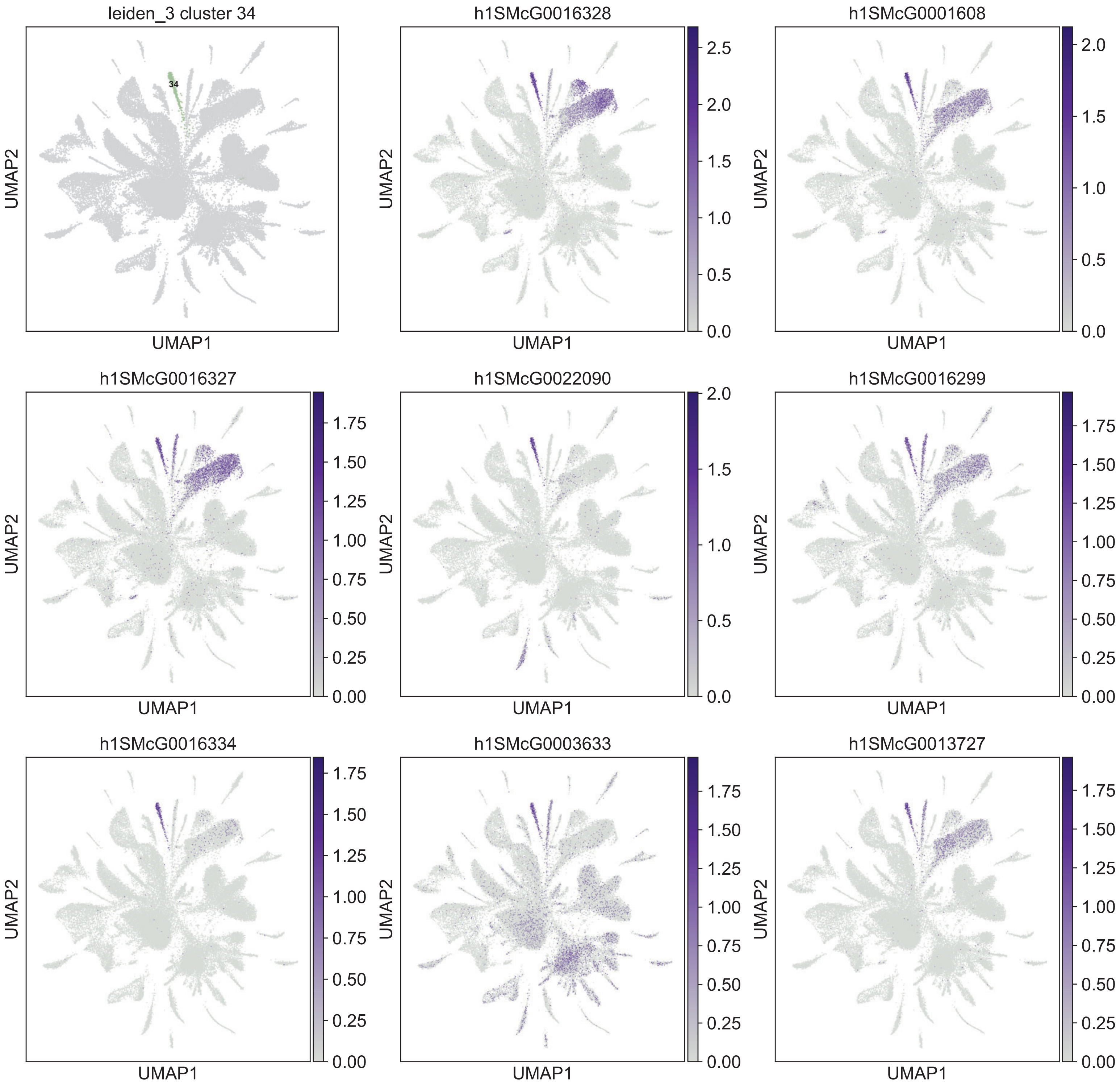

leiden_3 cluster 34
34
h1SMcG0016328
h1SMcG0001608
h1SMcG0016327
h1SMcG0022090
h1SMcG0016299
h1SMcG0016334
h1SMcG0003633
h1SMcG0013727
UMAP1
UMAP2

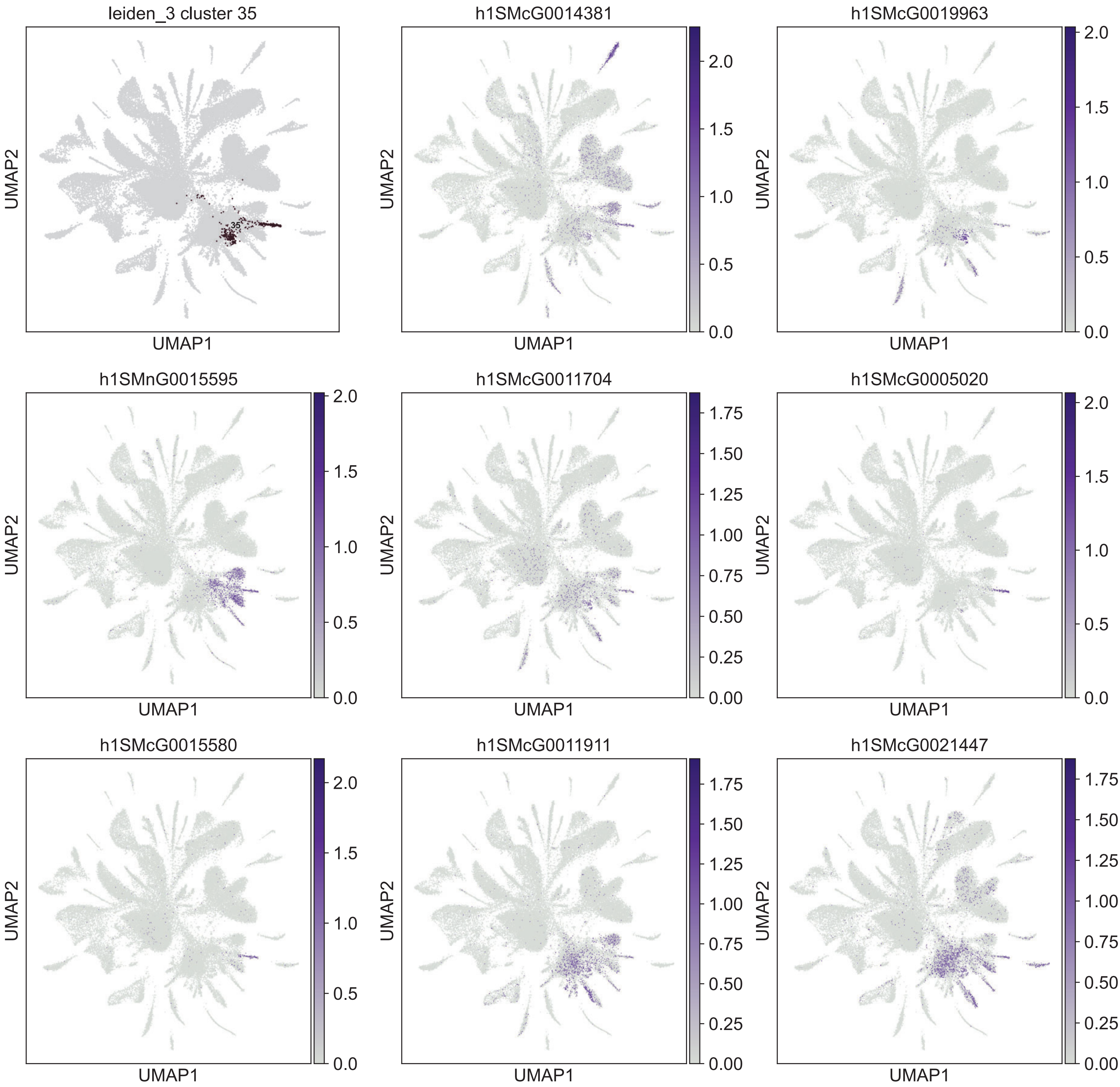
leiden_3 cluster 35
h1SMcG0014381
h1SMcG0019963
h1SMnG0015595
h1SMcG0011704
h1SMcG0005020
h1SMcG0015580
h1SMcG0011911
h1SMcG0021447
UMAP1
UMAP2
35

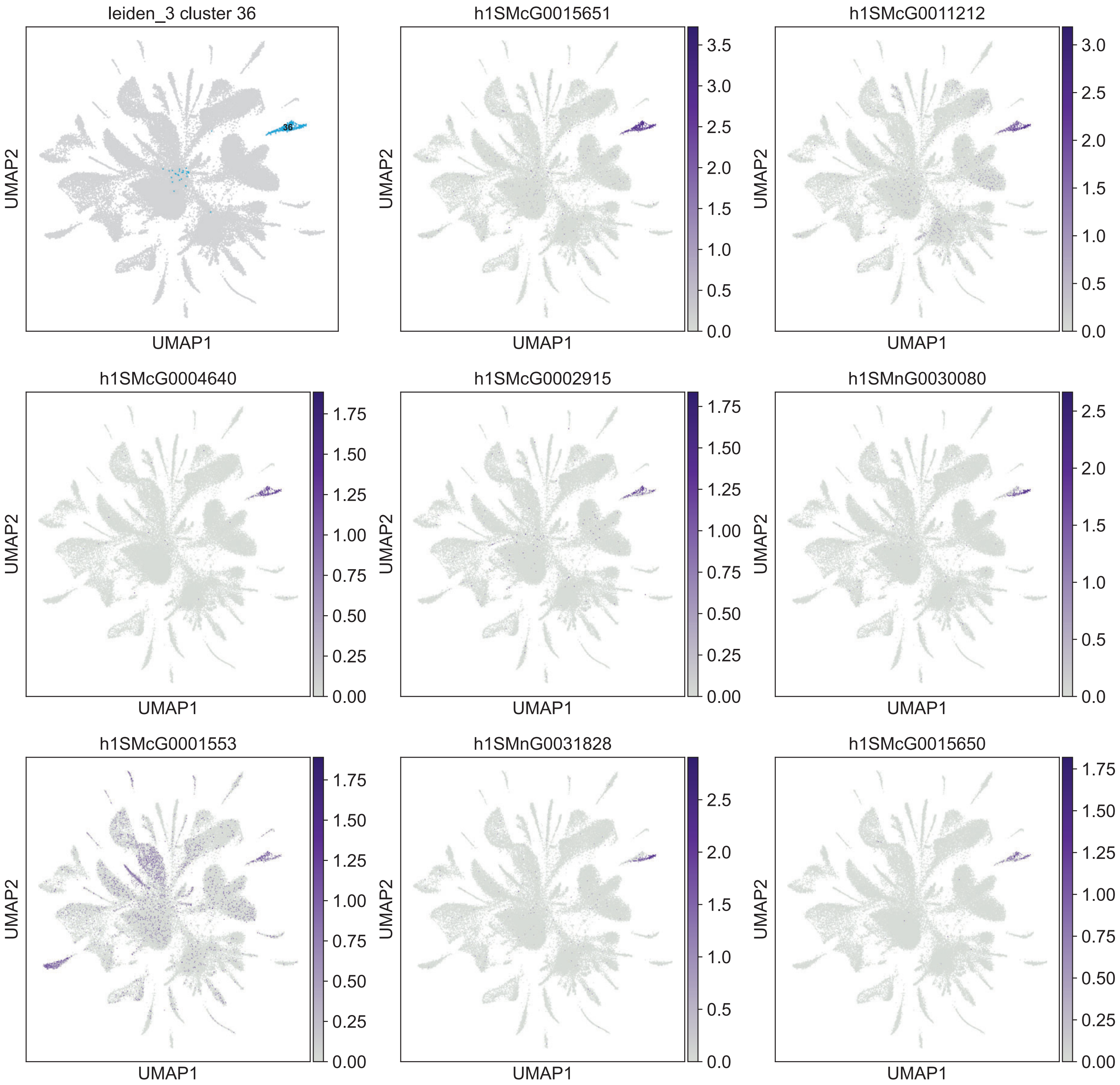

leiden_3 cluster 36
36
h1SMcG0015651
h1SMcG0011212
h1SMcG0004640
h1SMcG0002915
h1SMnG0030080
h1SMcG0001553
h1SMnG0031828
h1SMcG0015650
UMAP1
UMAP2

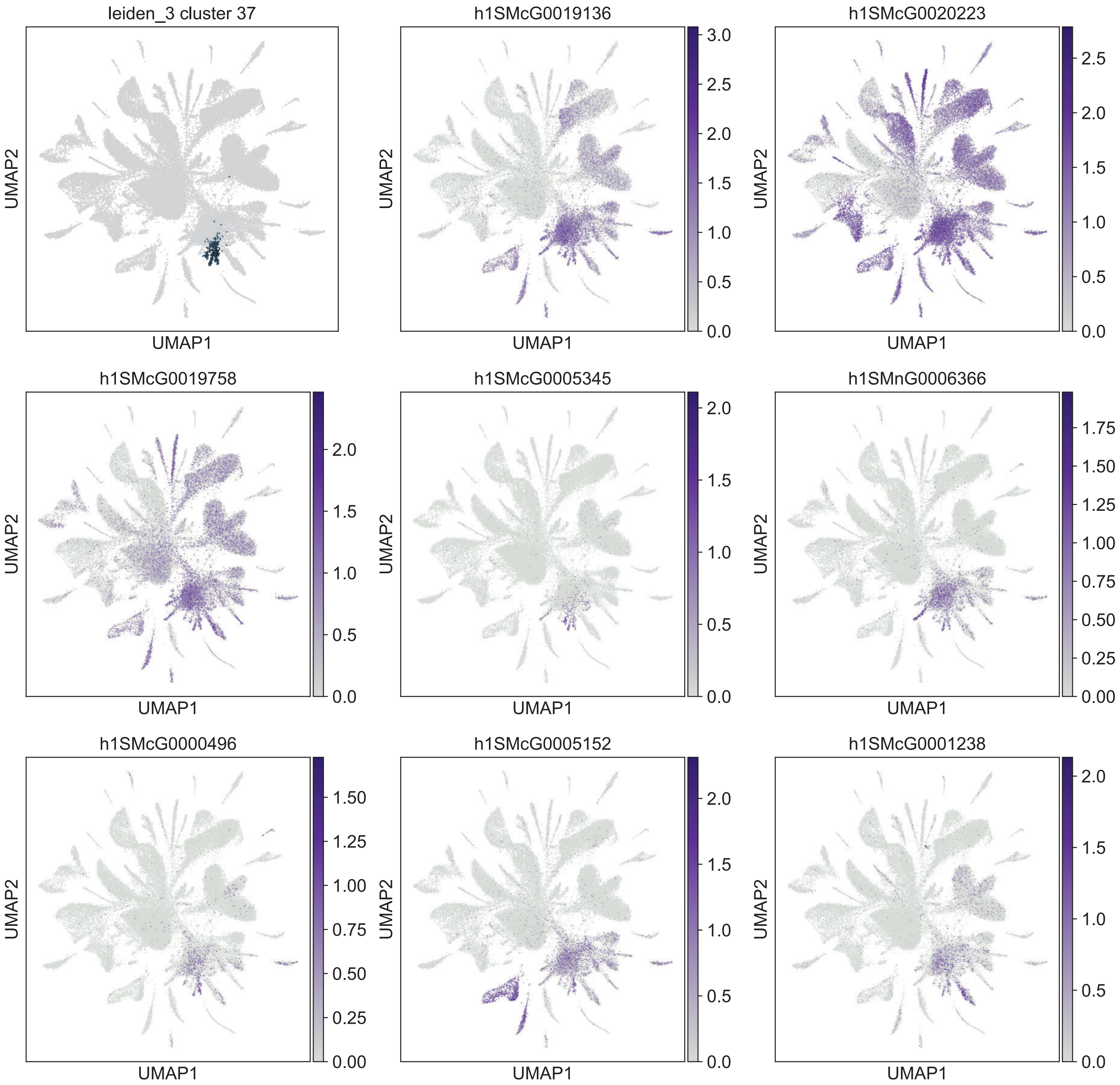
leiden_3 cluster 37
h1SMcG0019136
h1SMcG0020223
h1SMcG0019758
h1SMcG0005345
h1SMnG0006366
h1SMcG0000496
h1SMcG0005152
h1SMcG0001238
UMAP1
UMAP2

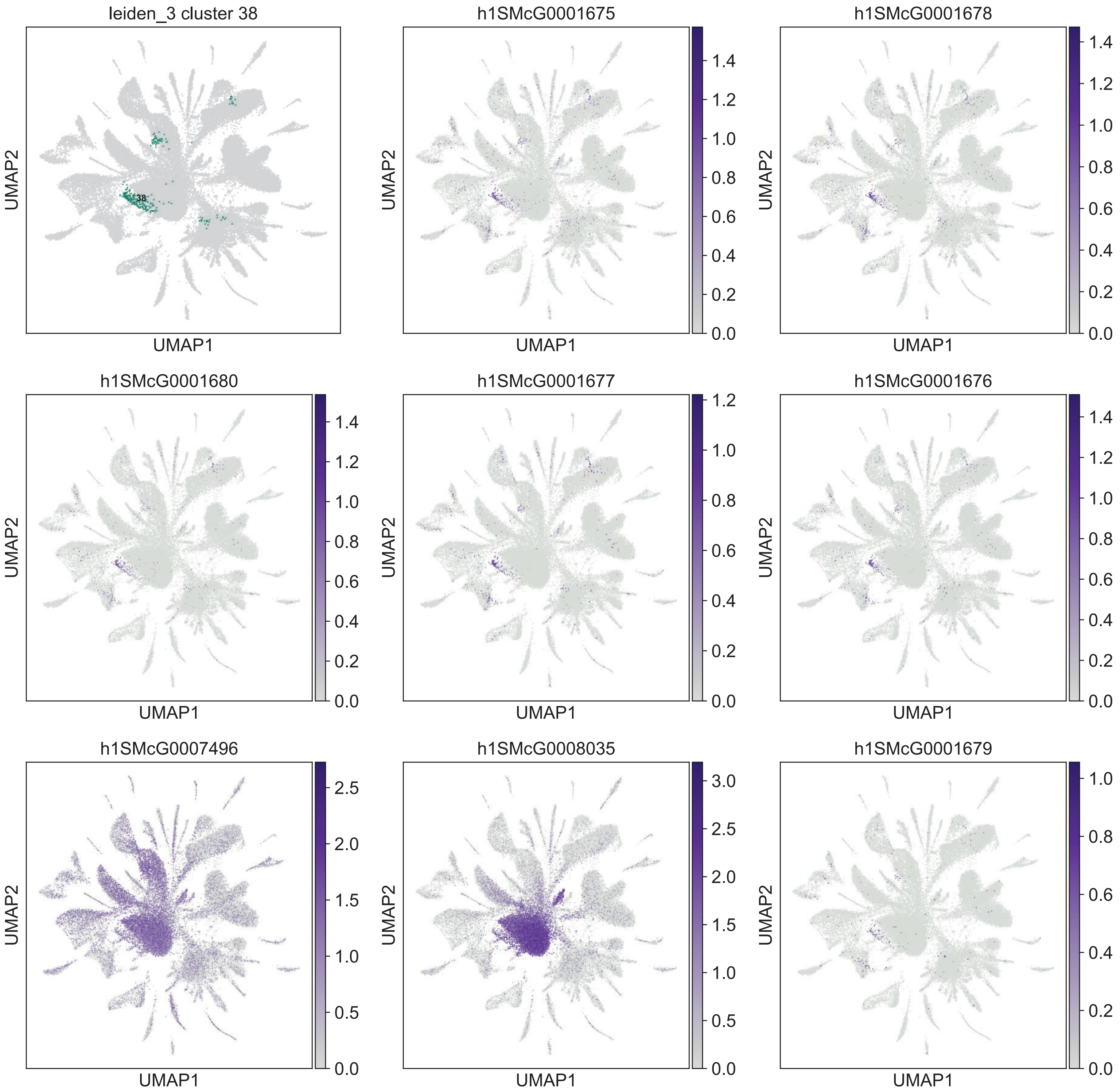
leiden_3 cluster 38
h1SMcG0001675
h1SMcG0001678
h1SMcG0001680
h1SMcG0001677
h1SMcG0001676
h1SMcG0007496
h1SMcG0008035
h1SMcG0001679
38
UMAP1
UMAP2
0.0
0.2
0.4
0.6
0.8
1.0
1.2
1.4
0.5
1.5
2.0
2.5
3.0

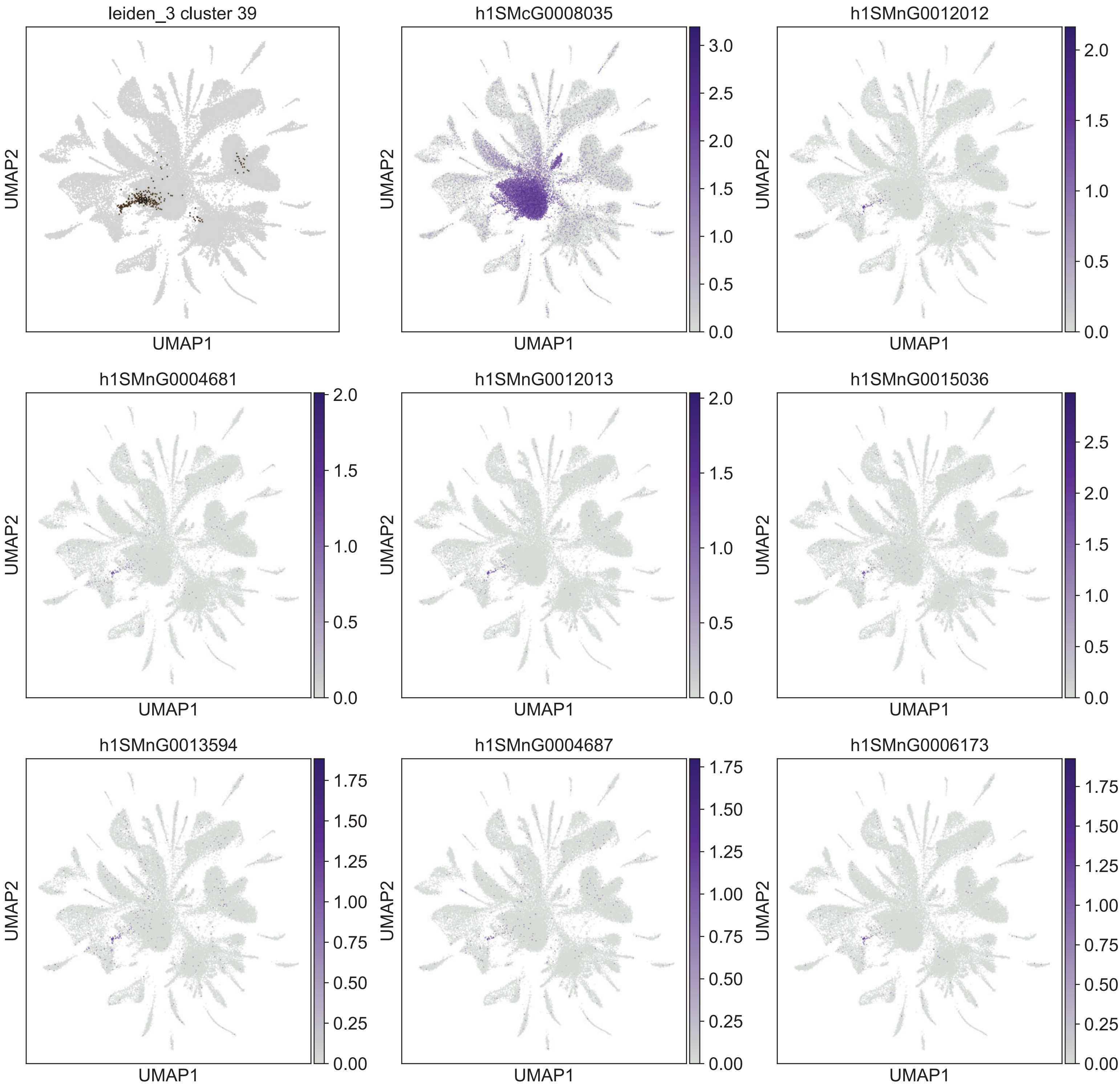

leiden_3 cluster 39
h1SMcG0008035
h1SMnG0012012
h1SMnG0004681
h1SMnG0012013
h1SMnG0015036
h1SMnG0013594
h1SMnG0004687
h1SMnG0006173
UMAP1
UMAP2

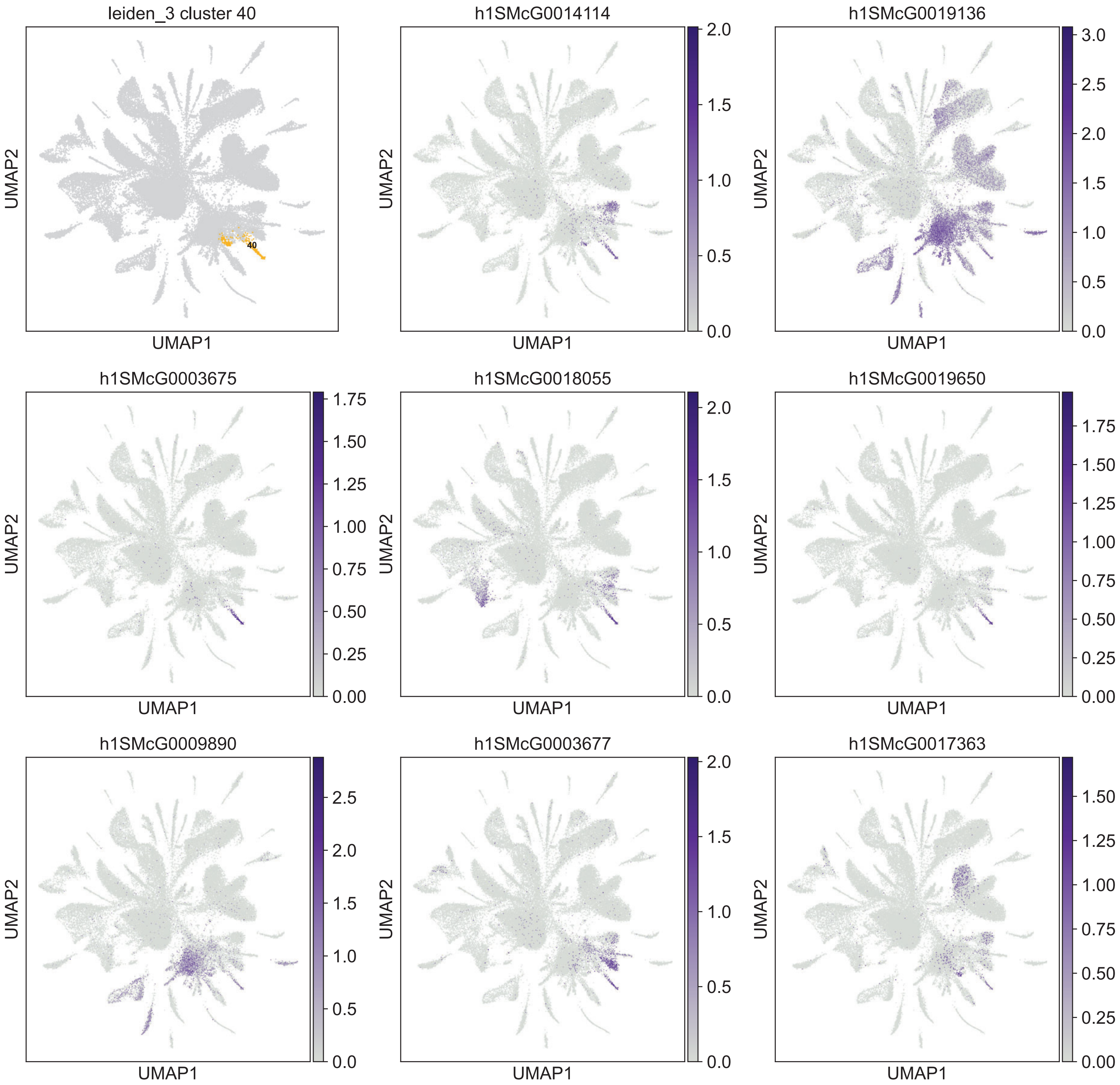

leiden_3 cluster 40
40
h1SMcG0014114
h1SMcG0019136
h1SMcG0003675
h1SMcG0018055
h1SMcG0019650
h1SMcG0009890
h1SMcG0003677
h1SMcG0017363
UMAP1
UMAP2

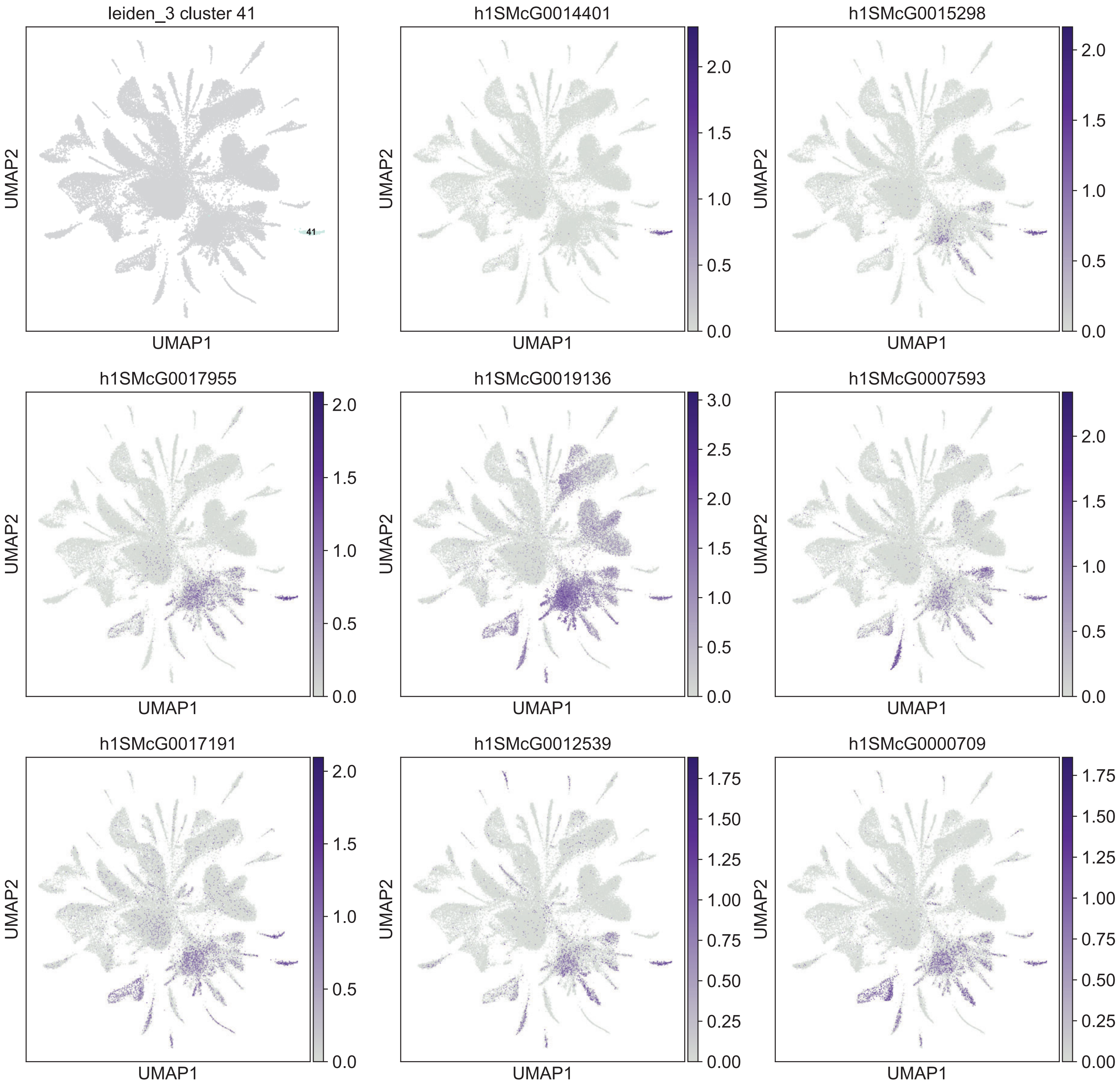
leiden_3 cluster 41
41
h1SMcG0014401
h1SMcG0015298
h1SMcG0017955
h1SMcG0019136
h1SMcG0007593
h1SMcG0017191
h1SMcG0012539
h1SMcG0000709
UMAP1
UMAP2

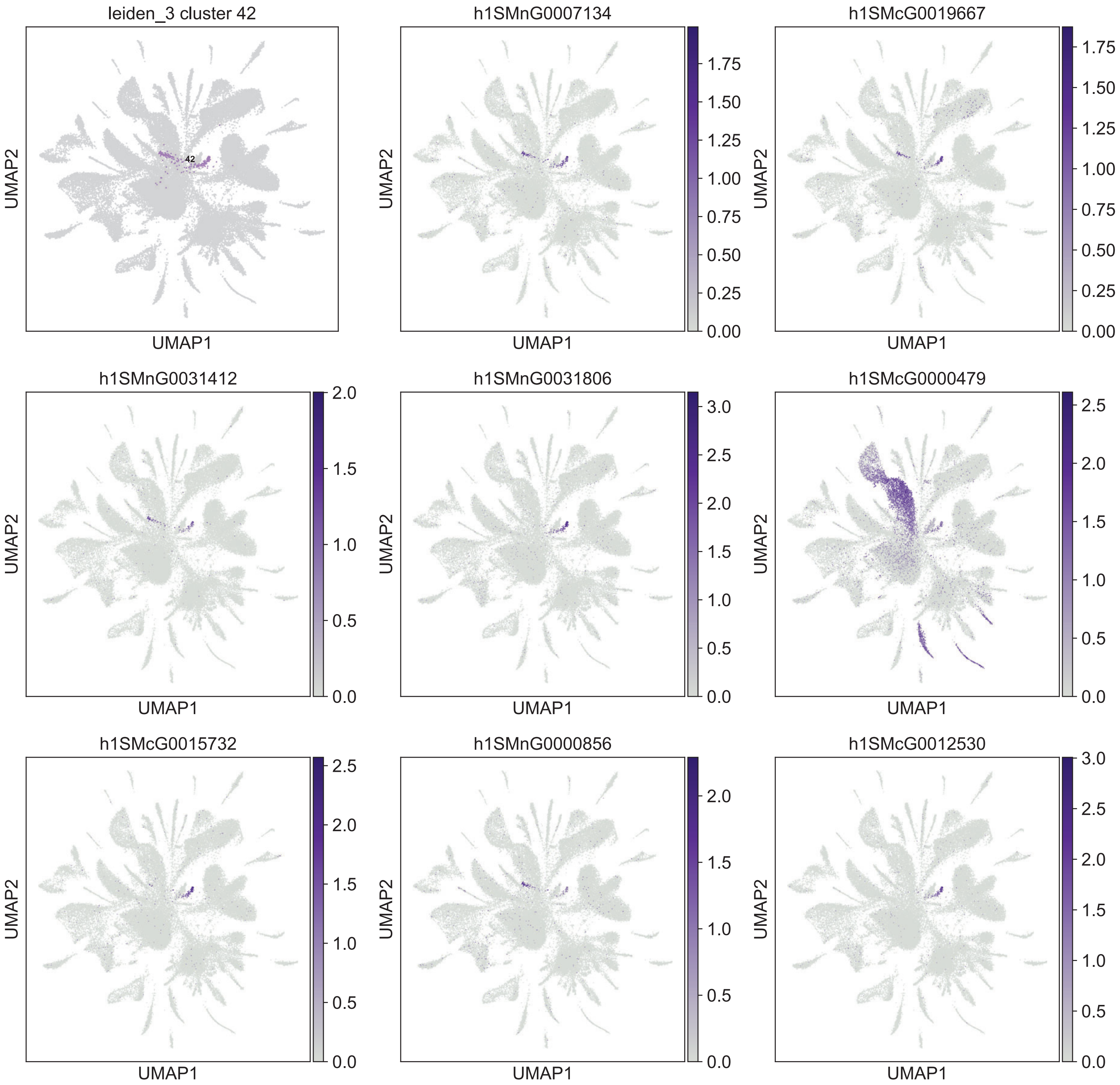

leiden_3 cluster 42
42
h1SMnG0007134
h1SMcG0019667
h1SMnG0031412
h1SMnG0031806
h1SMcG0000479
h1SMcG0015732
h1SMnG0000856
h1SMcG0012530
UMAP1
UMAP2
0.00
0.25
0.50
0.75
1.00
1.25
1.50
1.75
0.0
0.5
1.0
1.5
2.0
2.5
3.0

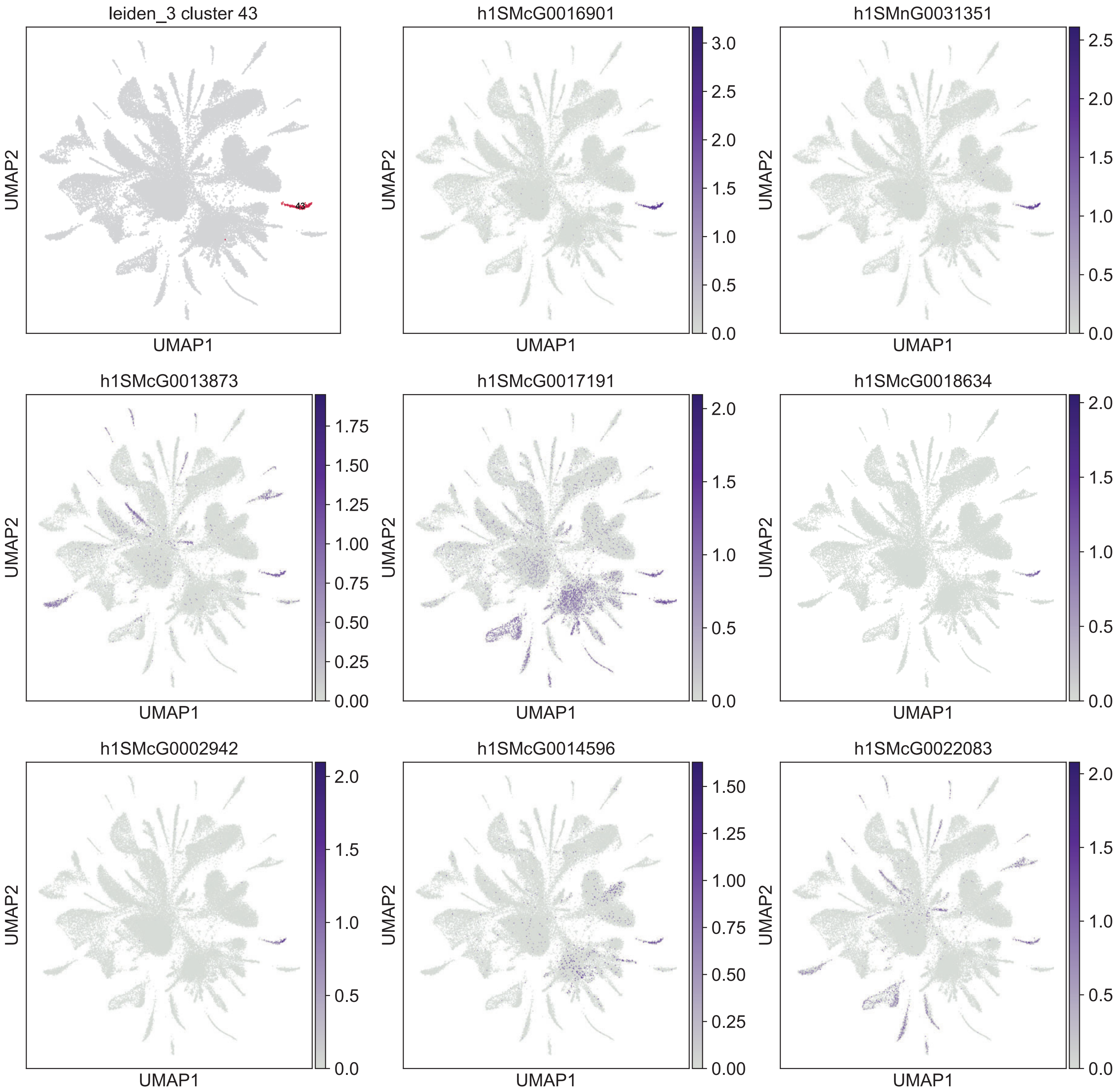

leiden_3 cluster 43
43
h1SMcG0016901
h1SMnG0031351
h1SMcG0013873
h1SMcG0017191
h1SMcG0018634
h1SMcG0002942
h1SMcG0014596
h1SMcG0022083
UMAP1
UMAP2

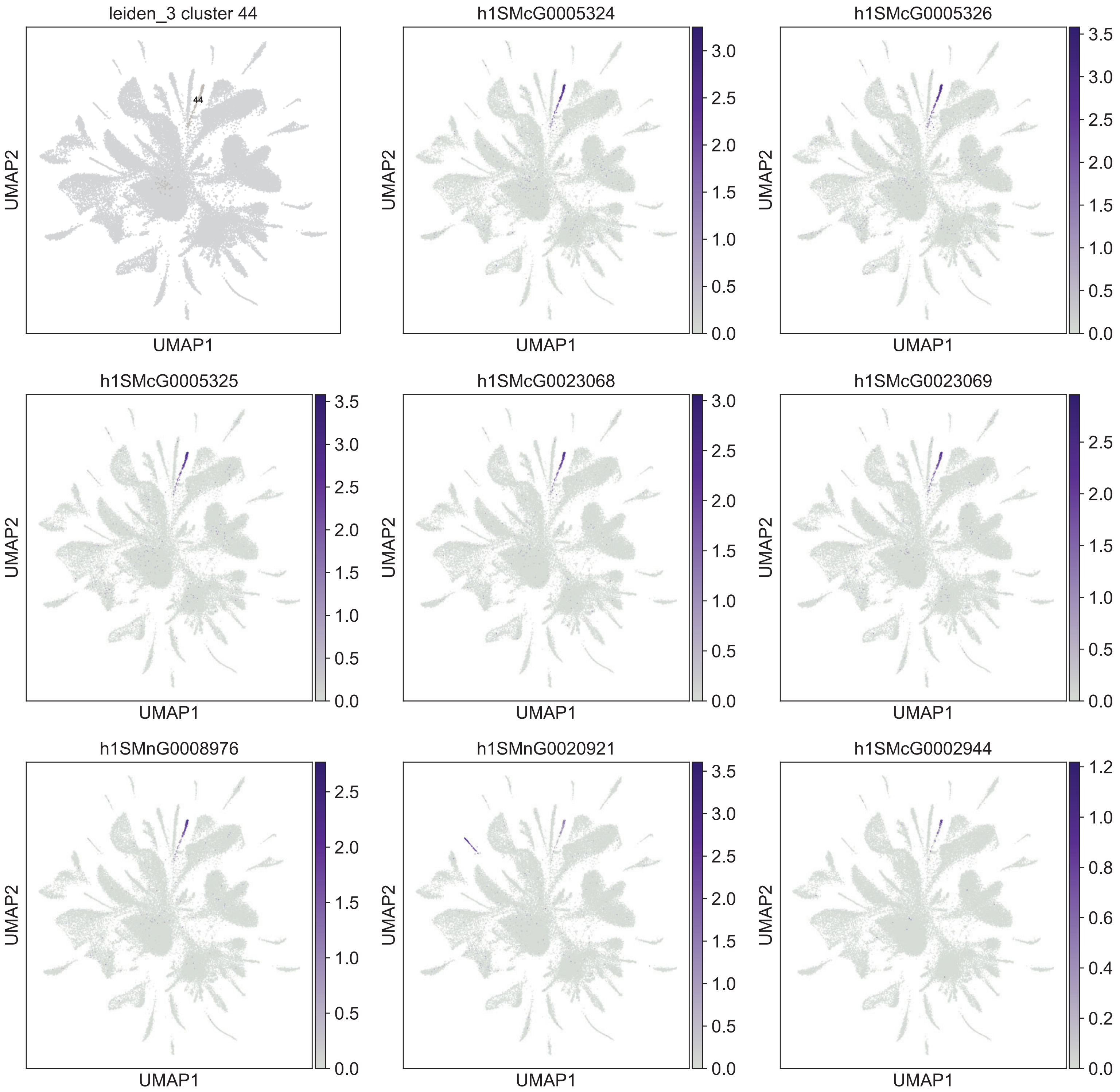
leiden_3 cluster 44
44
h1SMcG0005324
h1SMcG0005326
h1SMcG0005325
h1SMcG0023068
h1SMcG0023069
h1SMnG0008976
h1SMnG0020921
h1SMcG0002944
UMAP1
UMAP2

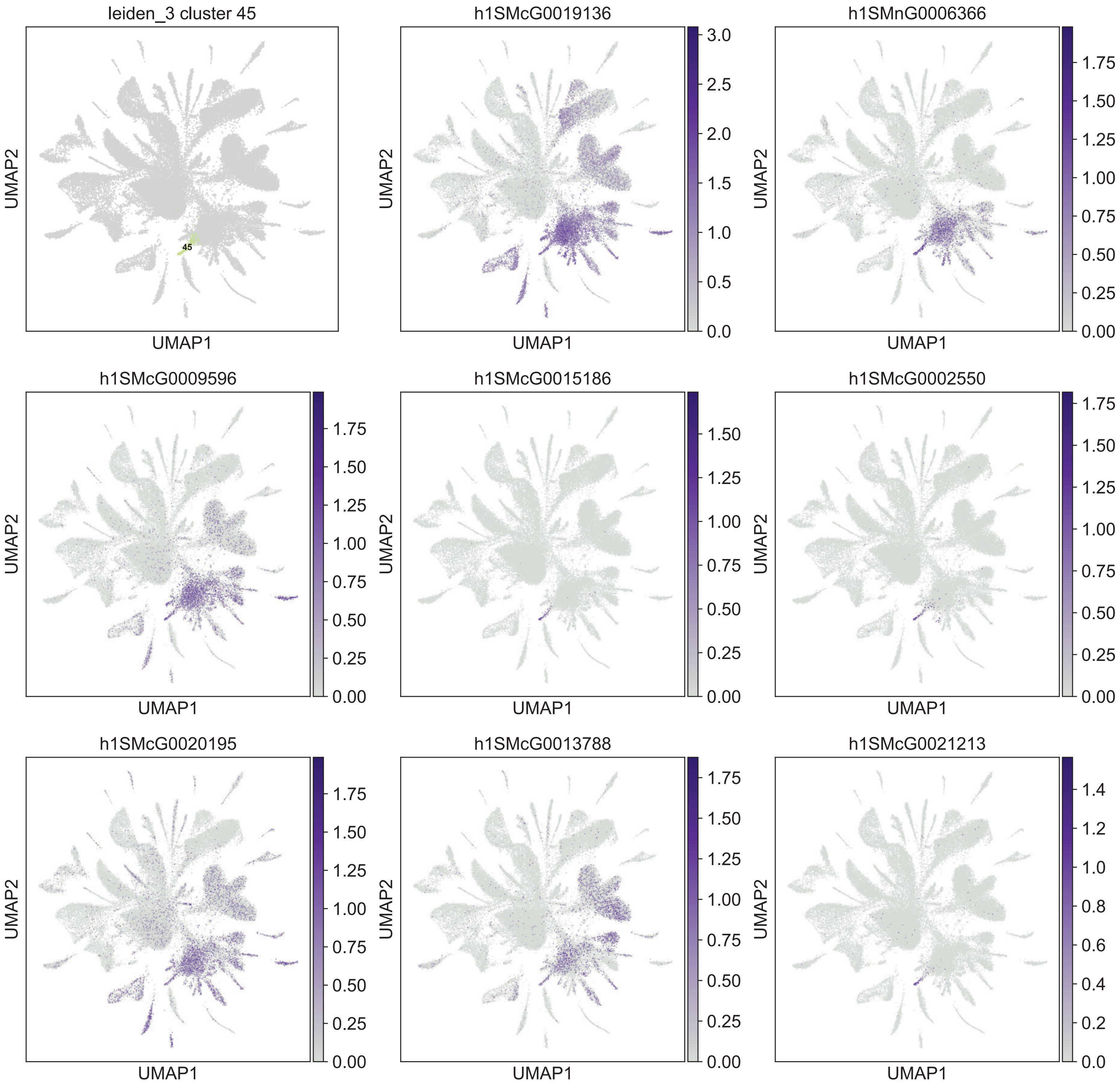

leiden_3 cluster 45
45
h1SMcG0019136
h1SMnG0006366
h1SMcG0009596
h1SMcG0015186
h1SMcG0002550
h1SMcG0020195
h1SMcG0013788
h1SMcG0021213
UMAP1
UMAP2

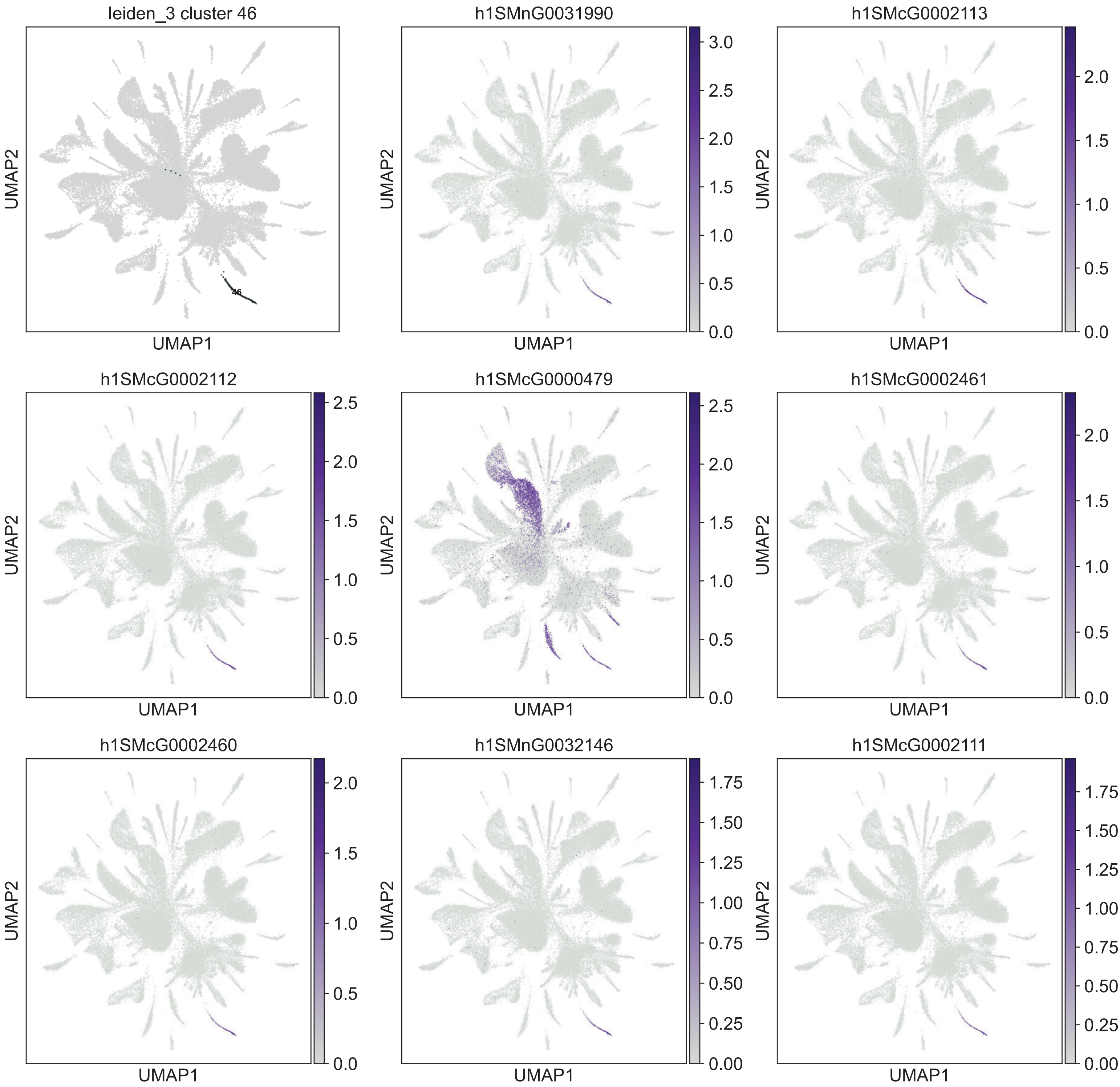
leiden_3 cluster 46
46
UMAP1
UMAP2
h1SMnG0031990
3.0
2.5
2.0
1.5
1.0
0.5
0.0
h1SMcG0002113
2.0
1.5
1.0
0.5
0.0
h1SMcG0002112
2.5
2.0
1.5
1.0
0.5
0.0
h1SMcG0000479
2.5
2.0
1.5
1.0
0.5
0.0
h1SMcG0002461
2.0
1.5
1.0
0.5
0.0
h1SMcG0002460
2.0
1.5
1.0
0.5
0.0
h1SMnG0032146
1.75
1.50
1.25
1.00
0.75
0.50
0.25
0.00
h1SMcG0002111
1.75
1.50
1.25
1.00
0.75
0.50
0.25
0.00

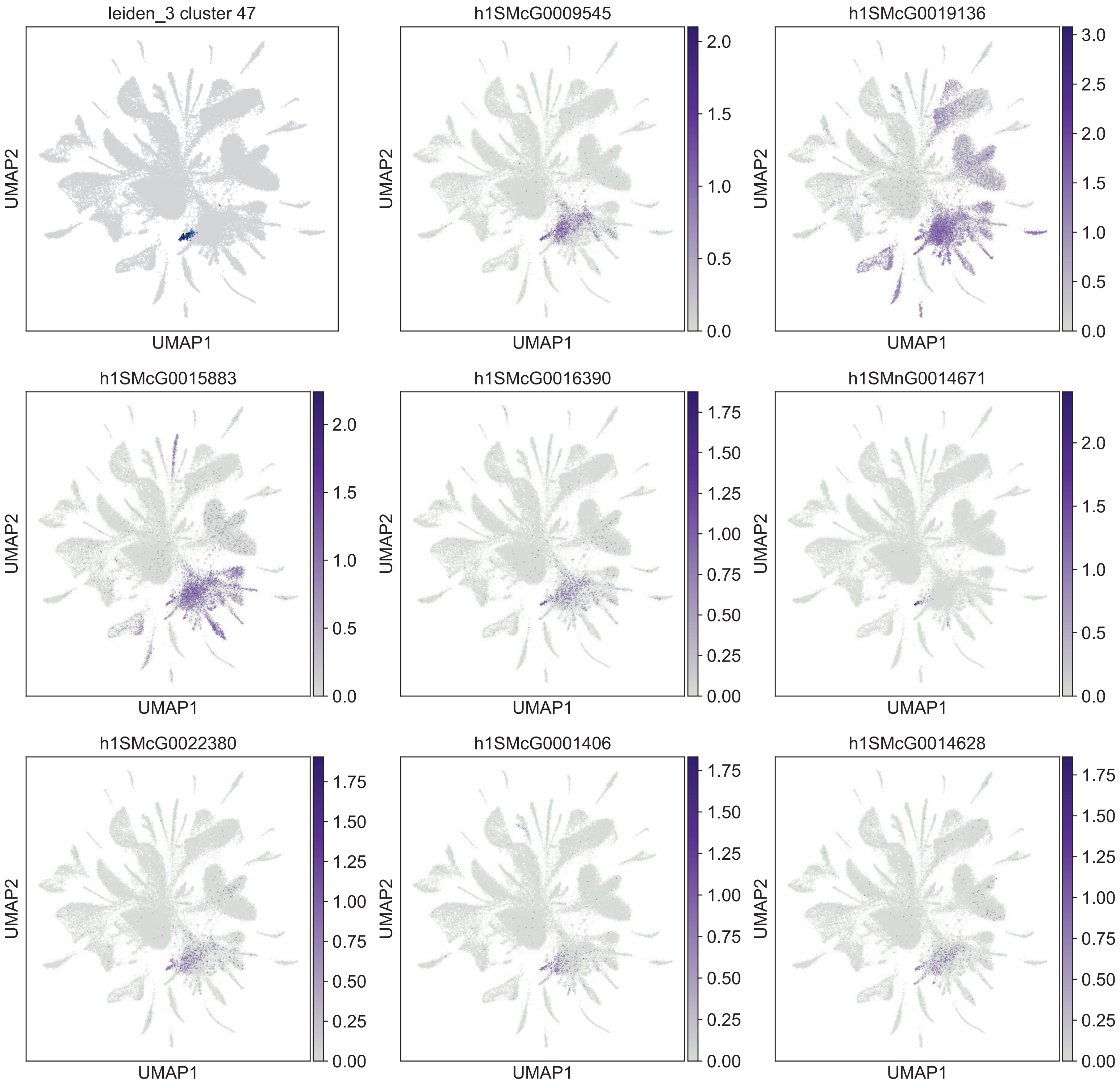

leiden_3 cluster 47
h1SMcG0009545
h1SMcG0019136
h1SMcG0015883
h1SMcG0016390
h1SMnG0014671
h1SMcG0022380
h1SMcG0001406
h1SMcG0014628
UMAP1
UMAP2

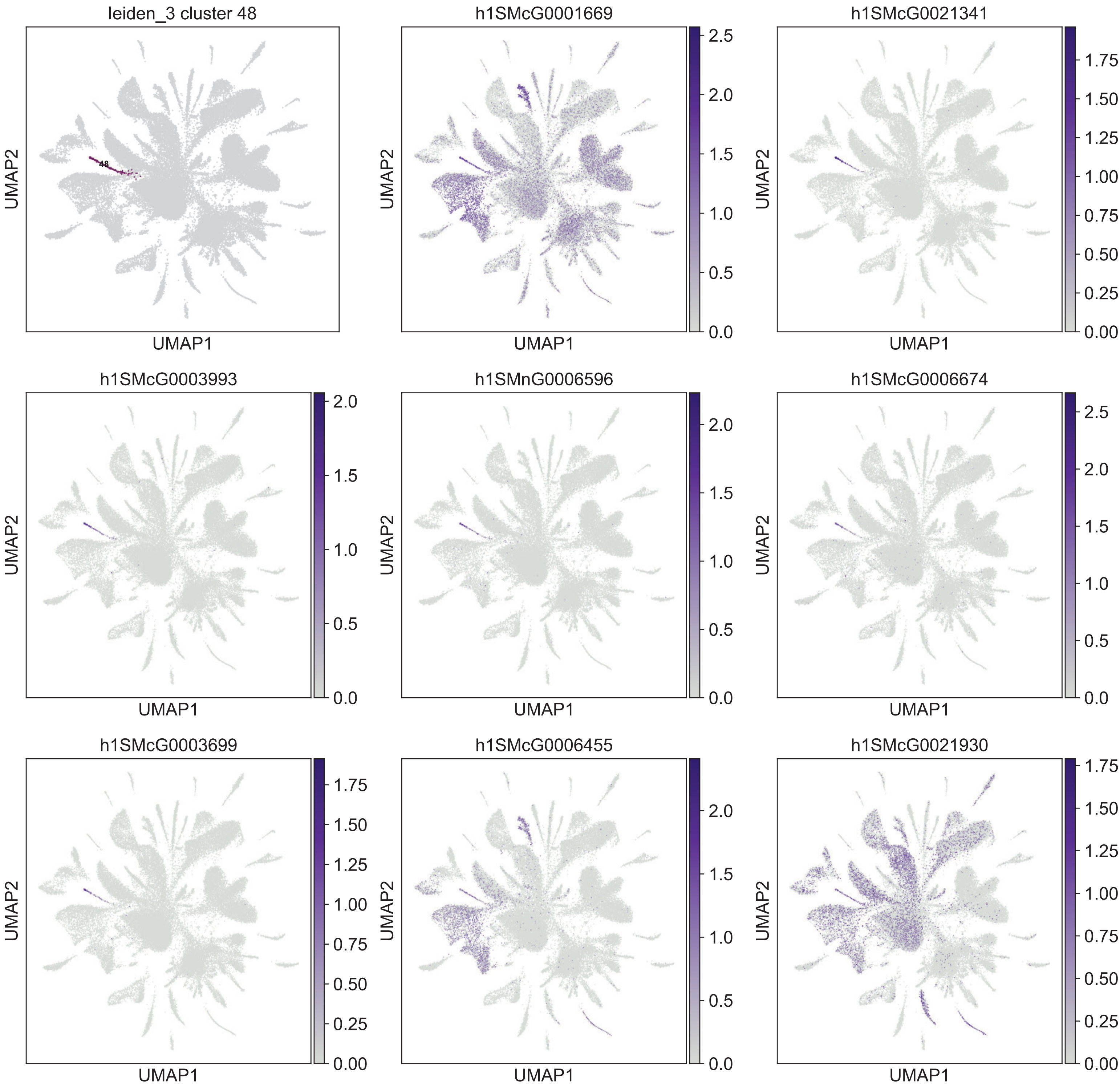
leiden_3 cluster 48
48
h1SMcG0001669
h1SMcG0021341
h1SMcG0003993
h1SMnG0006596
h1SMcG0006674
h1SMcG0003699
h1SMcG0006455
h1SMcG0021930
UMAP1
UMAP2

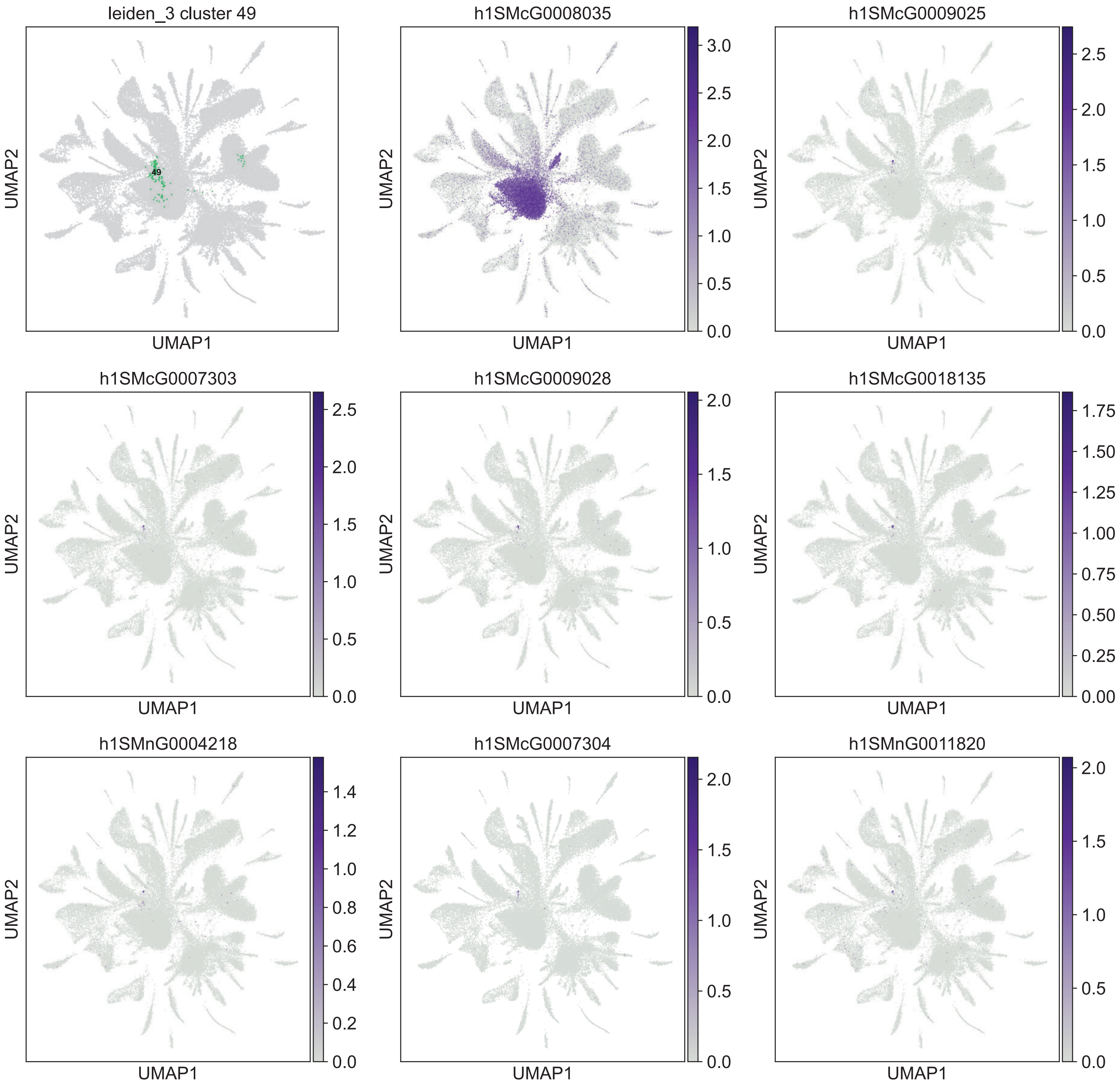
leiden_3 cluster 49
49
h1SMcG0008035
h1SMcG0009025
h1SMcG0007303
h1SMcG0009028
h1SMcG0018135
h1SMnG0004218
h1SMcG0007304
h1SMnG0011820
UMAP1
UMAP2

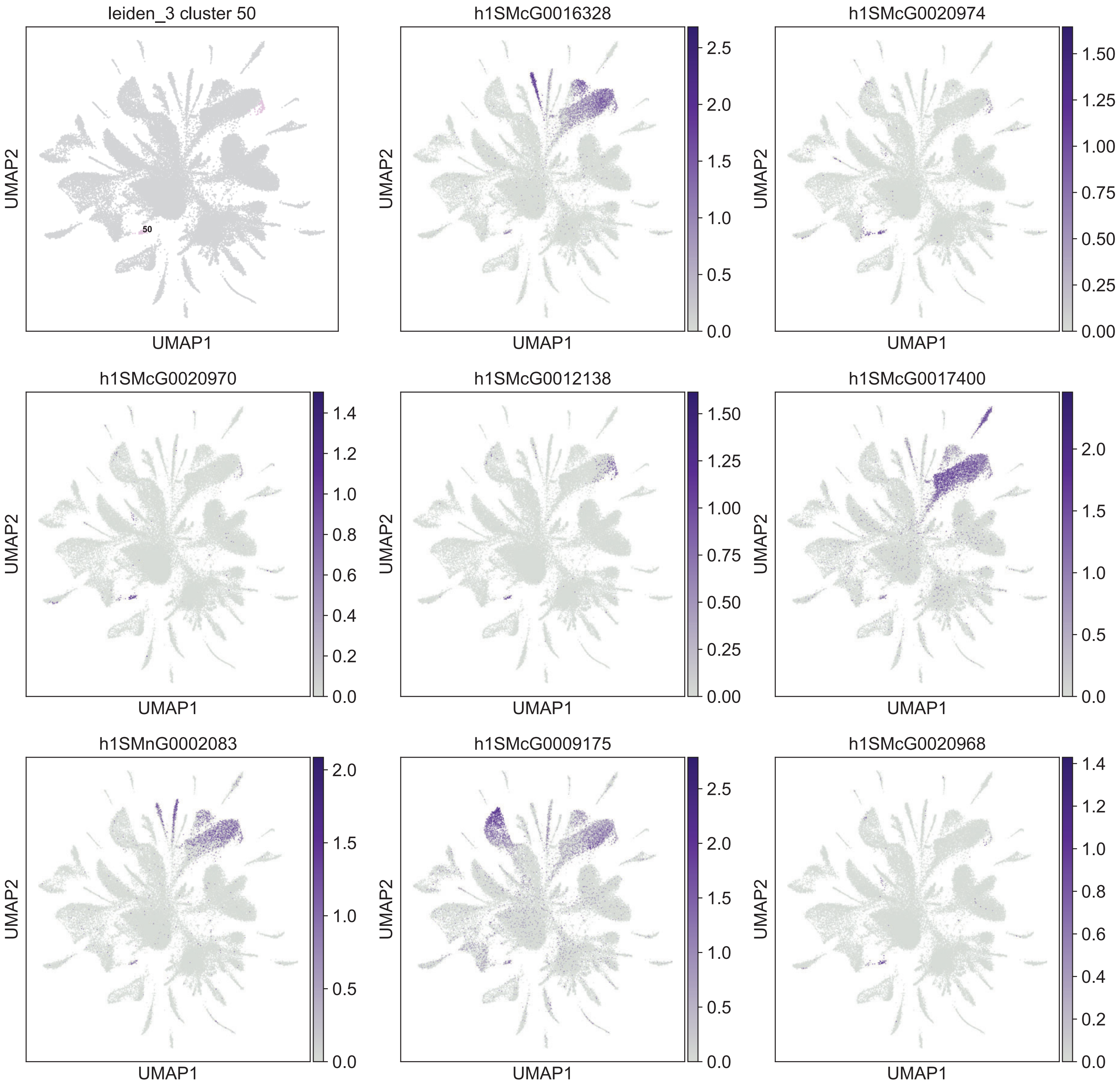

leiden_3 cluster 50
50
h1SMcG0016328
h1SMcG0020974
h1SMcG0020970
h1SMcG0012138
h1SMcG0017400
h1SMnG0002083
h1SMcG0009175
h1SMcG0020968
UMAP1
UMAP2

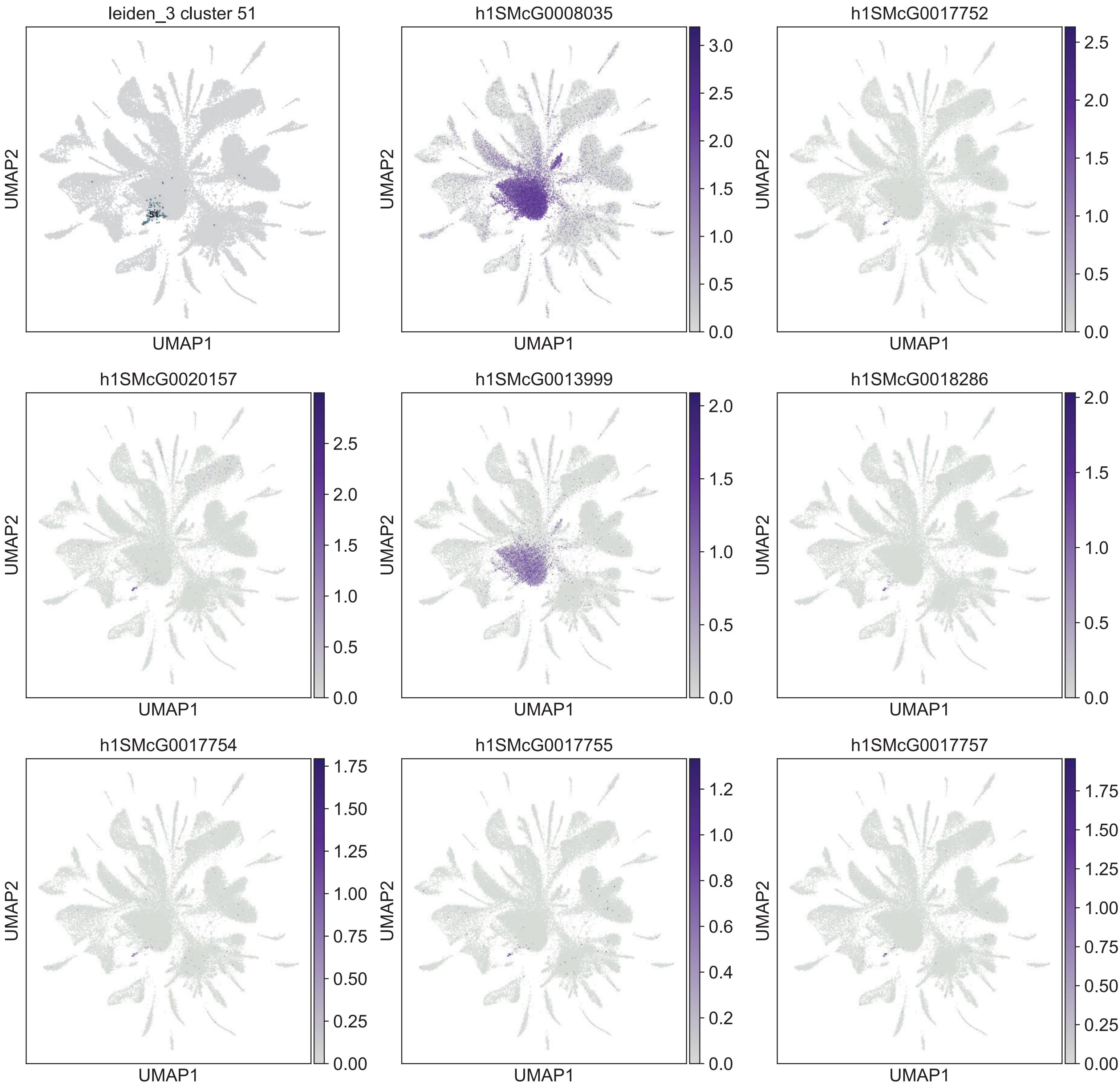
leiden_3 cluster 51
51
h1SMcG0008035
h1SMcG0017752
h1SMcG0020157
h1SMcG0013999
h1SMcG0018286
h1SMcG0017754
h1SMcG0017755
h1SMcG0017757
UMAP1
UMAP2
0.0
0.5
1.0
1.5
2.0
2.5
3.0
0.00
0.25
0.50
0.75
1.00
1.25
1.50
1.75
0.2
0.4
0.6
0.8
1.2

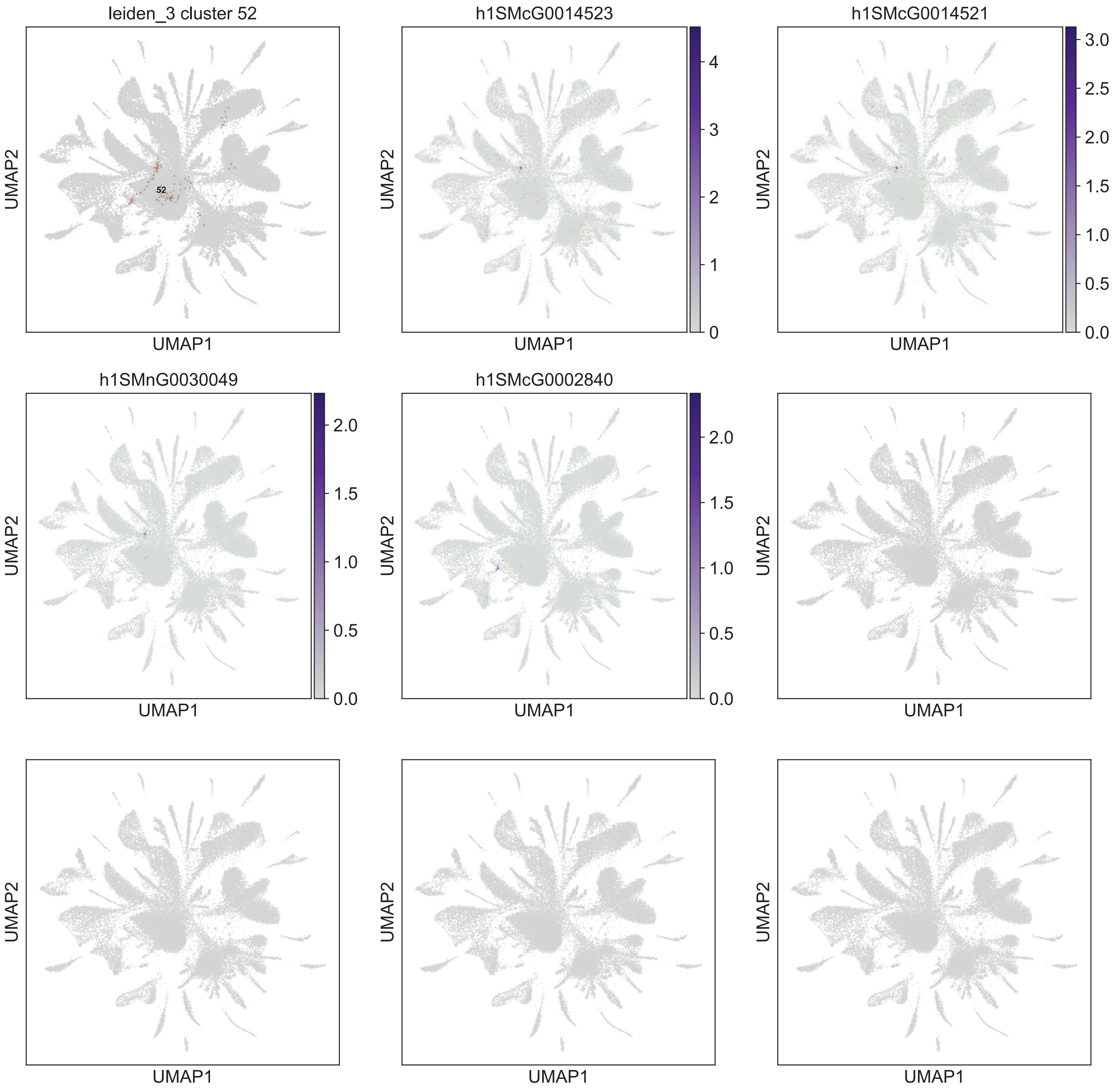
leiden_3 cluster 52
52
h1SMcG0014523
h1SMcG0014521
h1SMnG0030049
h1SMcG0002840
UMAP1
UMAP2

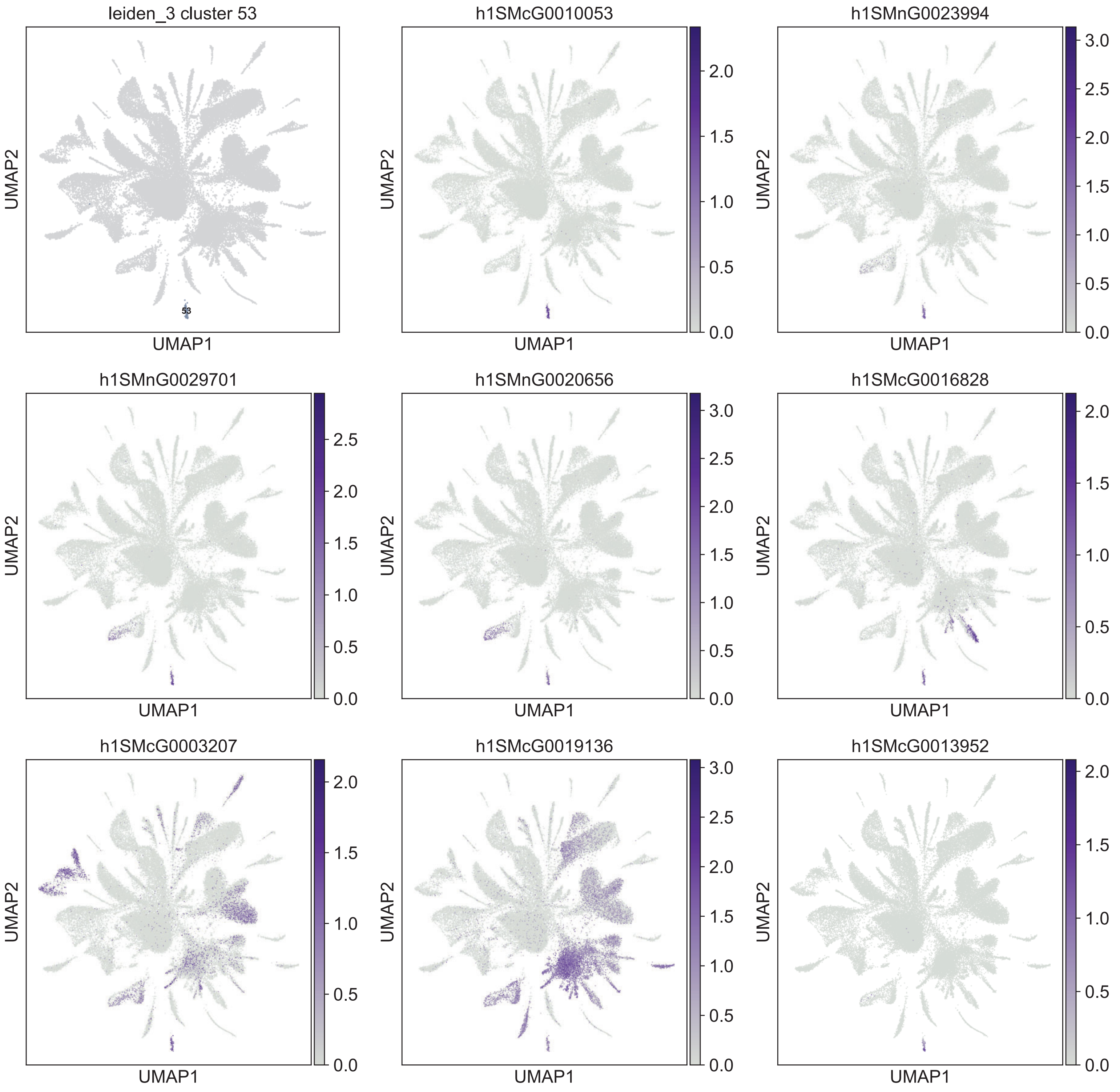

leiden_3 cluster 53
53
h1SMcG0010053
h1SMnG0023994
h1SMnG0029701
h1SMnG0020656
h1SMcG0016828
h1SMcG0003207
h1SMcG0019136
h1SMcG0013952
UMAP1
UMAP2

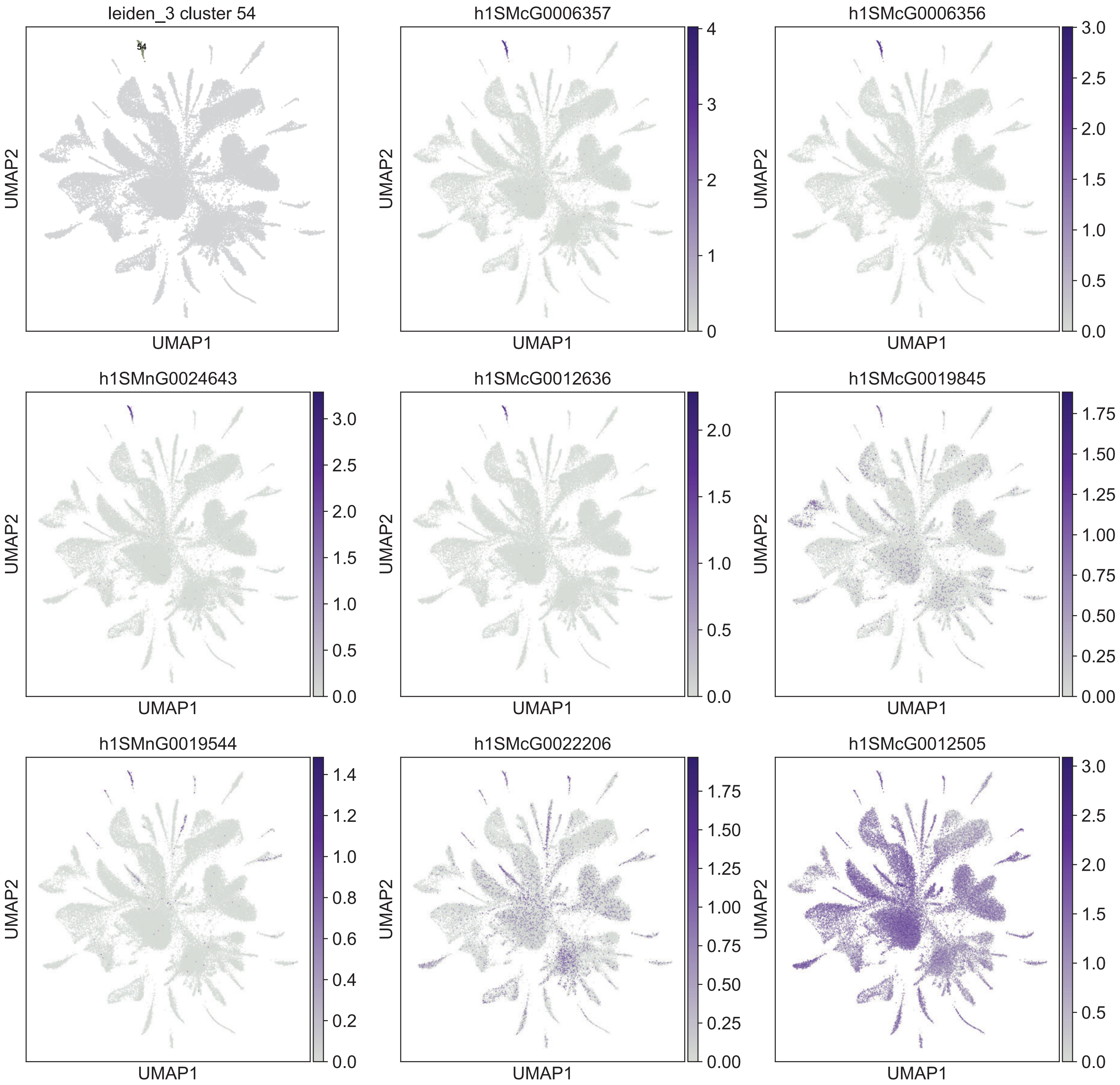
leiden_3 cluster 54
54
UMAP1
UMAP2
h1SMcG0006357
0
1
2
3
4
h1SMcG0006356
0.0
0.5
1.0
1.5
2.0
2.5
3.0
h1SMnG0024643
h1SMcG0012636
h1SMcG0019845
0.00
0.25
0.50
0.75
1.00
1.25
1.50
1.75
h1SMnG0019544
0.0
0.2
0.4
0.6
0.8
1.0
1.2
1.4
h1SMcG0022206
h1SMcG0012505

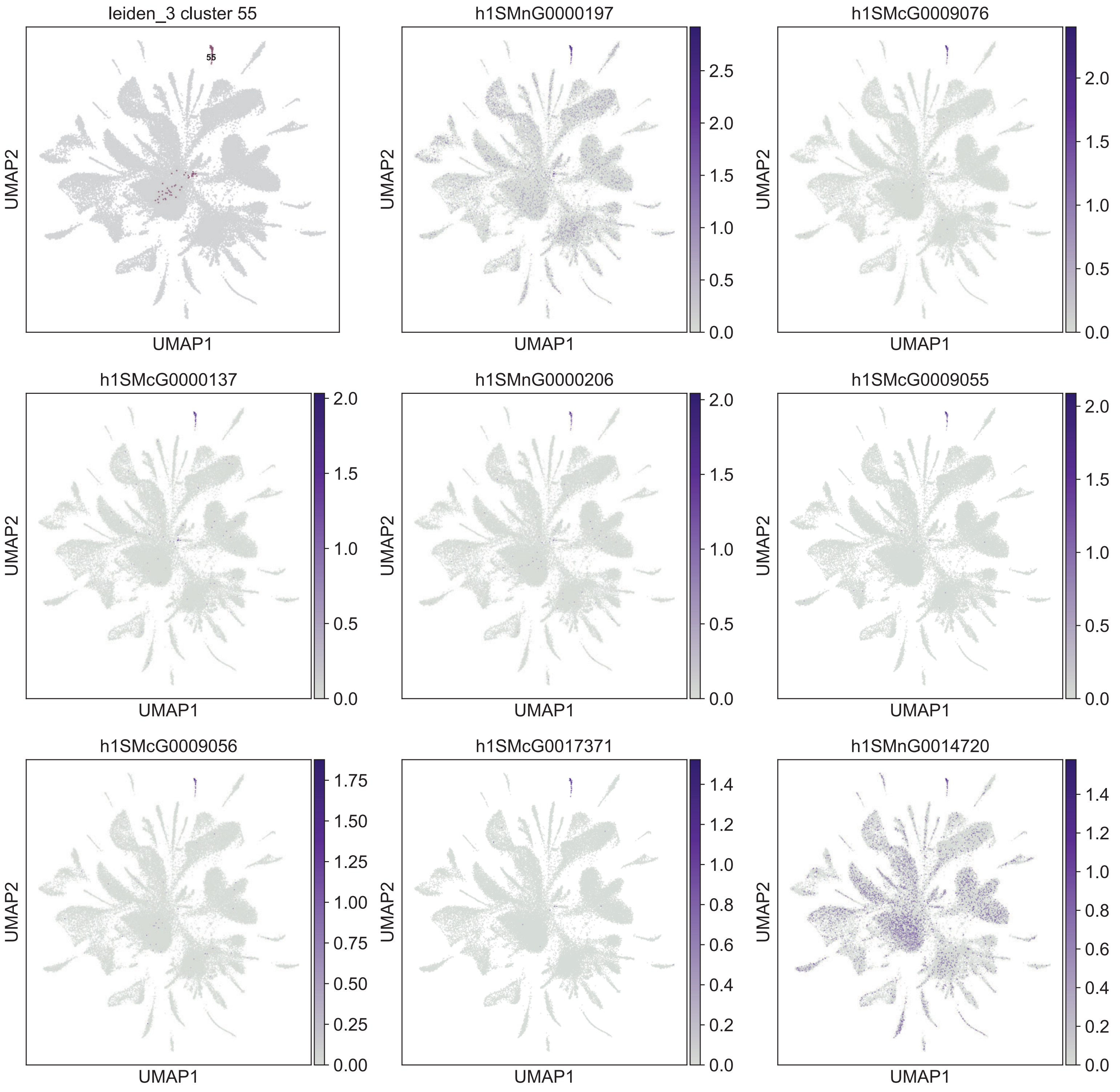
leiden_3 cluster 55
55
UMAP1
UMAP2
h1SMnG0000197
0.0 0.5 1.0 1.5 2.0 2.5
h1SMcG0009076
0.0 0.5 1.0 1.5 2.0
h1SMcG0000137
0.0 0.5 1.0 1.5 2.0
h1SMnG0000206
0.0 0.5 1.0 1.5 2.0
h1SMcG0009055
0.0 0.5 1.0 1.5 2.0
h1SMcG0009056
0.00 0.25 0.50 0.75 1.00 1.25 1.50 1.75
h1SMcG0017371
0.0 0.2 0.4 0.6 0.8 1.0 1.2 1.4
h1SMnG0014720
0.0 0.2 0.4 0.6 0.8 1.0 1.2 1.4

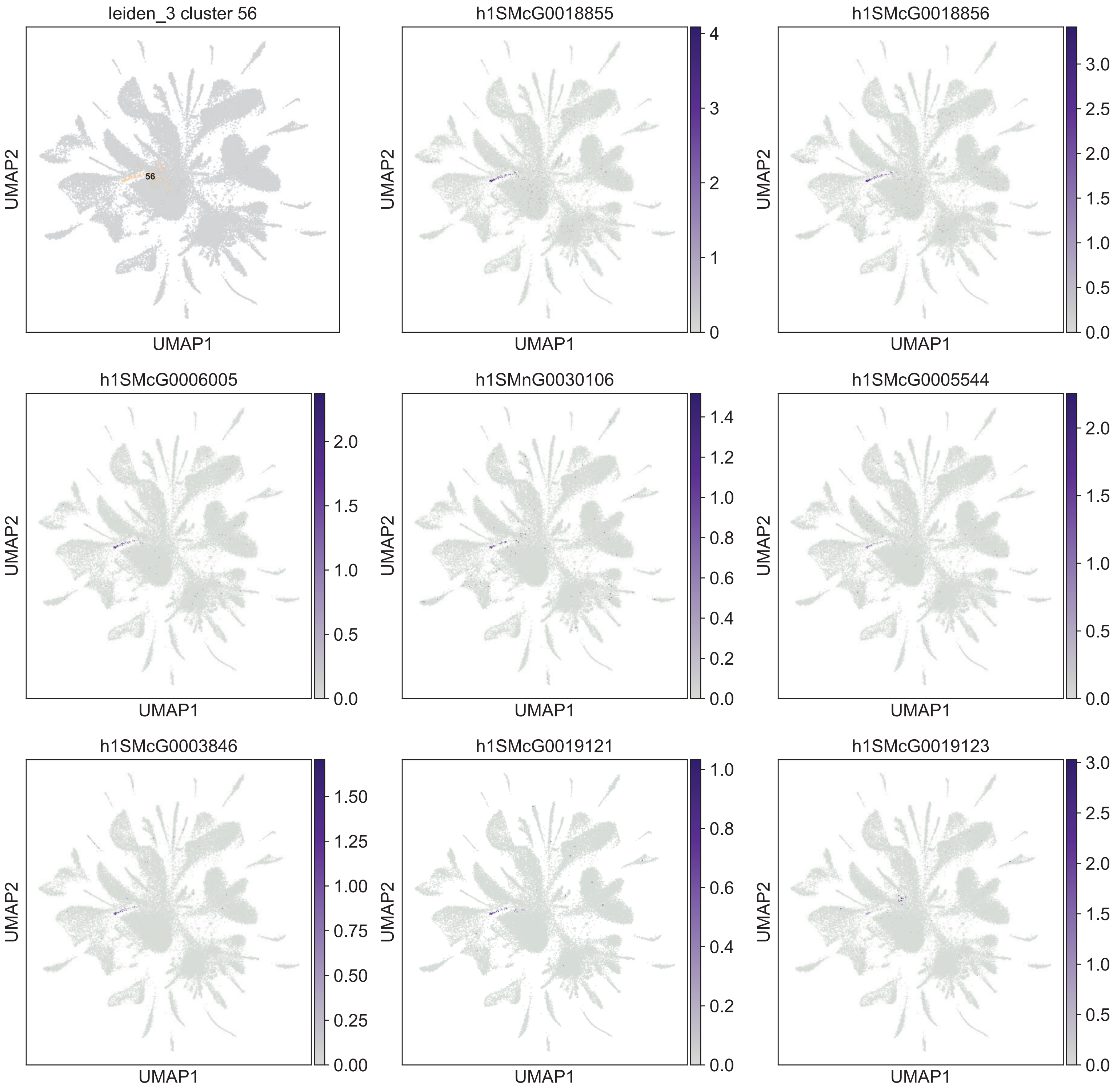
leiden_3 cluster 56
56
h1SMcG0018855
h1SMcG0018856
h1SMcG0006005
h1SMnG0030106
h1SMcG0005544
h1SMcG0003846
h1SMcG0019121
h1SMcG0019123
UMAP1
UMAP2
0
1
2
3
4
0.0
0.5
1.0
1.5
2.0
2.5
3.0
0.2
0.4
0.6
0.8
1.2
1.4
0.00
0.25
0.50
0.75
1.00
1.25
1.50

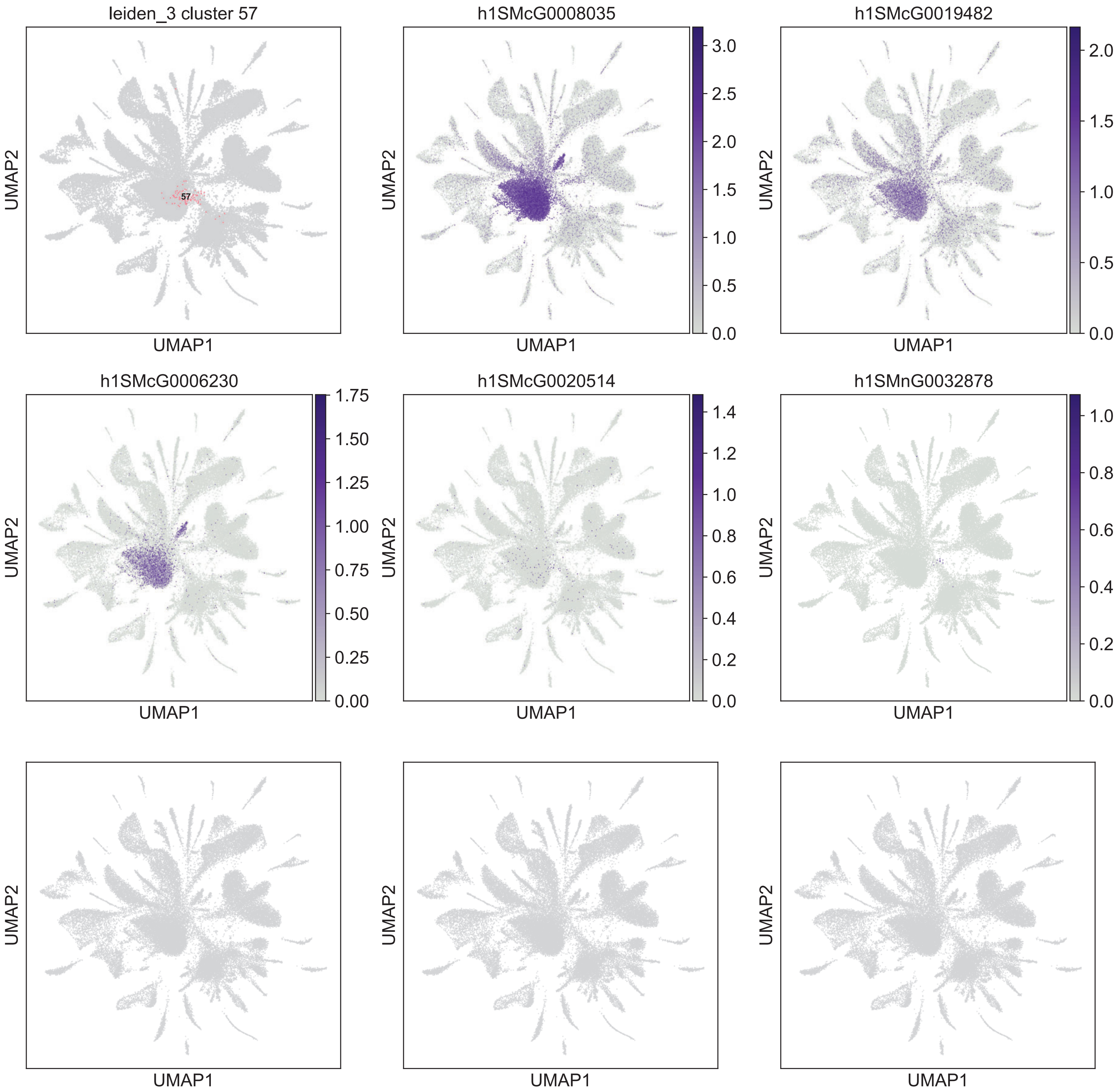
leiden_3 cluster 57
57
h1SMcG0008035
h1SMcG0019482
h1SMcG0006230
h1SMcG0020514
h1SMnG0032878
UMAP1
UMAP2
3.0
2.5
2.0
1.5
1.0
0.5
0.0
1.75
1.50
1.25
1.00
0.75
0.50
0.25
0.00
1.4
1.2
0.8
0.6
0.4
0.2

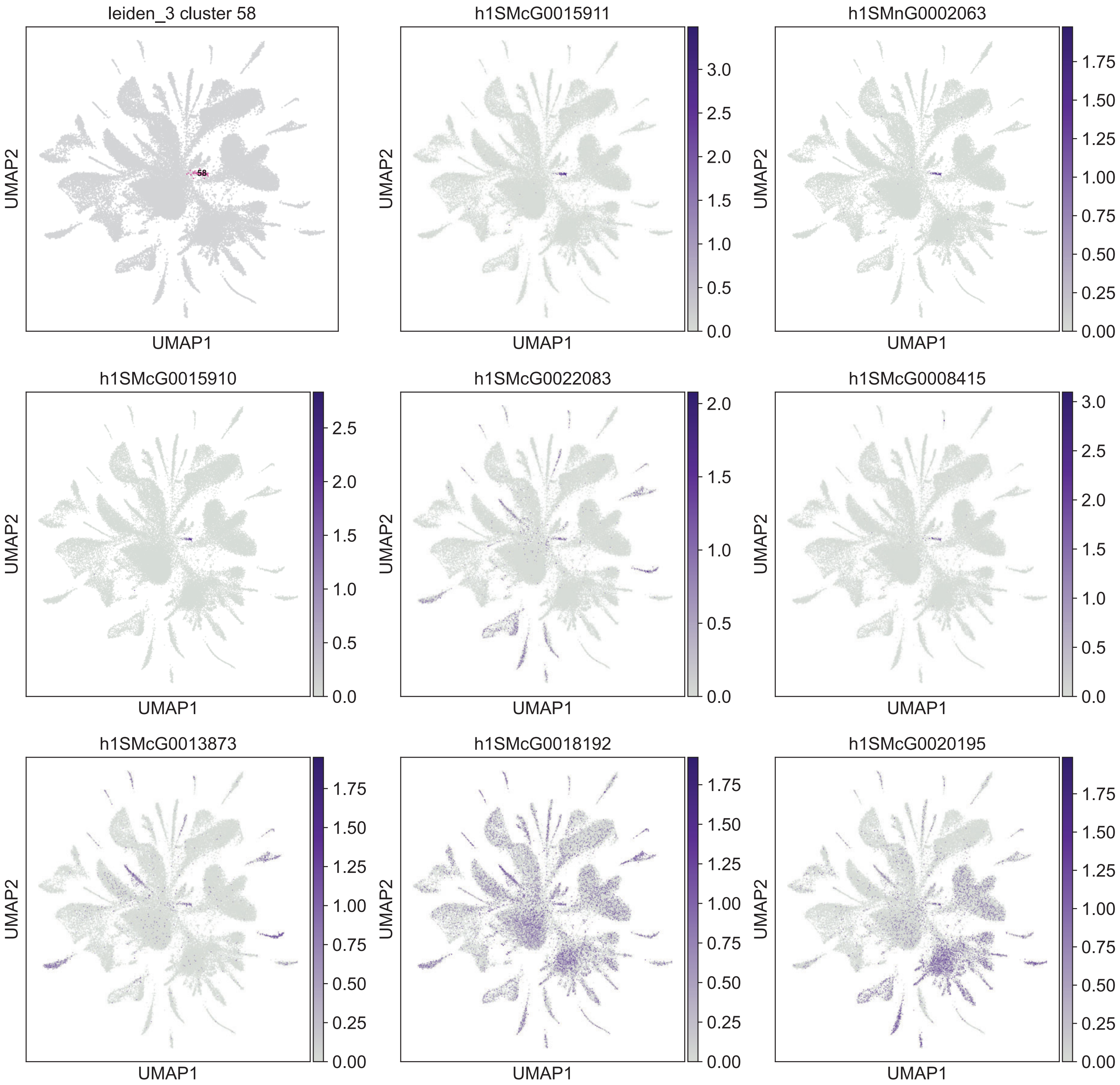

leiden_3 cluster 58
58
UMAP1
UMAP2
h1SMcG0015911
h1SMnG0002063
h1SMcG0015910
h1SMcG0022083
h1SMcG0008415
h1SMcG0013873
h1SMcG0018192
h1SMcG0020195

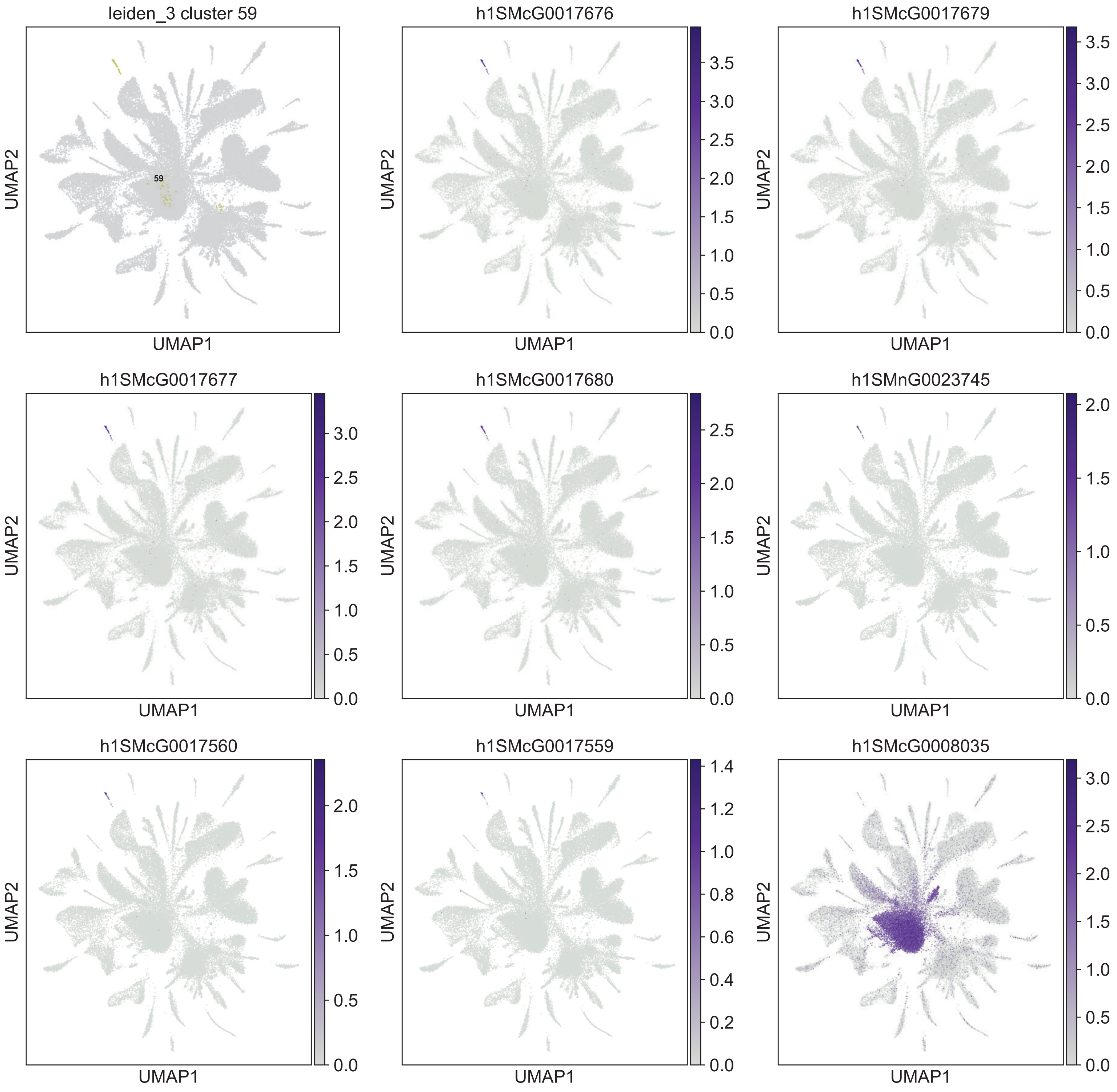
leiden_3 cluster 59
59
h1SMcG0017676
h1SMcG0017679
h1SMcG0017677
h1SMcG0017680
h1SMnG0023745
h1SMcG0017560
h1SMcG0017559
h1SMcG0008035
UMAP1
UMAP2

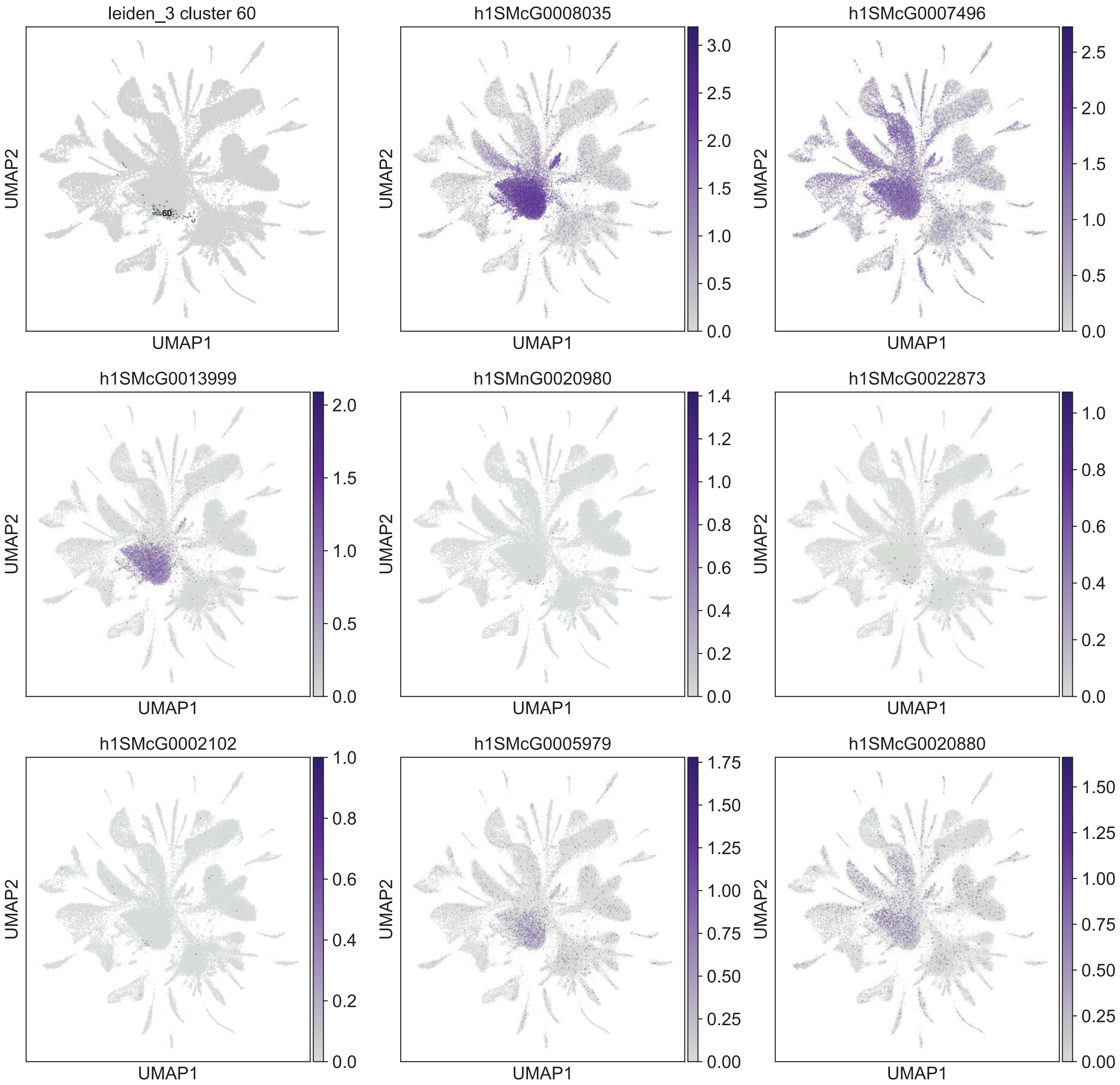

leiden_3 cluster 60
60
h1SMcG0008035
h1SMcG0007496
h1SMcG0013999
h1SMnG0020980
h1SMcG0022873
h1SMcG0002102
h1SMcG0005979
h1SMcG0020880
UMAP1
UMAP2

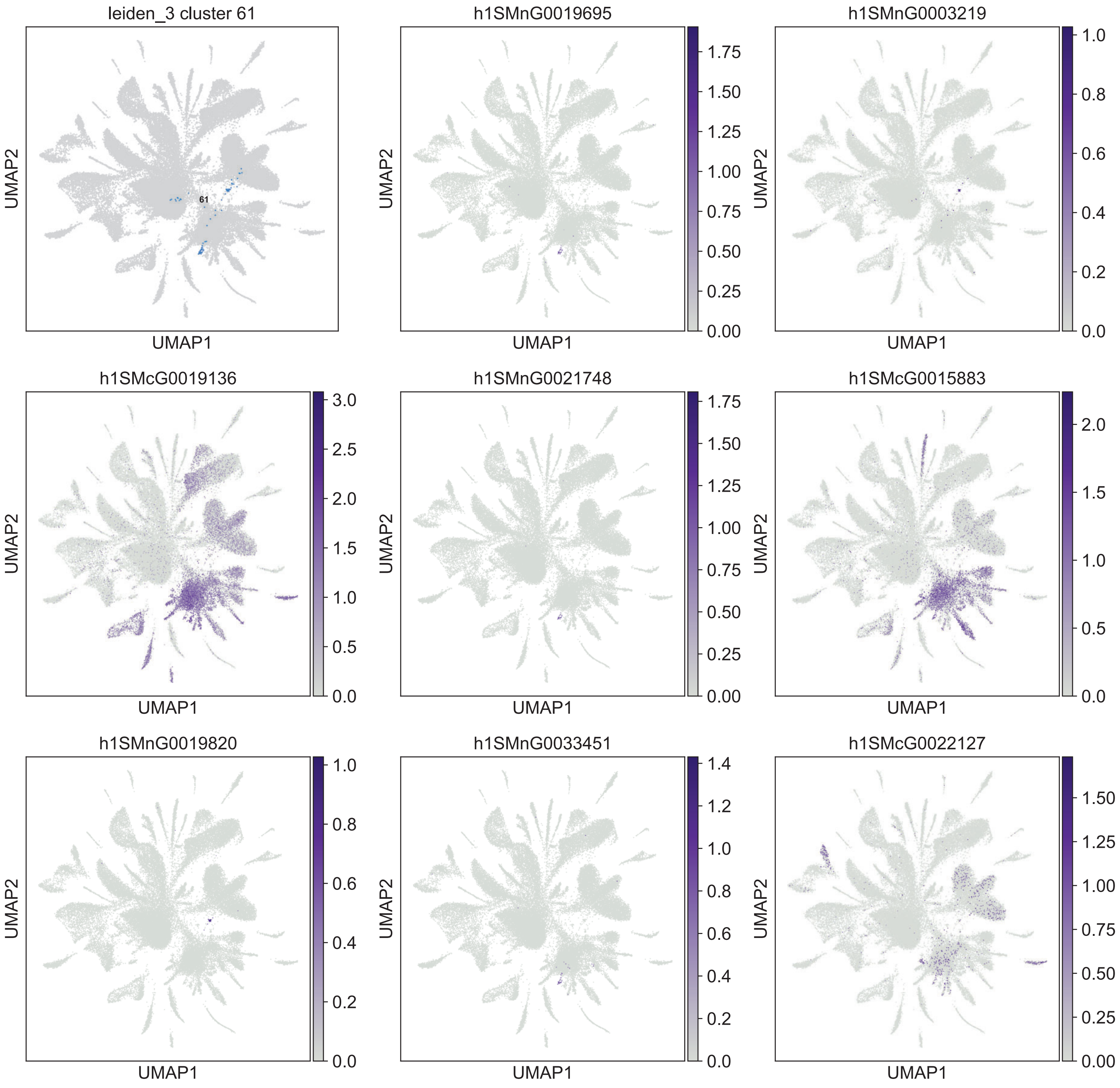

leiden_3 cluster 61
61
h1SMnG0019695
h1SMnG0003219
h1SMcG0019136
h1SMnG0021748
h1SMcG0015883
h1SMnG0019820
h1SMnG0033451
h1SMcG0022127
UMAP1
UMAP2

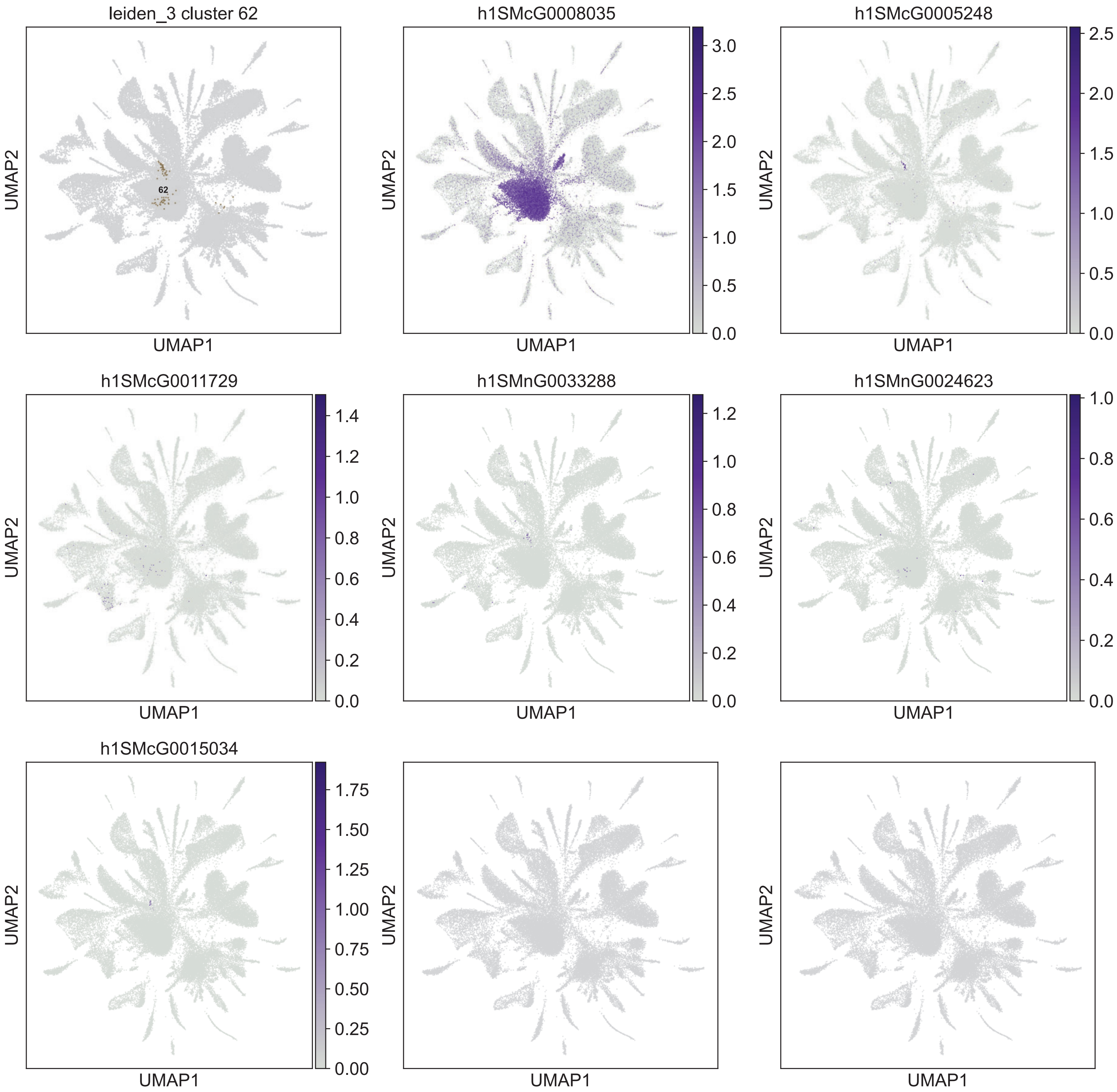

leiden_3 cluster 62
62
h1SMcG0008035
h1SMcG0005248
h1SMcG0011729
h1SMnG0033288
h1SMnG0024623
h1SMcG0015034
UMAP1
UMAP2
0.0
0.5
1.0
1.5
2.0
2.5
3.0
0.2
0.4
0.6
0.8
1.2
1.4
0.00
0.25
0.50
0.75
1.00
1.25
1.50
1.75

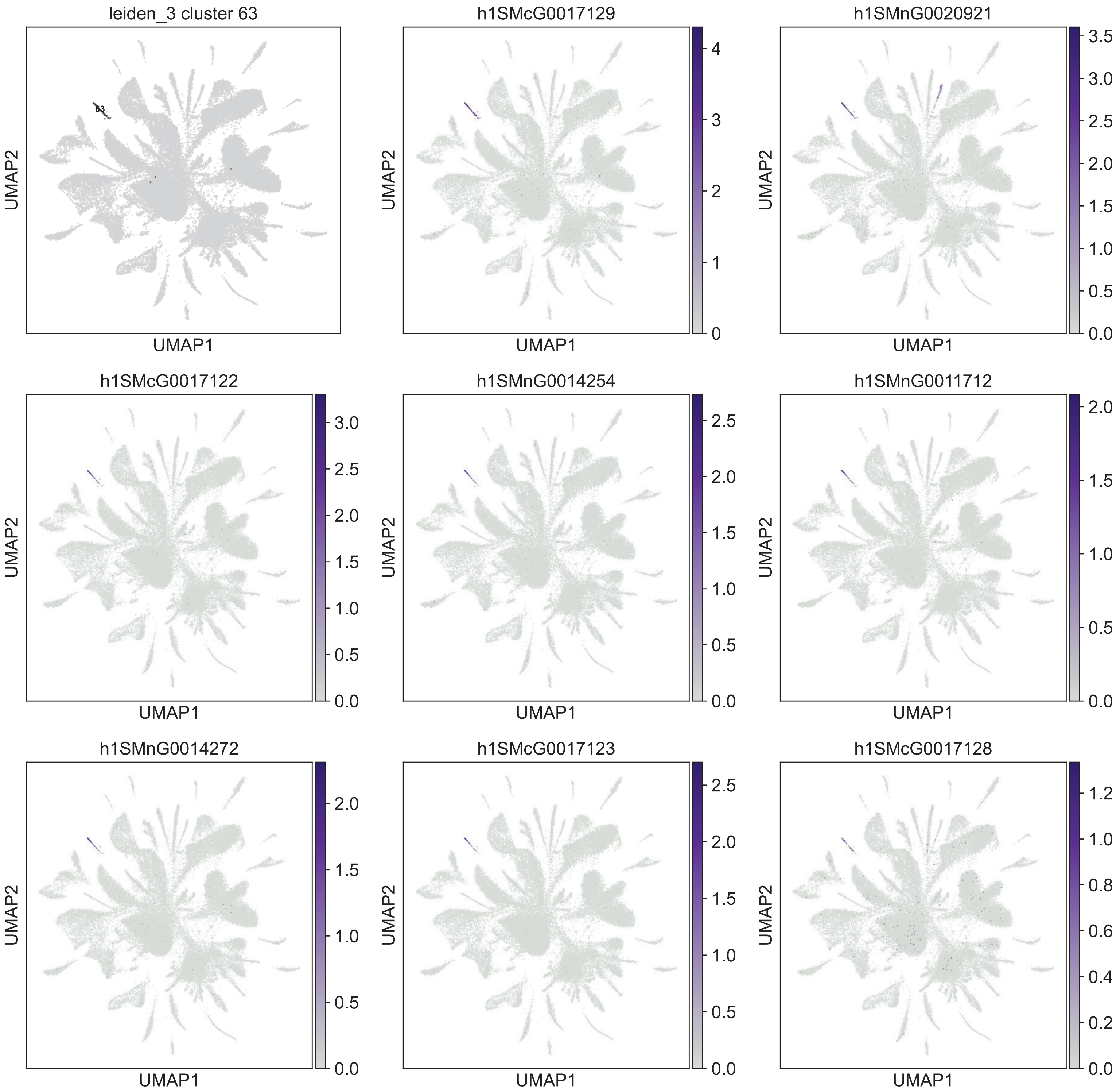

leiden_3 cluster 63
63
h1SMcG0017129
h1SMnG0020921
h1SMcG0017122
h1SMnG0014254
h1SMnG0011712
h1SMnG0014272
h1SMcG0017123
h1SMcG0017128
UMAP1
UMAP2

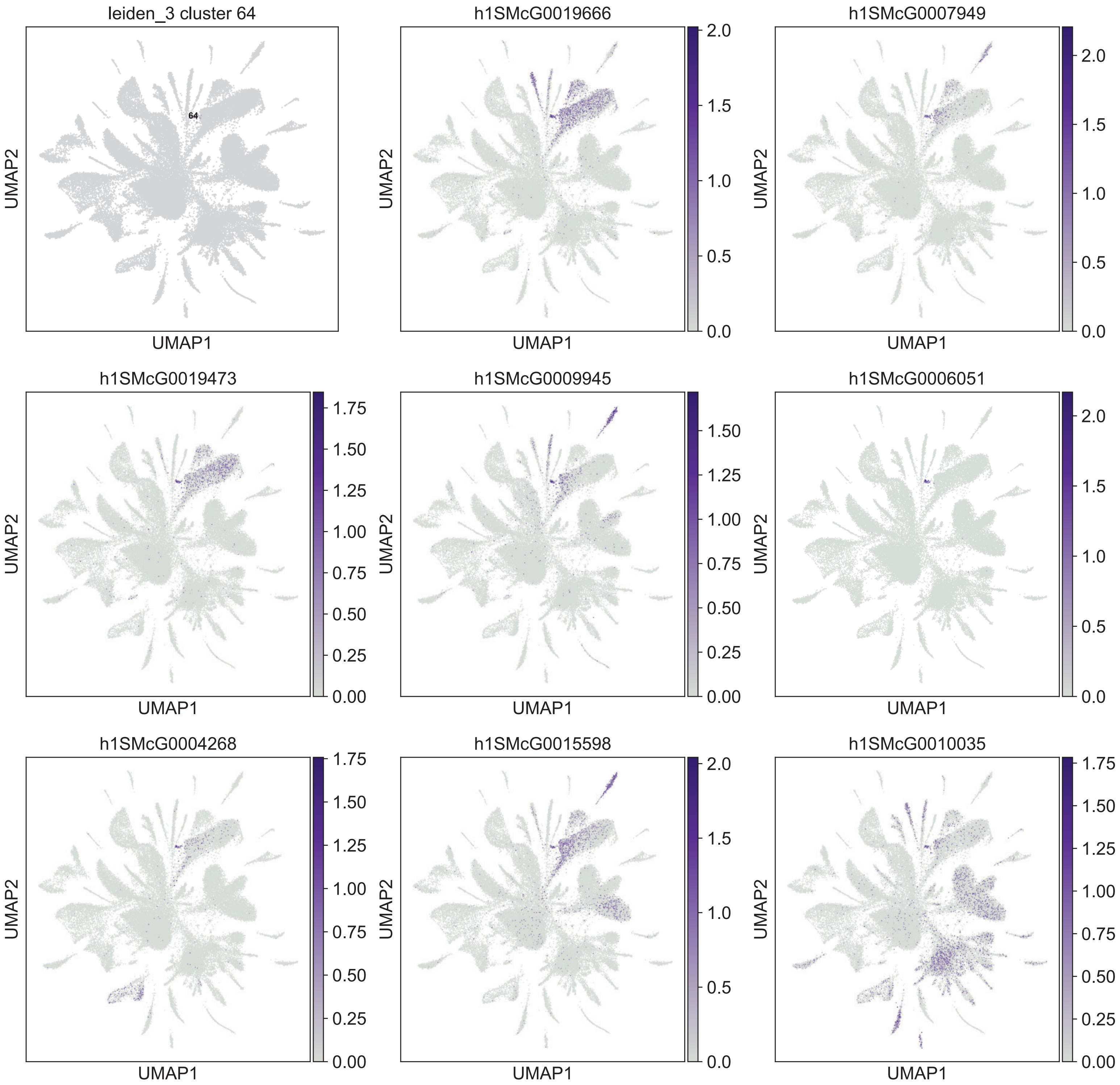

leiden_3 cluster 64
64
UMAP2
UMAP1
h1SMcG0019666
h1SMcG0007949
h1SMcG0019473
h1SMcG0009945
h1SMcG0006051
h1SMcG0004268
h1SMcG0015598
h1SMcG0010035

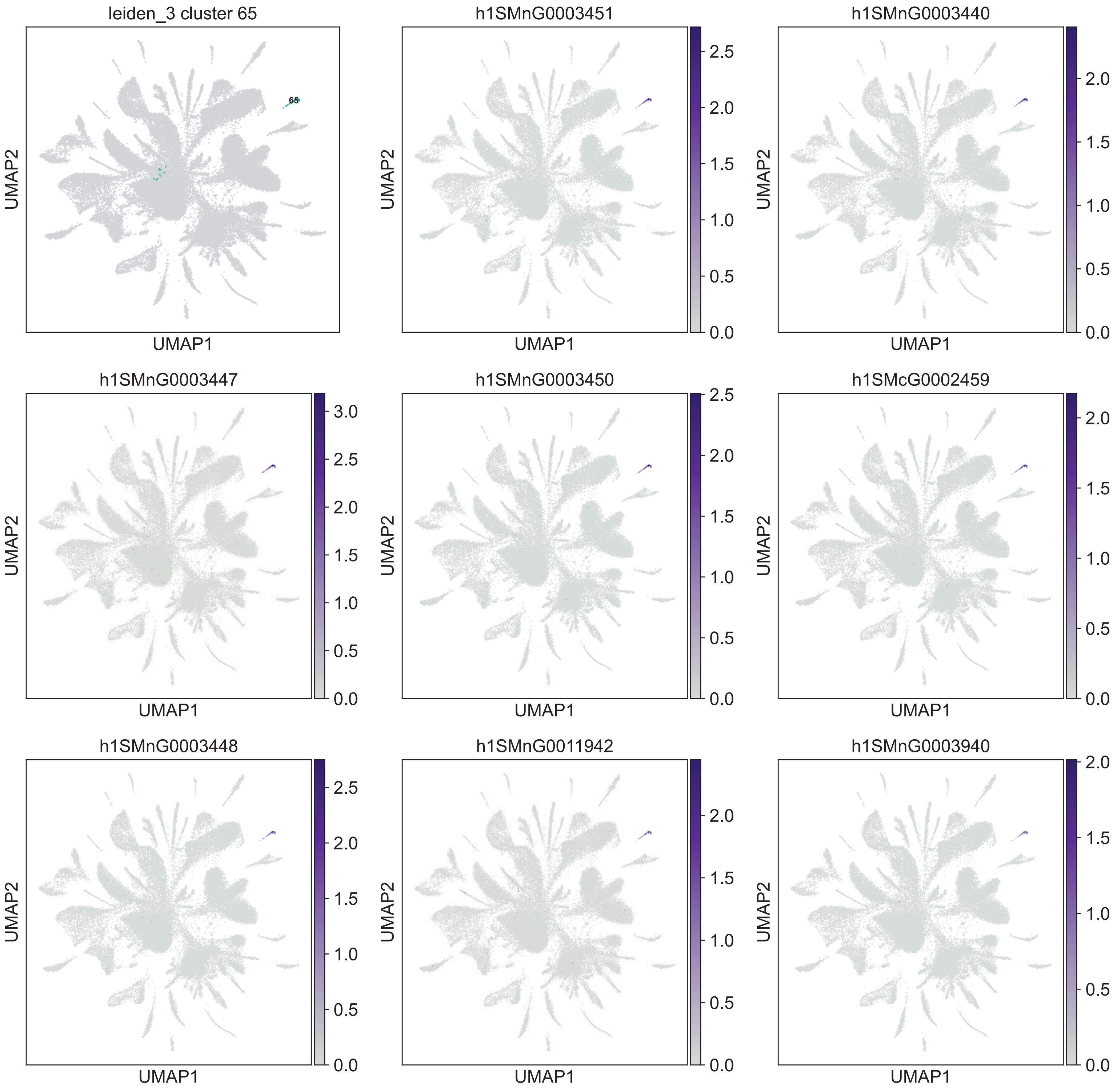
leiden_3 cluster 65
65
h1SMnG0003451
h1SMnG0003440
h1SMnG0003447
h1SMnG0003450
h1SMcG0002459
h1SMnG0003448
h1SMnG0011942
h1SMnG0003940
UMAP1
UMAP2

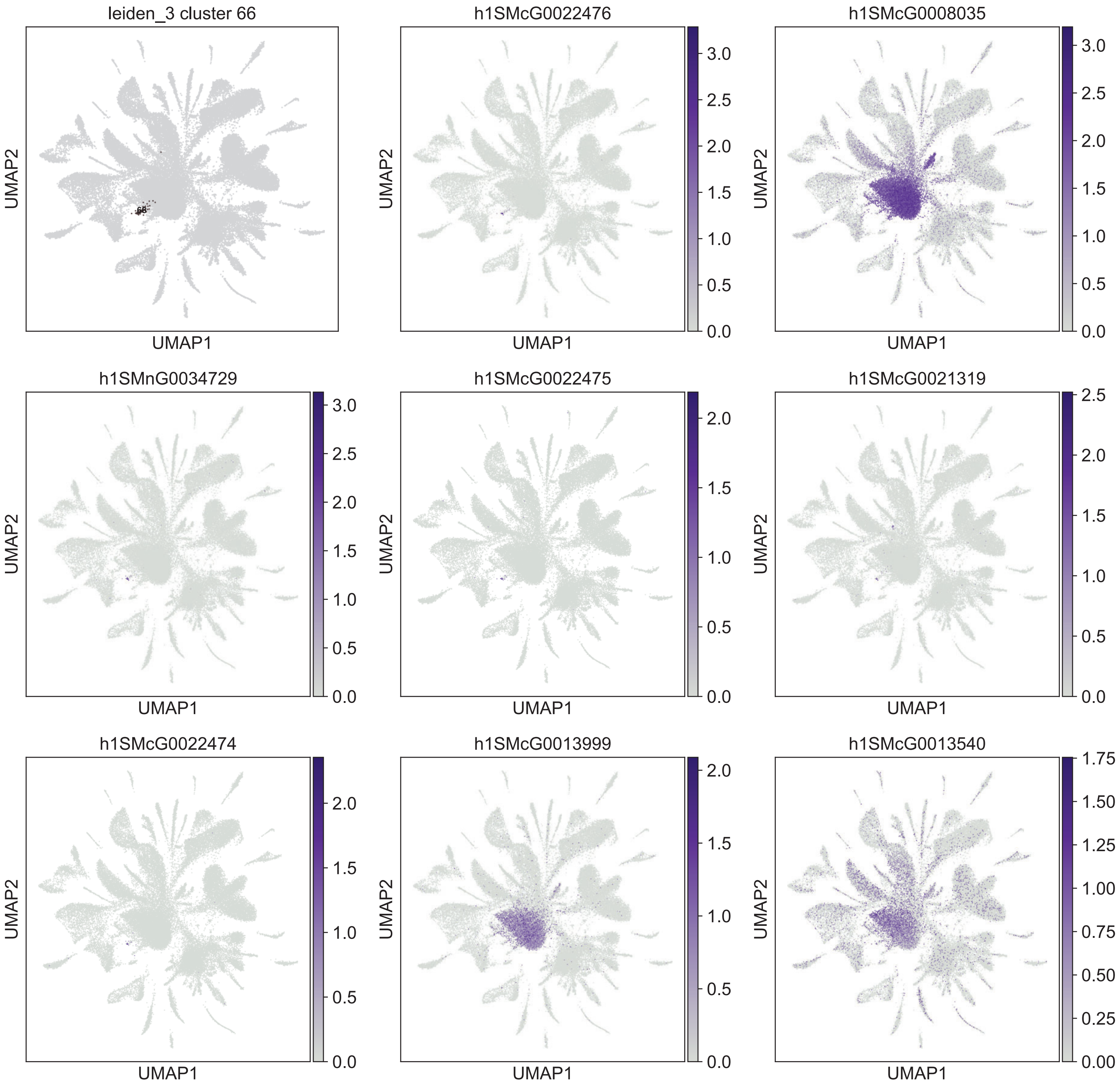

leiden_3 cluster 66
66
h1SMcG0022476
h1SMcG0008035
h1SMnG0034729
h1SMcG0022475
h1SMcG0021319
h1SMcG0022474
h1SMcG0013999
h1SMcG0013540
UMAP1
UMAP2
0.0
0.5
1.0
1.5
2.0
2.5
3.0
0.00
0.25
0.50
0.75
1.00
1.25
1.50
1.75

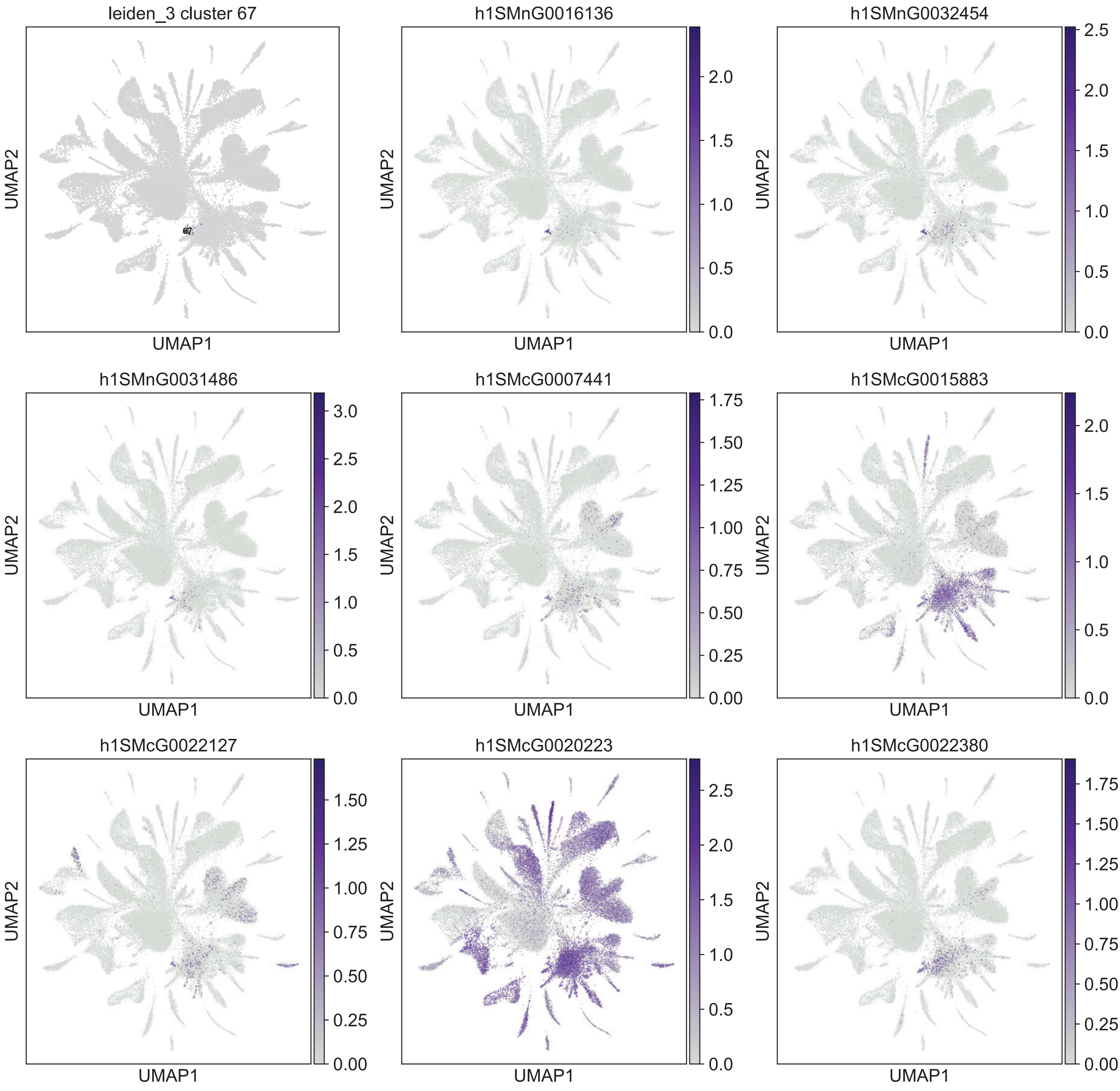
leiden_3 cluster 67
67
h1SMnG0016136
h1SMnG0032454
h1SMnG0031486
h1SMcG0007441
h1SMcG0015883
h1SMcG0022127
h1SMcG0020223
h1SMcG0022380
UMAP1
UMAP2

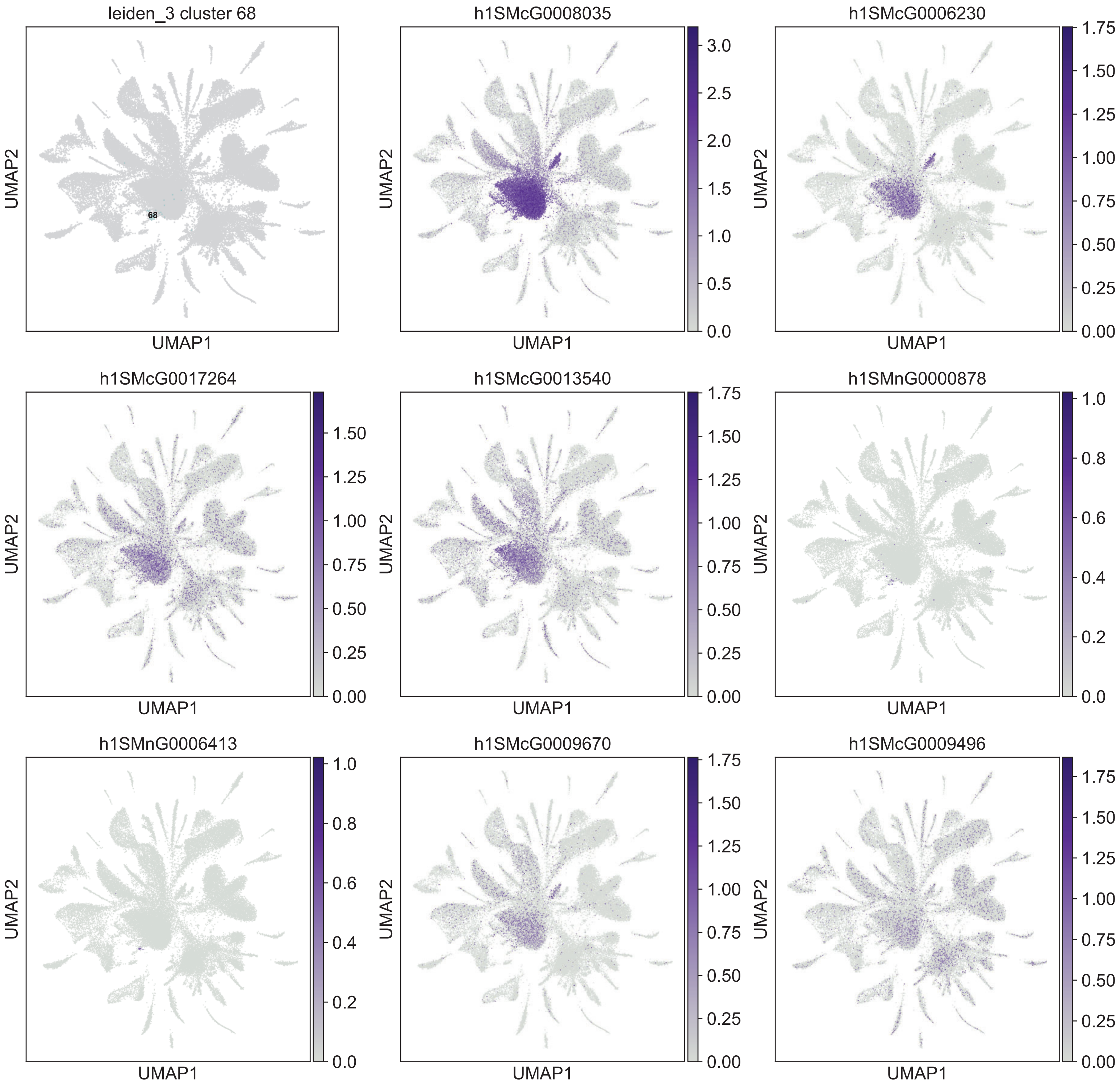
leiden_3 cluster 68
68
h1SMcG0008035
h1SMcG0006230
h1SMcG0017264
h1SMcG0013540
h1SMnG0000878
h1SMnG0006413
h1SMcG0009670
h1SMcG0009496
UMAP1
UMAP2

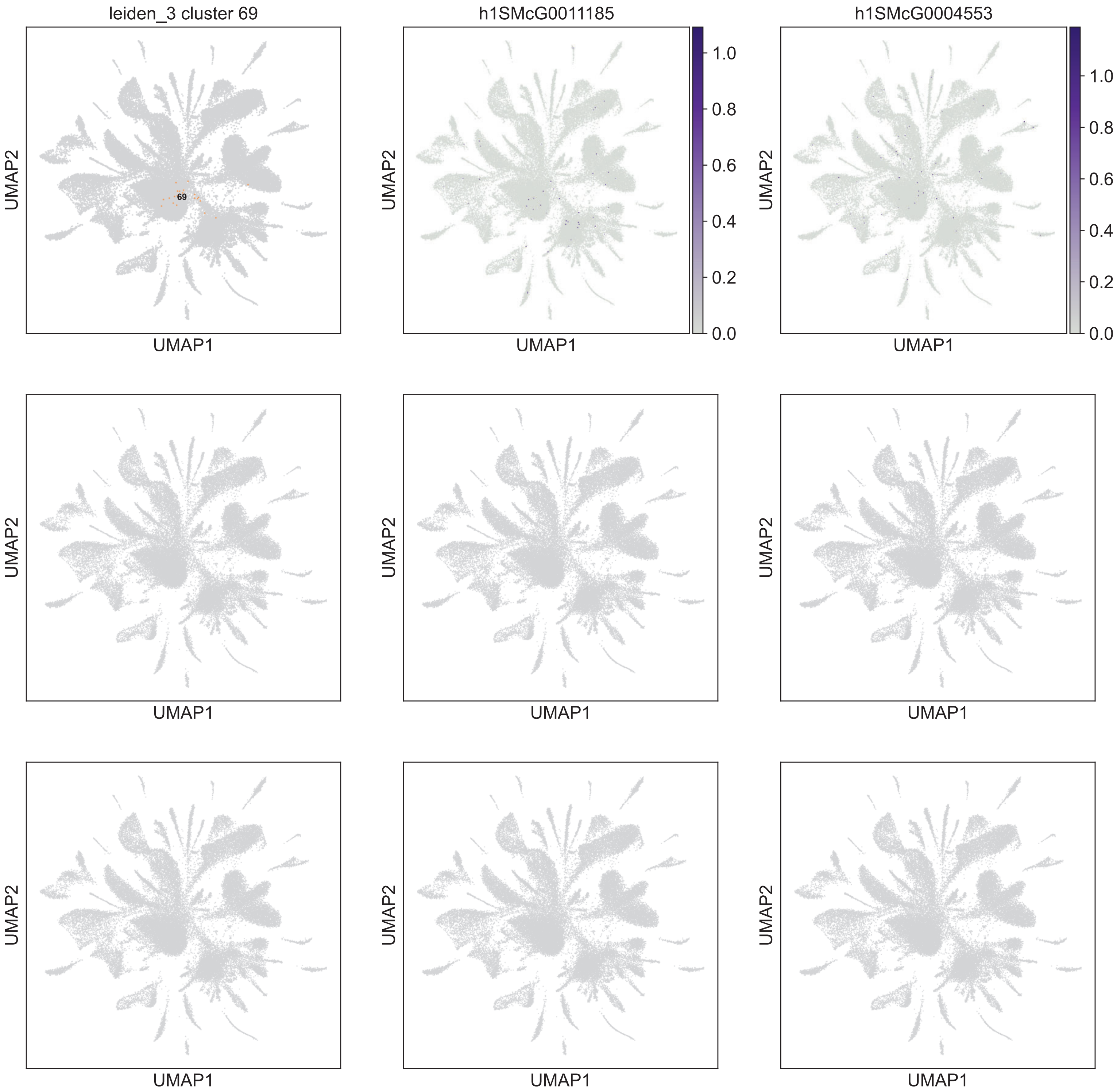
leiden_3 cluster 69
69
h1SMcG0011185
h1SMcG0004553
1.0
0.8
0.6
0.4
0.2
0.0
UMAP1
UMAP2

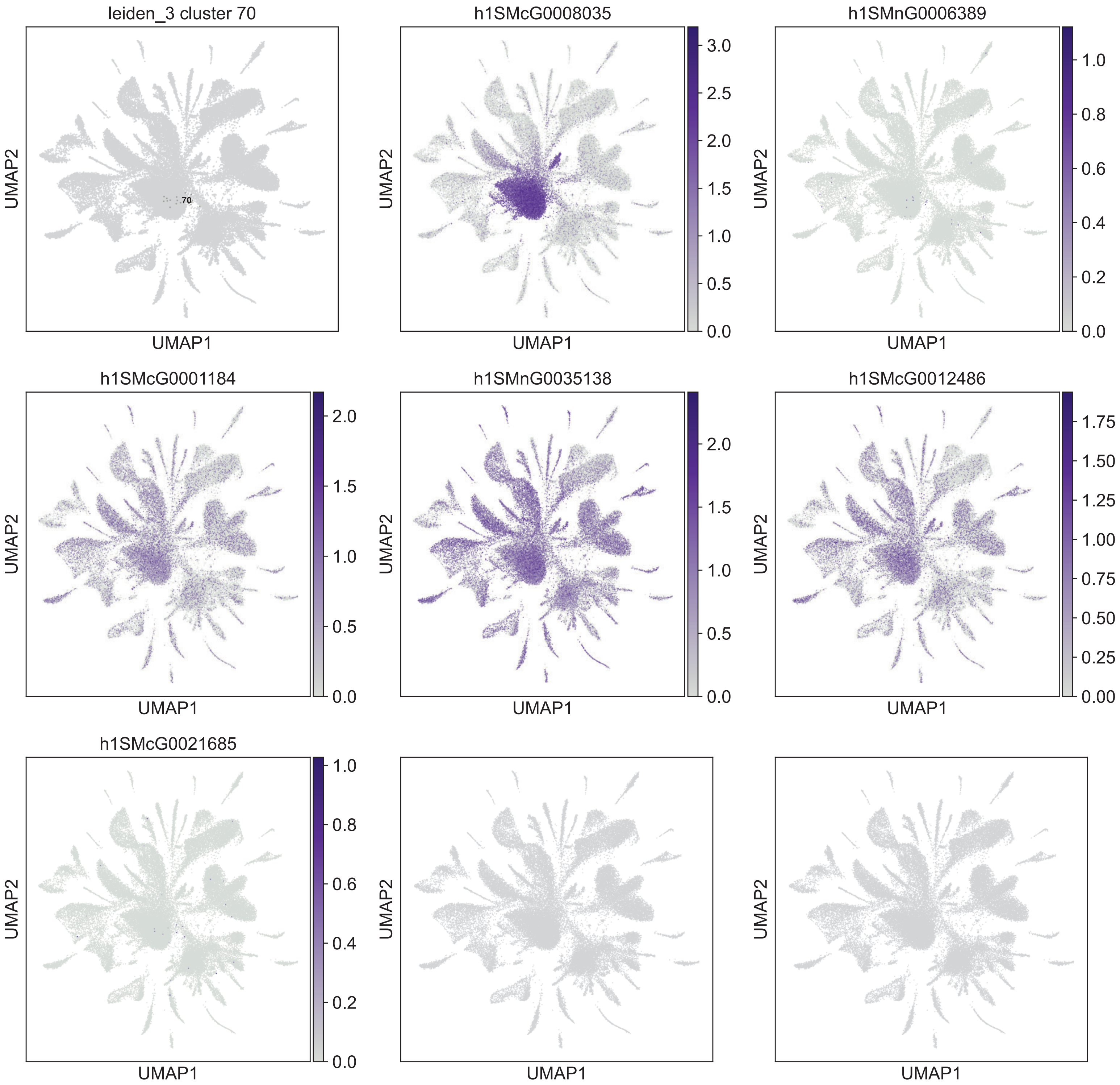
leiden_3 cluster 70
70
h1SMcG0008035
h1SMnG0006389
h1SMcG0001184
h1SMnG0035138
h1SMcG0012486
h1SMcG0021685
UMAP1
UMAP2
0.0
0.5
1.0
1.5
2.0
2.5
3.0
0.2
0.4
0.6
0.8
0.00
0.25
0.50
0.75
1.00
1.25
1.50
1.75

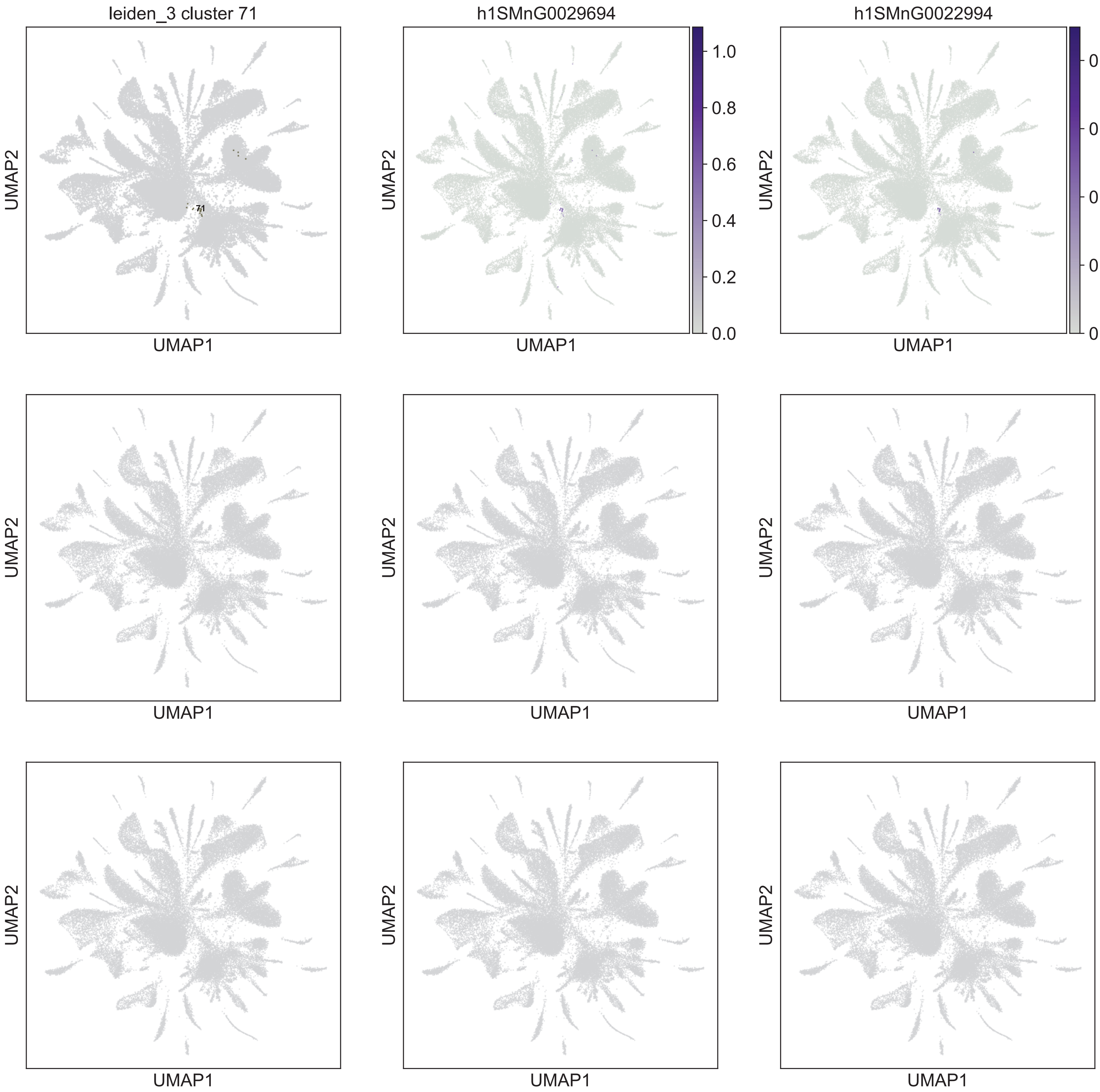

leiden_3 cluster 71
h1SMnG0029694
h1SMnG0022994
71
UMAP1
UMAP2
1.0
0.8
0.6
0.4
0.2
0.0

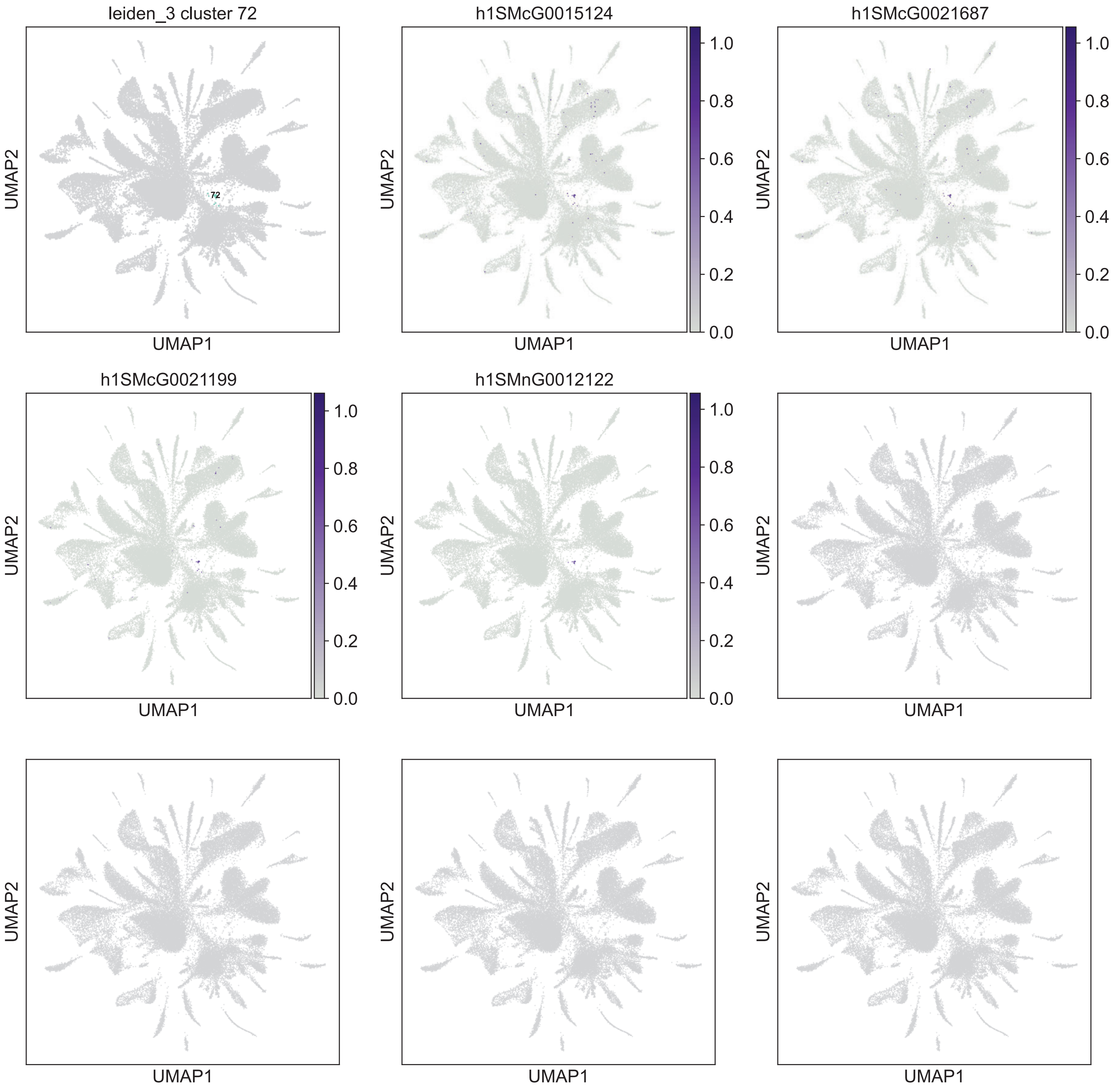

leiden_3 cluster 72
72
h1SMcG0015124
h1SMcG0021687
h1SMcG0021199
h1SMnG0012122
1.0
0.8
0.6
0.4
0.2
0.0
UMAP1
UMAP2

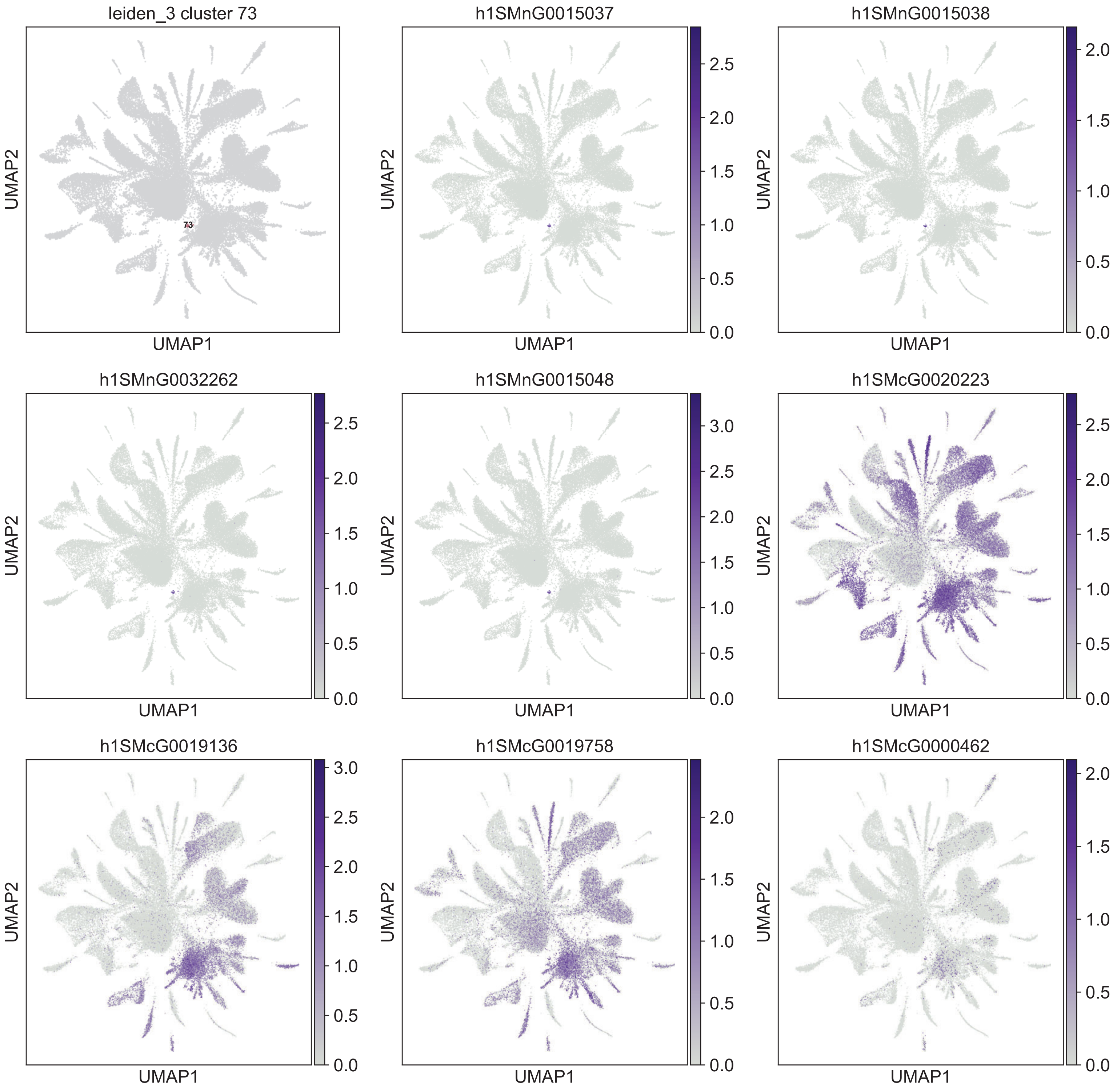
leiden_3 cluster 73
73
h1SMnG0015037
h1SMnG0015038
h1SMnG0032262
h1SMnG0015048
h1SMcG0020223
h1SMcG0019136
h1SMcG0019758
h1SMcG0000462
UMAP1
UMAP2
0.0
0.5
1.0
1.5
2.0
2.5
3.0
