## Supplementary figures and images for "Multiplex single-cell analysis of serotonergic neuron function in planarians reveals widespread effects in diverse cell types"

### Emili et al 2024 Serotonin Supplementary File 5 20240225 vs1.pdf

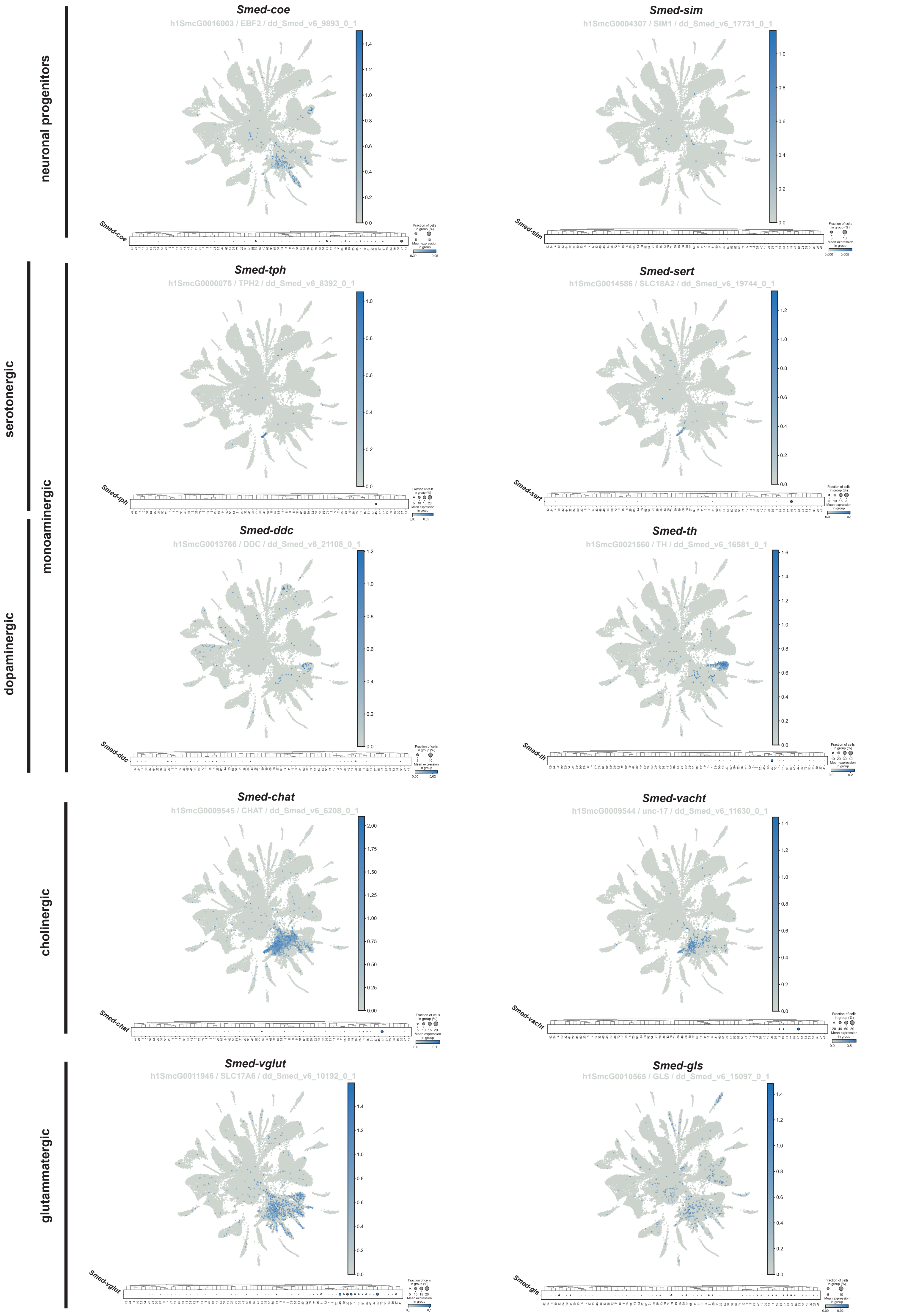

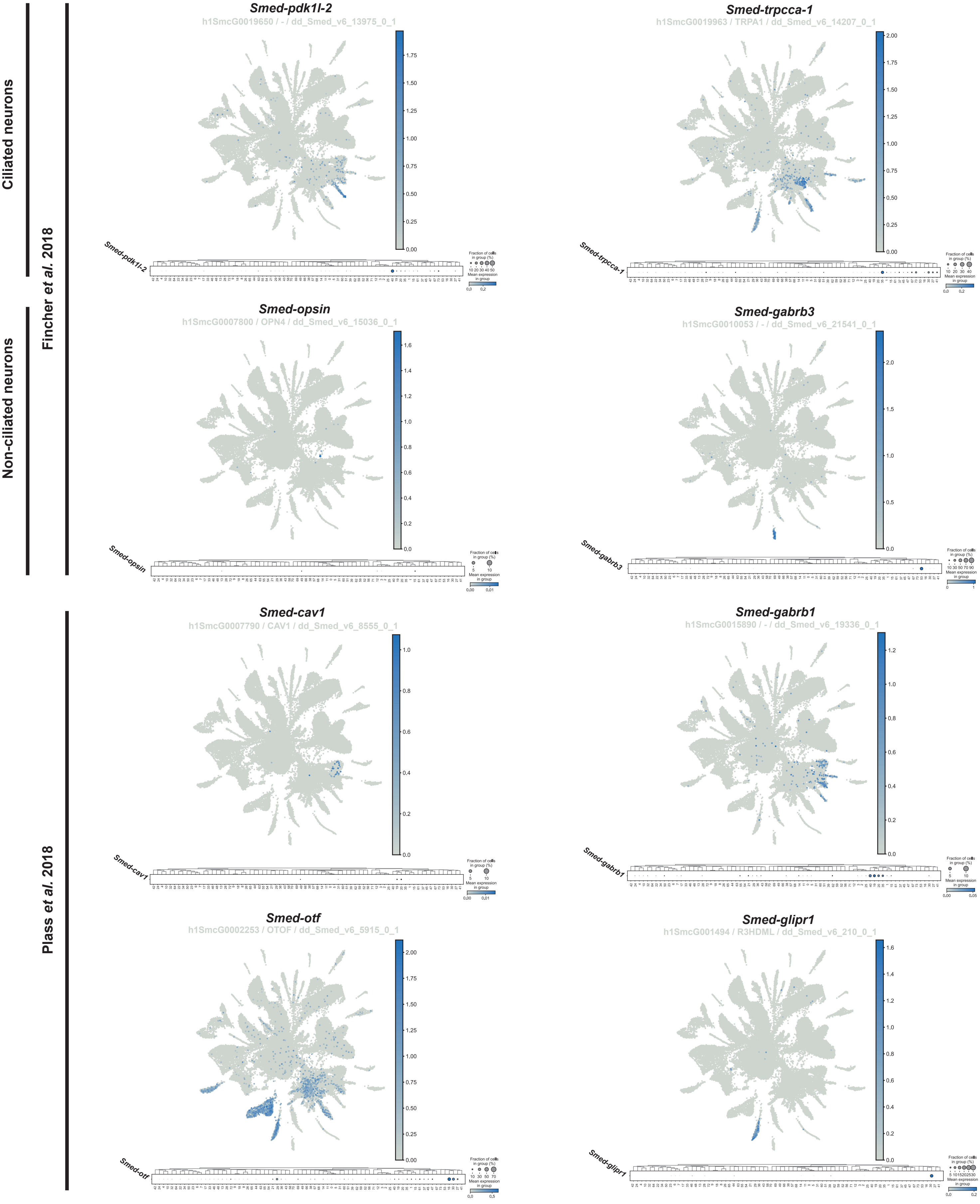

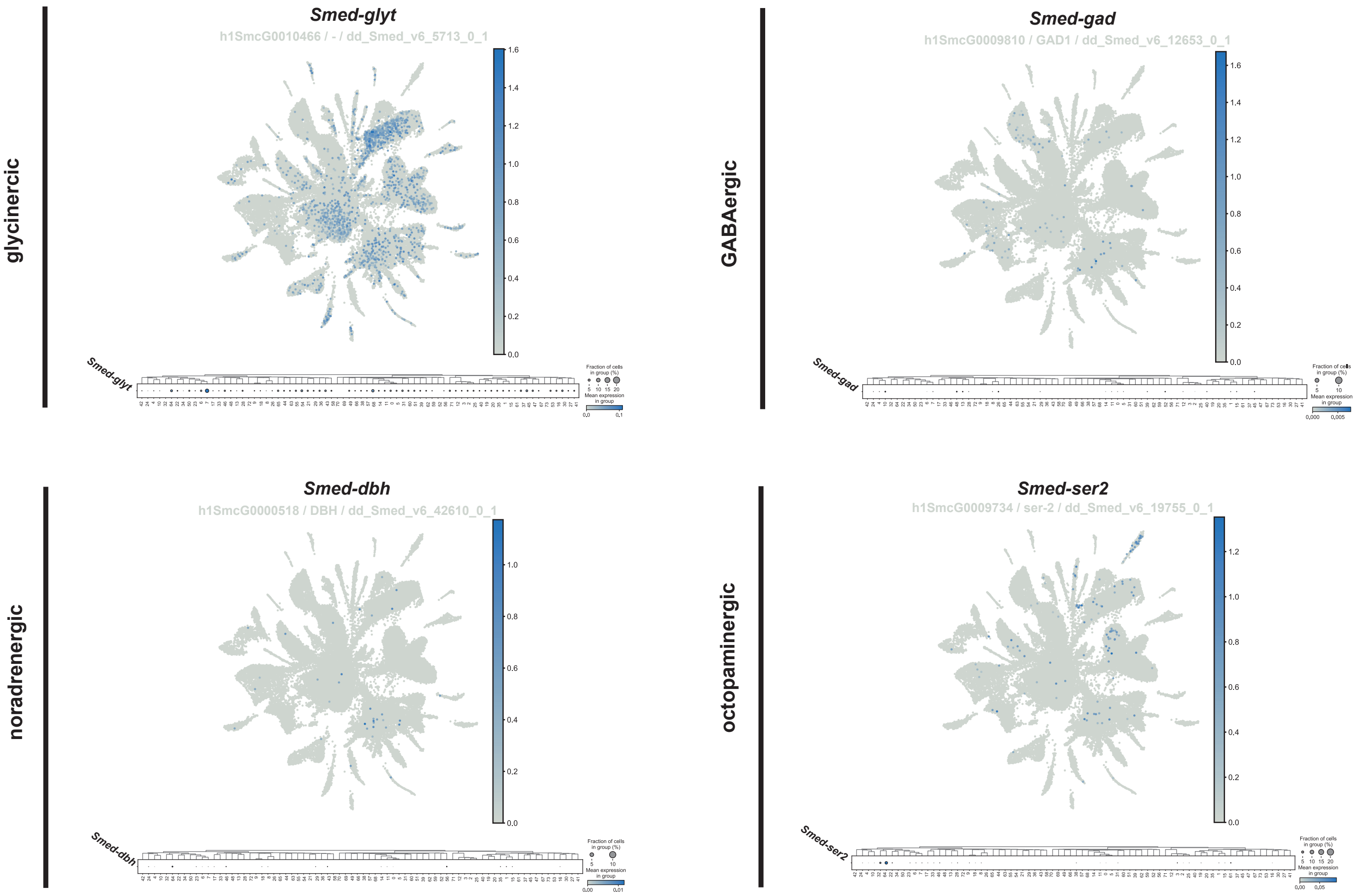

### Emili et al 2024 Serotonin Supplementary File 8 20240225 vs1.pdf

#### GO analysis for differentialy regulated genes

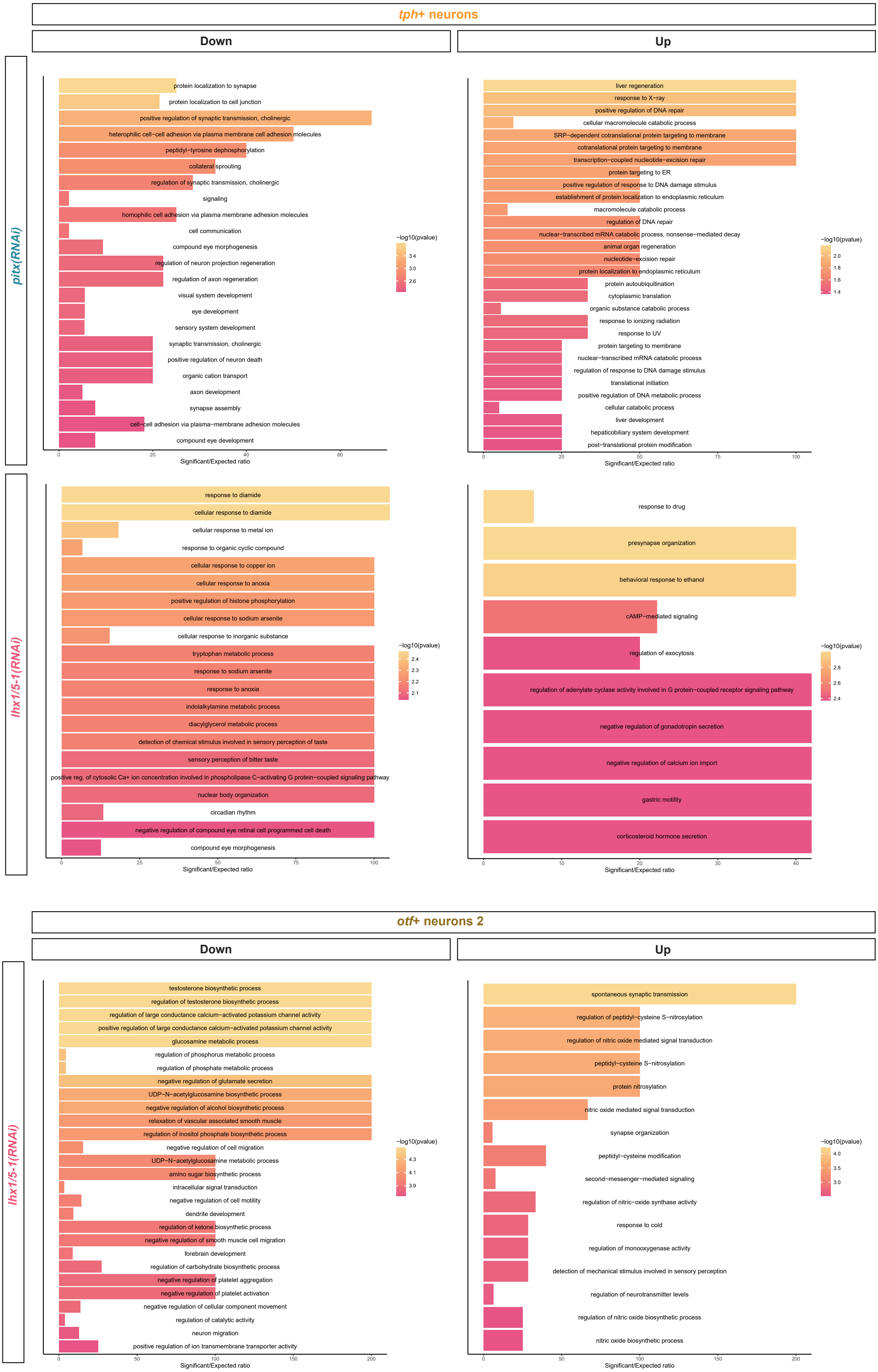

### Emili et al 2024 Serotonin Supplementary File 10 20240225 vs1.pdf

##### GO analysis for differentialy regulated genes

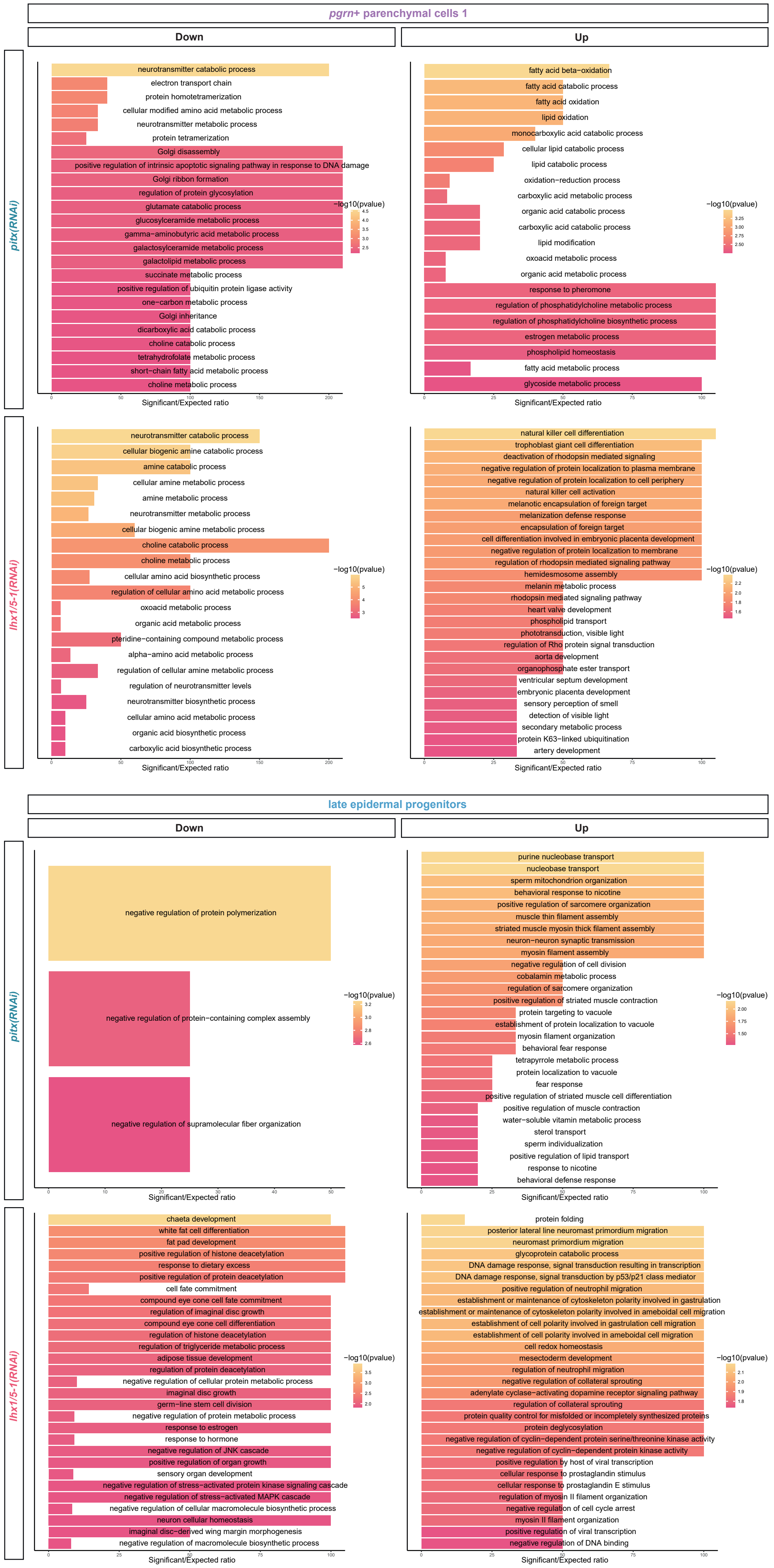

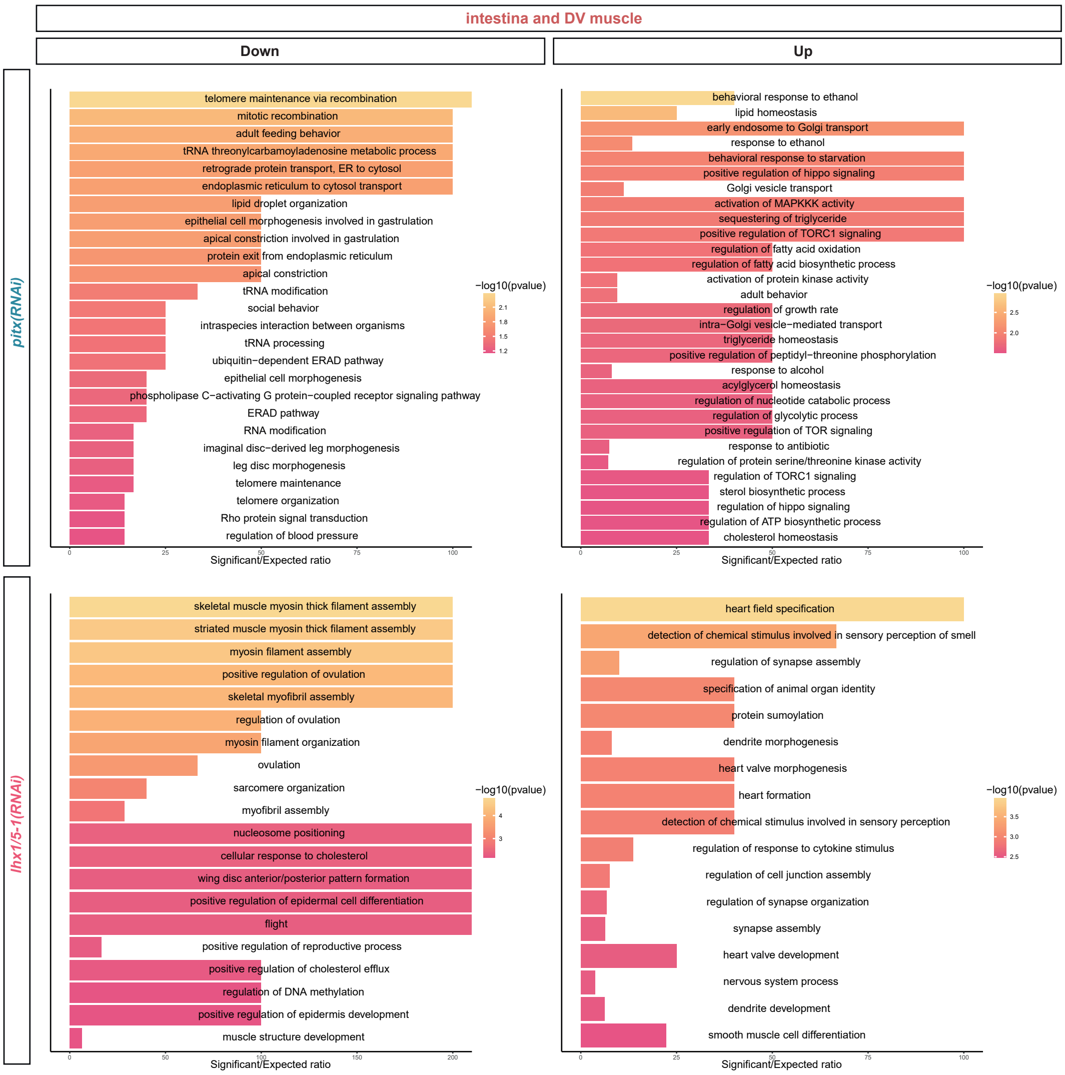
